# MethylBench: A comprehensive benchmark of DNA methylation profiling methods across diverse sequencing platforms

**DOI:** 10.64898/2026.04.28.721268

**Authors:** Lukas Laufer, Gilles Gasparoni, Thomas Hentrich, Linda Sofan, Jakob Admard, Elena Buena-Atienza, Michaela Pogoda, Stephan Ossowski, Nicolas Casadei, Olaf Rieß, Tobias B. Haack, Rebecca Buchert, Julia Schulze-Hentrich

## Abstract

**Background:** DNA methylation can be profiled using multiple technologies that vary in resolution, coverage and cost. Prior benchmarking efforts have laid important groundwork, yet critical gaps remain. The SEQC2 EpiQC study[14] provided a multi-platform QC framework across whole-genome bisulfite sequencing (WGBS), oxidative bisulfite sequencing, enzymatic deamination, ONT and Illumina 850k arrays, showing high overall concordance but limited replication and no coverage of current enzymatic conversion, hybridization-based panels or modern long-read chemistries. More recently, Sigurpalsdottir et al.[40] systematically compared methylation detection tools within the long-read domain, demonstrating high ONT accuracy relative to oxidative bisulfite sequencing, but restricted their analysis to ONT and PacBio, excluding short-read, targeted-panel and array-based platforms as well as the biological interpretability of differential methylation across platform classes. A comprehensive cross-platform benchmark addressing coverage heterogeneity and annotation redundancy in the interpretation of differentially methylated cytosines thus remains absent.

**Methods:** We compared six widely used technologies - Illumina EPIC array, TWIST, Whole-Genome Enzymatic Conversion (WGEC), Reduced Representation Bisulfite Sequencing (RRBS), Pacific Biosciences (PacBio) and Oxford Nanopore Technologies (ONT) - using GIAB reference samples and ten blood/fibroblast samples from 5 individuals. We assessed CpG coverage, consistency of differentially methylated cytosine (DMC) detection, and genomic annotation, with particular attention to overlapping signals across assays.

**Results:** Despite major assay differences, all technologies consistently identified DMCs enriched in promoter and intronic regions, marking these as robust hotspots of epigenetic variability. Annotation redundancy strongly shaped initial interpretations, with CpG island–related categories largely disappearing once annotations were collapsed to unique features. Sequencing-based methods (WGEC, TWIST, ONT) achieved the most comprehensive coverage, while EPIC arrays reliably captured promoter-associated differences despite limited scope. ONT showed strong concordance with short-read methods after coverage filtering, but required higher, more uniform coverage for reproducible CpG-level agreement. PacBio showed a coverage-independent discrepancy, with concordance plateauing regardless of mean coverage - pointing to residual technology-specific bias rather than a simple coverage effect.

**Conclusions:** Cross-platform benchmarking yields coherent biological insights once coverage and annotation redundancies are addressed. EPIC arrays remain valuable for promoter-focused cohort studies, WGEC and TWIST enable genome-wide discovery and ONT offers unique phasing and multimodal potential - together guiding method selection and more robust interpretation of DNA methylation data.

## Introduction

Epigenetic regulation of genomic information represents a fundamental mechanism of gene expression control and plays a crucial role in numerous biological processes as well as in the pathogenesis of various diseases, particularly cancer[9]. One of the most extensively studied epigenetic modifications is DNA methylation, in which a methyl group is added to the fifth carbon of the cytosine ring, forming 5-methylcytosine (5mC)[5]. This modification predominantly occurs at CpG dinucleotides and influences chromatin structure, gene silencing and cellular differentiation[27], with regulatory relevance that depends strongly on genomic context.

Against the backdrop of the growing relevance of epigenetic biomarkers in both research and clinical contexts, the accurate and cost-effective detection of 5mC has become increasingly important[4]. However, the diversity of available profiling technologies, differing in detection chemistry, genomic resolution, coverage breadth and cost, complicates the selection of appropriate methods for specific research or clinical questions and introduces the risk of platform-dependent analytical artifacts when results are compared across studies.

Prior benchmarking efforts have addressed parts of this challenge, but critical gaps remain. The SEQC2 EpiQC study[14] provided an influential multi-platform quality control framework, evaluating multiple whole-genome bisulfite sequencing library preparation protocols, oxidative bisulfite sequencing, enzymatic deamination sequencing, targeted methylation capture, single molecule ONT nanopore sequencing and Illumina 850k arrays across seven human cell lines[14]. While demonstrating overall high inter-assay concordance, EpiQC also identified meaningful differences in read mapping efficiency, CpG capture rates and platform-specific performance characteristics and explicitly noted that the limited number of replicates per protocol constrained the ability to distinguish assay-specific signal from technical noise[14]. Importantly, EpiQC was conducted prior to the widespread adoption of current whole-genome enzymatic conversion protocols(WGEC), hybridization-based targeted methylation panels, such as TWIST and modern long-read chemistries, including PacBio HiFi and ONT R10.4 and therefore does not reflect the methodological landscape, as it stands today. More recently, Sigurpalsdottir et al.[40] delivered a large-scale systematic comparison of methylation detection tools within the long-read domain, demonstrating high accuracy of ONT-based CpG methylation detection relative to oxidative bisulfite sequencing across 7,129 samples and introducing quality filters that substantially improve per-site reliability[40]. However, the scope of the study was explicitly restricted to ONT and PacBio and did not encompass cross-technology comparisons involving short-read sequencing, targeted enrichment panels or array-based profiling. Critically, neither study addressed the question of how annotation redundancy, the inflation of apparent platform-specific differences arising from overlapping genomic feature categories, confounds the biological interpretation of differentially methylated cytosines across platform classes, nor did they evaluate the impact of coverage heterogeneity in cross-platform DMC concordance in primary human tissue.

The MethylBench project was designed to address these open questions through a systematic comparison of six methodologically distinct platforms that together represent the current state of the field. Both technical and biological aspects, as well as economic considerations are taken into account in order to provide a robust and practically oriented framework for selecting suitable technologies for diverse applications in basic research, translational medicine and clinical diagnostics.

The project evaluates a range of established and emerging platforms for methylation analysis, which differ fundamentally in their underlying detection chemistry, sequencing modality and genomic coverage (Figure 1):

- **LR-GS: ONT** enables direct detection of 5mC by analyzing characteristic changes in electrical current as DNA strands pass through nanopores[7]. This technology offers long read lengths and low infrastructure costs, although methylation detection can be limited by bioinformatic uncertainty[7].
- **LR-GS: PacBio** employs Single-Molecule, Real-Time (SMRT) sequencing to detect epigenetic modifications such as 5mC through kinetic signatures of DNA polymerase activity[13]. While the method has been reported to provide high accuracy[13], our benchmark identifies residual, coverage-independent discordance with other platforms (see Results and Discussion).
- **Reduced Representation Bisulfite Sequencing (RRBS)** utilizes bisulfite conversion and enzymatic digestion to enrich for CpG-dense regions, thereby offering a cost-efficient approach focused on promoter regions[26]. However, it covers only a small fraction of the methylome[26].
- **Whole-Genome Enzymatic Conversion (WGEC)** is considered less damaging to the DNA than the historical gold standard using Whole-Genome Bisulfite Sequencing (WGBS)[32][23]. Both are constrained by high sequencing costs.
- The **Illumina EPIC BeadChip array** offers a cost-effective solution for large-scale cohort studies by enabling the quantitative analysis of over 850,000 CpG sites in the human genome. However, its coverage is limited and does not allow for the discovery of novel methylation sites.
- The **TWIST Methylation Panel** uses hybridization-based target enrichment and allows for flexible panel design[15]. In combination with high-throughput sequencing, it offers a balanced trade-off between cost, depth and genome-wide coverage, though with limited sensitivity to broader epigenetic contexts[15].

**Figure 1:**
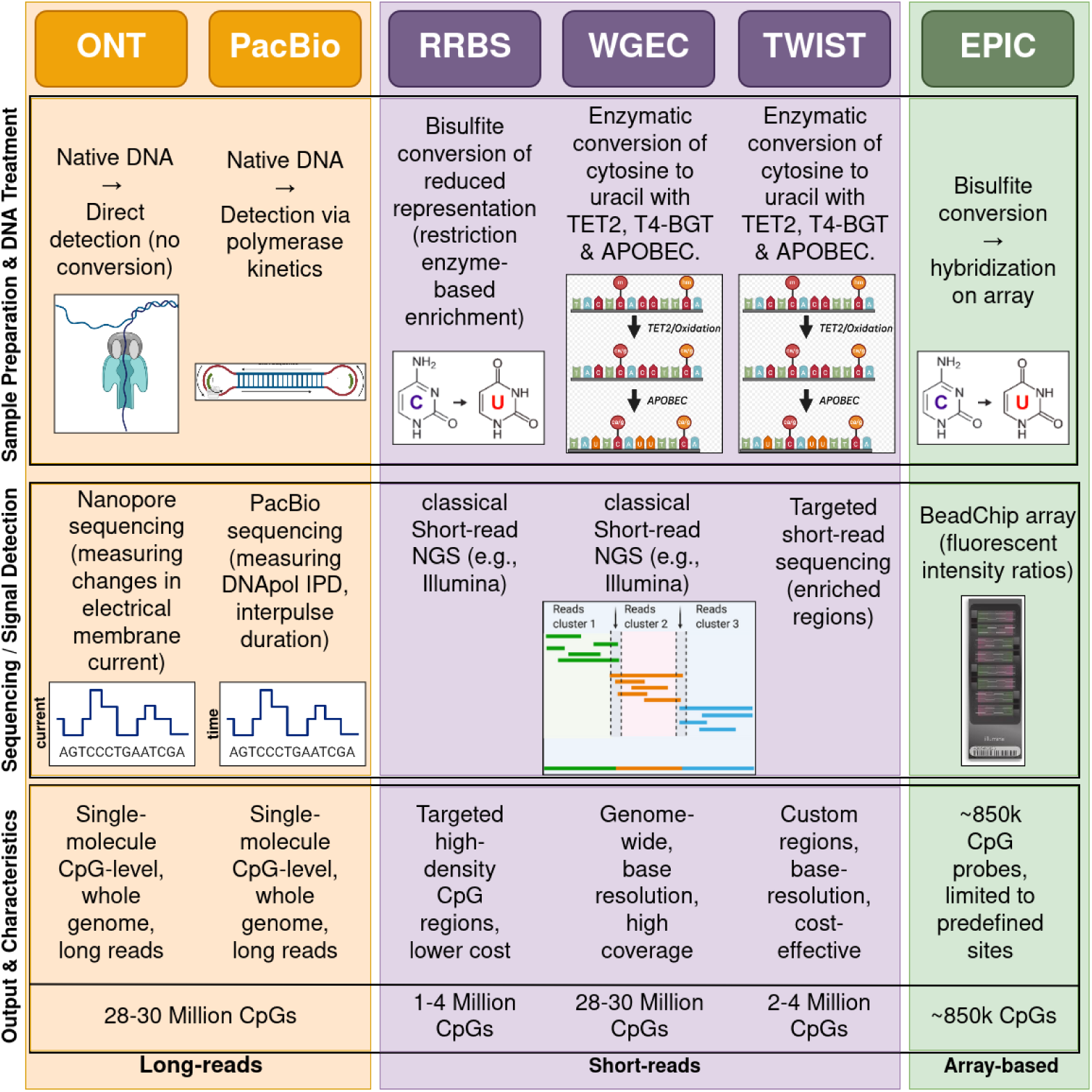
Overview of DNA methylation detection methods and their characteristics. This table compares six widely used genomic and epigenomic technologies in terms of their ability to detect 5-methylcytosine (5mC), underlying detection principles and genome-wide coverage. Methods such as Oxford Nanopore and PacBio allow for direct or indirect detection of 5mC without chemical treatment, while conversion-based methods (e.g., RRBS, WGEC, TWIST and EPIC arrays) rely on chemical distinction of methylated and unmethylated cytosines. Targeted approaches like TWIST panels offer high specificity but limited genome-wide coverage. Long-read sequencing technologies enable the direct detection of multiple DNA base modifications, including both 5mC and 5-hydroxymethylcytosine (5hmC), without the need for chemical conversion. For the GIAB2 (HG002) PacBio dataset used in this study, on-instrument base-modification calling additionally provided 5hmC and N6-methyladenine (6mA) calls alongside 5mC; these additional modification channels were not analyzed here but illustrate the expanding multimodal capability of current long-read chemistries (see Methods). In contrast, short-read sequencing approaches vary in their capacity to resolve cytosine modifications: whole-genome enzymatic conversion (WGEC), including targeted implementations such as the TWIST panel, allows discrimination of 5hmC from 5mC through selective enzymatic processing.[32]

MethylBench conducts a comparative analysis of these technologies with respect to sensitivity, specificity, reproducibility, data complexity and economic factors such as scalability, cost per sample and infrastructure requirements. The overarching goal is to develop a nuanced assessment framework to inform the selection of epigenetic methods for applications in basic research, translational medicine and clinical diagnostics.

To facilitate the interpretation of methylation data in functional and regulatory contexts, cytosine-phosphate-guanine (CpG) sites are commonly annotated with respect to both their spatial relation to CpG islands (CGIs) and gene structure. Figure 2 illustrates the canonical classification of CpG regions into CpG islands, shores (up to 2 kb flanking the CGI), shelves (2–4 kb from the CGI) and open sea regions, which reside distal to CGIs. These categories have been shown to exhibit distinct methylation dynamics and regulatory potential, with shores in particular being enriched for tissue-specific methylation changes.

**Figure 2:**
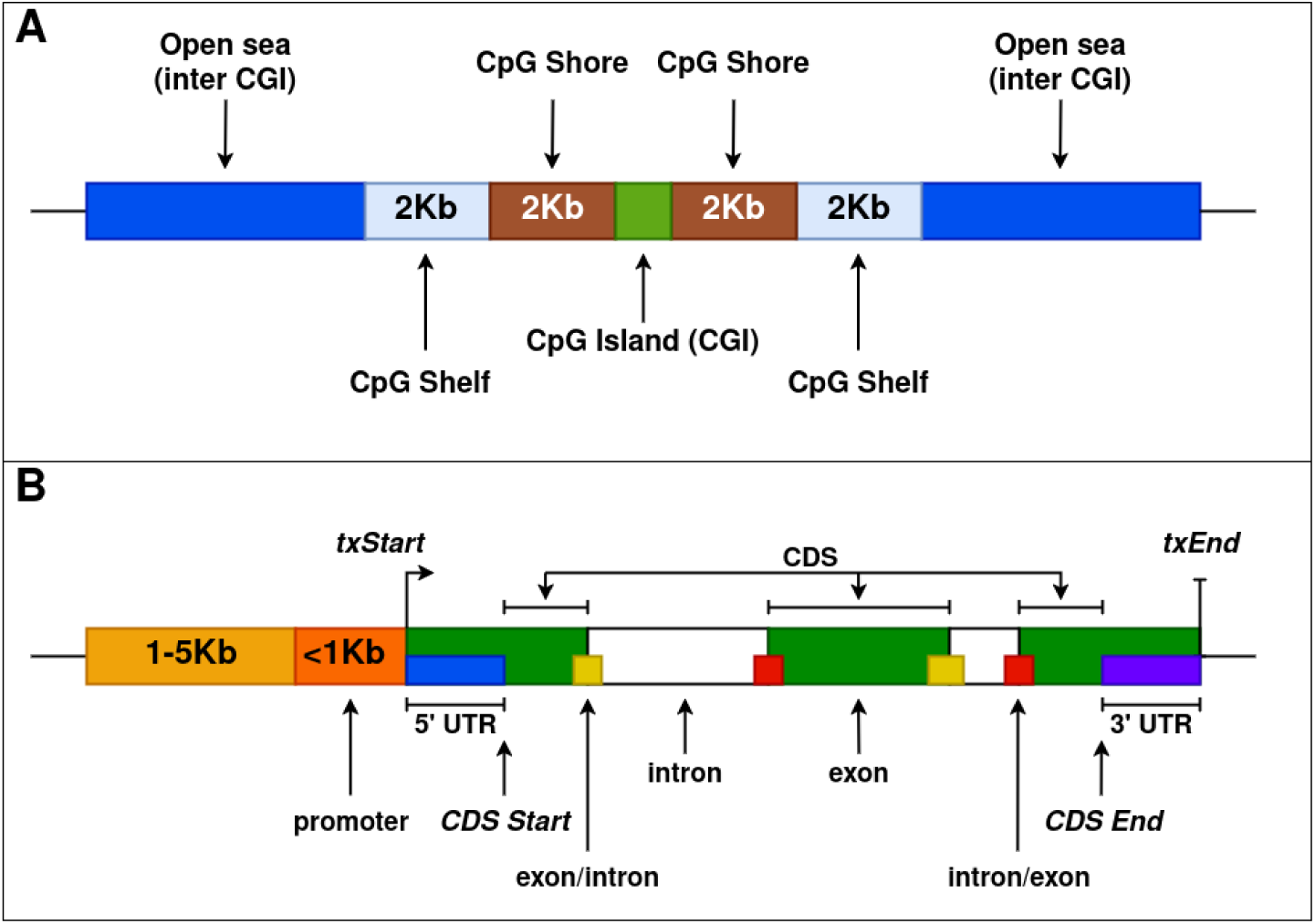
Classification of CpG Sites Based on Proximity to CpG Islands & Genomic Contextual Annotation of CpG Sites Relative to Gene Structure. (A): Illustration of CpG islands (CGIs), shores, shelves and open sea regions along the genome, highlighting their relative distances and functional annotation contexts. **(B):** Schematic representation of gene elements including promoters, untranslated regions (UTRs), exons, introns, exon/intron boundaries, intron/exon boundaries and coding sequences (CDS), emphasizing their utility in CpG site annotation. Additionally the transcription start and end sites (txStart & txEnd respectively) are shown.

Complementing this, Figure 2 provides an overview of commonly used genomic annotations in the context of DNA methylation and gene regulation analyses. Panel A illustrates a CpG-centric annotation model, where genomic regions are categorized based on their proximity to CpG islands (CGIs). CGIs (green) are typically associated with regulatory elements such as promoters and are flanked by CpG shores (brown; up to 2 kb from the CGI) and CpG shelves (light blue; 2–4 kb from the CGI). Regions beyond these are classified as open sea (dark blue), often representing intergenic or less CpG-dense regions.This annotation schema provides a granular understanding of how methylation events map onto functionally distinct genomic segments. Promoter-associated methylation, especially within 1 kb upstream of transcription start sites (TSS), is typically associated with transcriptional repression, while gene body methylation may correlate positively with transcriptional activity in certain contexts.

Panel B depicts a gene-centric annotation scheme, showing key gene features relative to the transcription start site (txStart) and end (txEnd). This includes upstream promoter regions (1-5 kb, orange; *<*1 kb, red), 5’ and 3’ untranslated regions (UTRs), coding sequence (CDS), exons (colored segments) and introns (white). This hierarchical structure enables the functional categorization of genomic elements, supporting the interpretation of regulatory mechanisms, especially in epigenomic studies. The concept of this figure was taken from the annotatr package[6].

Beyond promoters and gene bodies, DNA methylation plays a central role at distal regulatory elements and repetitive sequences, with important functional consequences that are best captured by whole-genome assays. Several WGEC-based studies and methylome atlases have shown that dynamic CpG methylation is enriched at distal regulatory regions and enhancers and that changes at these sites strongly associate with cell-type specific regulatory programs[42][47][24]. In parallel, DNA methylation constitutes a principal mechanism for repressing transposable elements (TEs) and maintaining heterochromatin stability; loss or redistribution of methylation at repetitive elements can lead to TE reactivation and genome instability[10][17]. These roles underline the distinct advantage of genome-wide, base-resolution methods (WGEC, long-read methylation calling) over targeted approaches when the aim is to interrogate enhancer activity, repetitive regions, or long-range regulatory interactions that are poorly sampled by probe-based assays.

Aberrant DNA methylation underlies multiple classes of human disease. Classic imprinting disorders illustrate how localized methylation changes at imprinting control regions produce profound developmental phenotypes[11]. Repeat-expansion diseases provide a clear mechanistic example in which expansion of a trinucleotide repeat leads to regional hypermethylation and stable gene silencing[31][37]. In oncology, large-scale reprogramming of the methylome - global hypomethylation of repeats coupled with focal hypermethylation of CpG islands and promoters - represents a hallmark of many cancers and has functional consequences for genome stability and tumor suppressor silencing[3][41]. Collectively, these disease-associated patterns argue for using comprehensive methylome assays when the biological question involves imprinting, repeat biology, or widespread methylation remodeling-situations in which targeted panels or arrays may miss diagnostically or mechanistically important signals.

## Results

### Interpretation of quality control metrics reveals method-specific strengths and limitations in DNA methylation sequencing

We systematically compared five DNA methylation profiling technologies-ONT, PacBio, RRBS, WGEC and TWIST-across blood, fibroblast and GIAB reference samples (Figure 3). The analysis highlights clear trade-offs between short- and long-read platforms.

**Figure 3:**
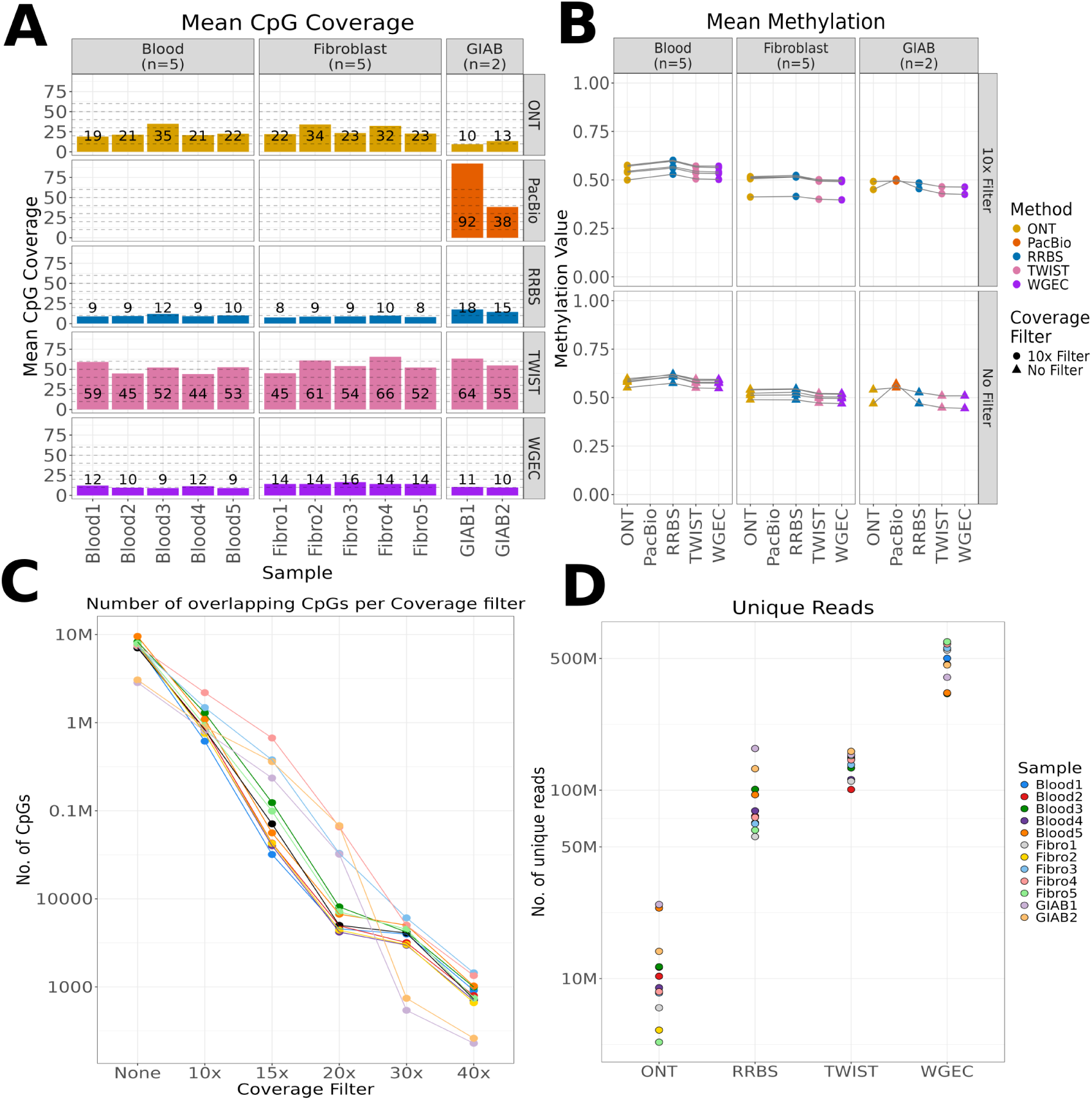
Long-Read vs. Short-Read DNA methylation profiling reveals trade-offs between coverage depth, CpG retention and read characteristics. This figure summarizes key quality control and performance metrics across five DNA methylation profiling techniques (ONT, PacBio, RRBS, TWIST, WGEC) applied to blood, fibroblast and GIAB samples. **(A):** Barplots showing the mean CpG coverage per sample and method; dashed lines indicate multiples of a 10× coverage threshold & numbers represent the rounded mean CpG coverage. **(B):** Mean methylation levels per sample and method, shown both unfiltered and with a 10× coverage cutoff, revealing reduced variance across platforms at higher coverage. In the GIAB cohort, the deviation observed for one PacBio sample does not track with sequencing coverage - the sample with markedly higher coverage did not show closer agreement with the other platforms - reflecting a technology-specific property of PacBio methylation calling rather than insufficient CpG-level sampling. **(C):** Log-scale line plot of the number of overlapping CpG sites retained at increasing coverage thresholds, indicating a rapid decline in shared CpGs with stricter filters. **(D):** Dot plot comparing the number of unique reads obtained per sample-method combination, with long-read methods showing fewer but larger alignments.

Short-read methods (RRBS, TWIST, WGEC) generally achieved a more uniform CpG coverage, making them well suited for large-scale comparative studies. In contrast, long-read platforms (ONT, PacBio) provided lower CpG coverage but offered distinct advantages in phasing[25][38] and long-range methylation context. Notably, some coverage variability was observed within platforms. For instance the CpG coverage was considerably deviating from the mean (genomic) coverage in ONT datasets for Blood5 and Fibro2. The mean genome coverage for ONT datasets was even across the samples (Figure 3A/Supplementary Figures S10 & S11). Lower sample purity, differences in read length or limitations in accuracy might influence the capture of methylation signals at the selected CpG sites. Average methylation levels across all analyzed samples (blood, fibroblasts and Genome in a Bottle references) clustered around 50–60%, largely consistent across methods. Applying a 10x coverage filter substantially reduced inter-platform variance-particularly between ONT and PacBio (Figure 3B). These observations align well with published human methylomes: whole-genome bisulfite sequencing (WGBS) of human peripheral blood mononuclear cells (PBMCs) reported a global CpG methylation level of ca. 68%[23]. Although this reference value was obtained using bisulfite rather than enzymatic conversion, WGBS and WGEC are considered methodologically comparable with respect to global CpG methylation estimates, as both rely on selective conversion of unmethylated cytosines and have been shown to yield concordant genome-wide methylation levels[32], while large-scale reference maps of human fibroblasts demonstrated mean methylation levels around 50–70% depending on genomic context[24][16]. Thus, the 50–60% averages observed here fall within the expected range for human blood and fibroblast methylomes, reinforcing that our data accurately recapitulate known epigenetic baselines while also highlighting the benefit of coverage thresholds for harmonizing platform-specific measurements. An exception to this overall consistency was observed in the GIAB reference samples, where one PacBio dataset showed a pronounced deviation in mean methylation relative to the other platforms (Figure 3B). This sample was sequenced at the highest average coverage among all GIAB replicates (92x), compared to 38x for the second PacBio GIAB sample. Notably, the deviation was not resolved, and if anything more pronounced, in the higher-coverage sample, and persisted even when applying a 10x CpG coverage filter. This indicates that the observed shift in mean methylation estimates is not explained by insufficient molecule sampling but instead reflects a technology-specific measurement property of PacBio polymerase kinetics–based methylation calling that is independent of, and not resolved by, sequencing depth - consistent with the coverage-independent discordance described in detail below (Figure 4).

**Figure 4:**
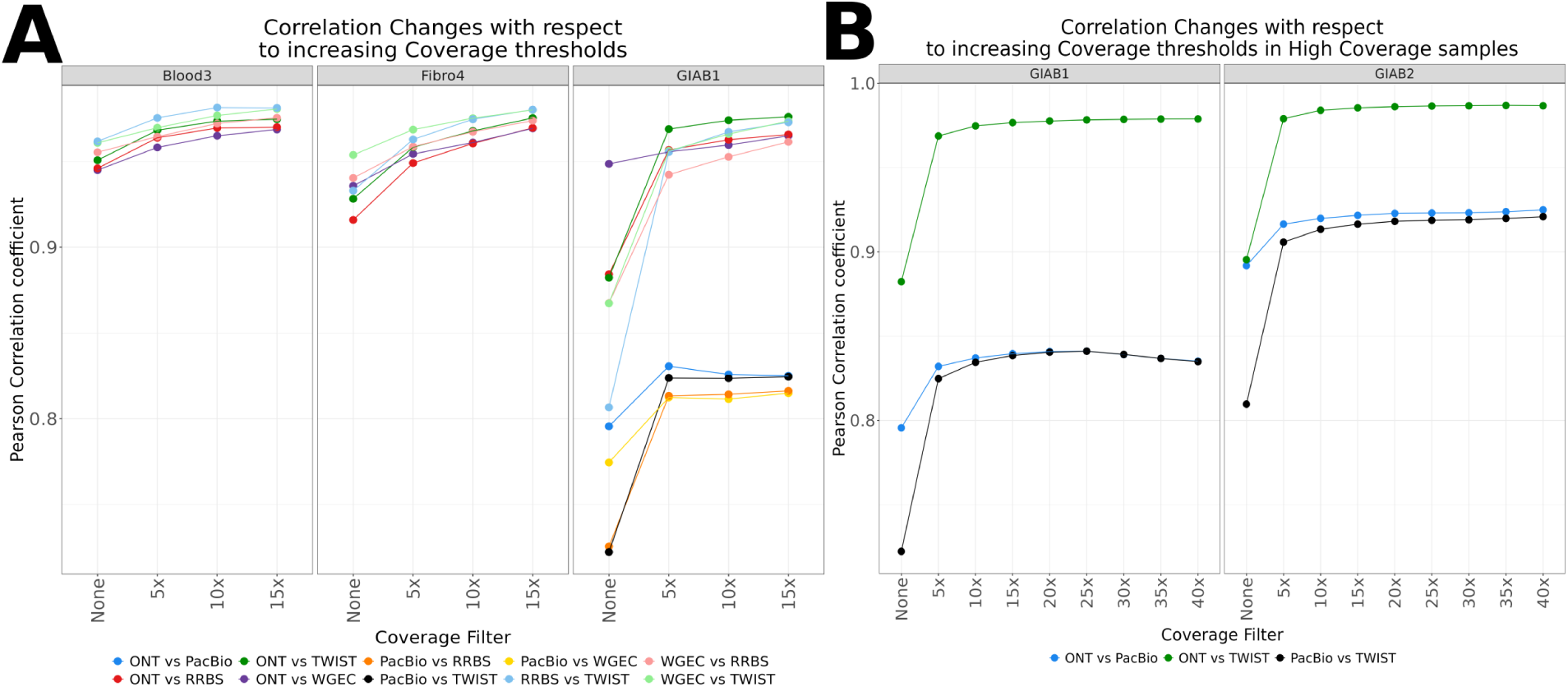
Increasing coverage thresholds improve cross-platform methylation concordance. (A): Pearson correlation coefficients of CpG methylation levels between all pairwise combinations of five methylation profiling methods (ONT, RRBS, TWIST, WGEC and PacBio) across different samples. PacBio data was only available for the GIAB sample. Correlations were computed at increasing minimum coverage thresholds (x-axis) and are shown separately for each sample. A consistent increase in inter-method correlation is observed with rising coverage cutoffs, particularly in comparisons involving ONT and RRBS, indicating that higher coverage thresholds reduce noise and improve measurement agreement between technologies. In the GIAB1 sample, correlations involving PacBio plateau despite increasing coverage thresholds, indicating that residual inter-platform differences persist even at high PacBio coverage (92x; Figure 3A) and are not primarily driven by low-coverage CpGs. **(B):** We restricted this direct comparison to ONT, TWIST & PacBio, as these represent the highest coverage methods among the platforms assessed. Applying progressively stricter coverage filters revealed a rapid increase in correlation up to approximately 5–15x, beyond which correlation plateaued at *r* = 0.98–0.99 for ONT vs. TWIST. Agreement involving PacBio was consistently lower, plateauing at *r ≈* 0.83–0.84 for GIAB1 and *r ≈* 0.92 for GIAB2.

The number of shared CpGs declined sharply with stricter coverage cutoffs, disproportionately affecting long-read data (Figure 3C). Supplementary Figure S2 confirms this trend across blood, fibroblast and GIAB samples, showing steadily increasing cross-platform correlations with higher thresholds. Based on this combined evidence, we selected a 10x cutoff as an optimal balance, retaining approximately one million CpGs for downstream analyses.

Finally, read-level comparisons revealed that short-read methods yielded many more unique alignments per sample, whereas ONT and PacBio produced fewer but substantially longer reads (Figure 3D). This trade-off in distinct read length distributions across technologies is illustrated in Supplementary Figure S5.

Taken together, these results emphasize a central trade-off: short-read methods maximize CpG retention and cross-platform comparability, while long-read sequencing provides complementary strengths in phasing[25][38] and structural methylation analysis, albeit at the cost of reduced coverage and reproducibility.

### Coverage filtering improves concordance between methylation profiling methods

To quantify the effect of coverage thresholds on cross-platform agreement, we computed pairwise Pearson correlations of CpG methylation values across ONT, RRBS, TWIST and WGEC in blood and fibroblast samples, with additional PacBio data for the GIAB samples (Figure 4A). Correlations consistently improved with increasing minimum coverage, confirming that low-coverage CpGs contribute substantial noise. The effect was most pronounced in comparisons involving ONT and RRBS, which started below r = 0.92 at unfiltered levels but improved markedly at 10–15x coverage.

In contrast, WGEC and TWIST, which provided more even and deeper coverage, showed high baseline concordance and smaller relative gains with filtering. Supplementary Figure S2 extends these observations to blood and GIAB samples, demonstrating that the positive effect of coverage thresholds on correlation is consistent across tissues. Collectively, these results establish coverage filtering as an essential step for reliable integration of methylation data across platforms.

Notably, in the GIAB1 sample - where PacBio achieved the highest mean coverage observed among all GIAB replicates (92x; Figure 3A) - correlations between PacBio and the other platforms nonetheless plateaued at the lowest level observed in this study (*r ≈* 0.83–0.84; Figure 4B), with little to no further improvement beyond a 5x threshold. This pattern is not explained by insufficient PacBio coverage: in GIAB2, where PacBio coverage was substantially lower (38x), agreement between PacBio and the other platforms was higher (*r ≈* 0.92) than in the higher-coverage GIAB1 sample. This indicates that cross-platform disagreement for PacBio is not simply a function of sequencing depth and that in this dataset higher raw coverage did not predict better cross-platform agreement - reinforcing that residual discrepancies reflect technology-specific differences in methylation signal interpretation rather than coverage-related noise. As a result, increasing coverage thresholds alone is insufficient to further harmonize PacBio methylation estimates with other methods in this context. Notably, GIAB1 and GIAB2 PacBio data were generated using different chemistry and instrument software generations (standard Revio SPRQ chemistry and Platinum Pedigree processing for GIAB1, versus SPRQ-Nx multi-use chemistry with an updated basecalling model for GIAB2; see Methods), a factor considered further in the Discussion.

In contrast, short-read methods such as WGEC and TWIST, characterized by deeper and more uniform coverage, showed high baseline concordance and smaller relative gains with additional filtering. These findings highlight that platform-specific coverage depth strongly influences apparent methylation agreement, particularly for long-read technologies.

For the GIAB reference samples, we further assessed the relationship between coverage and concordance at higher stringency, directly comparing ONT and TWIST (Figure 4B). Applying progressively stricter coverage filters revealed a rapid increase in correlation up to around 15–20x, after which values plateaued near *r* = 0.98 *−* 0.99. This saturation effect indicates that beyond a certain threshold, additional sequencing depth yields diminishing returns, as remaining differences largely reflect systematic platform biases rather than stochastic sampling noise. These findings indicate that applying moderate coverage thresholds (10–15×) optimizes the trade-off between retaining sufficient data and ensuring reproducibility across platforms, whereas setting excessively high thresholds provides little additional improvement in methylation concordance analyses.

### Coverage threshold improves methylation signal consistency across technologies

To assess how different sequencing methods and coverage thresholds shape the distribution of DNA methylation values across samples, we visualized the methylation beta value densities using ridge plots stratified by method, sample and coverage (Figure 5A). Each row corresponds to a distinct biological sample, with curves colored by sequencing method. Black-outlined curves represent unfiltered data (No coverage filter), while red-outlined curves reflect a stringent 10× coverage filter.

**Figure 5:**
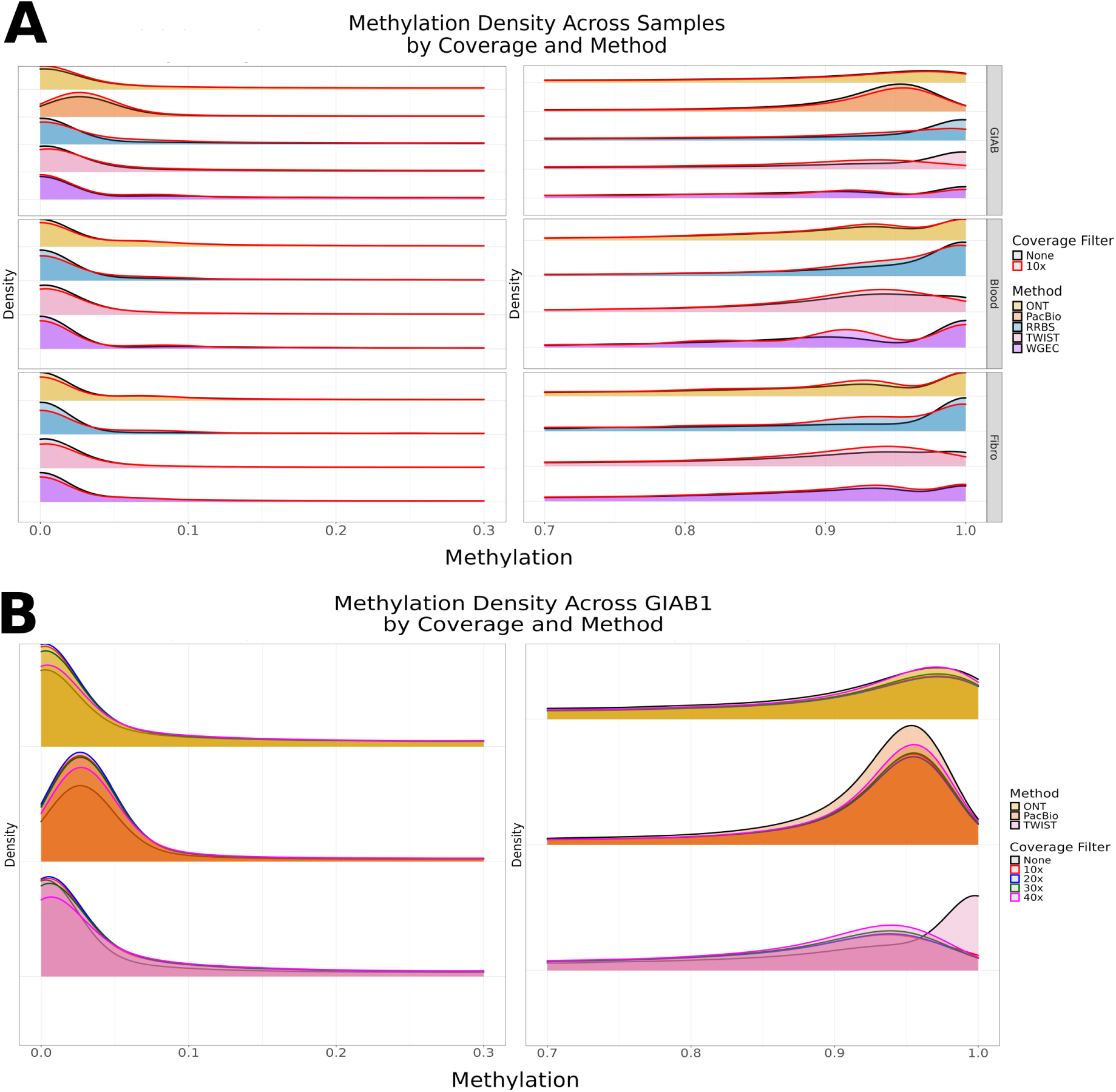
Impact of coverage filtering on methylation density distributions across methods and samples. (A): The figure shows methylation density distributions in the ranges from 0-30% and 70-100% methylation from one fibroblast, one blood and one GIAB sample profiled with four different methylation detection methods (ONT, RRBS, TWIST, WGEC and PacBio(only GIAB)). Methylation values without coverage filtering are shown as filled ridgelines with black outlines, while red contours indicate methylation distributions after applying a 10× coverage threshold. After filtering, methylation values cluster more distinctly near 0 and 1, supporting the biological expectation of largely unmethylated or fully methylated CpGs. **(B):** Panel B employs the same analytical framework as subfigure A, focusing exclusively on ONT, PacBio and TWIST methodologies to facilitate more rigorous coverage filtering. The bimodal distribution pattern, with peaks at 0-5% and 94-98% methylation, indicates cellular heterogeneity in differentiation states, accounting for the absence of perfect 0% or 100% methylation levels across the population. As representative samples, the samples with the highest mean coverage were selected. Fibro := Fibroblast sample 4, Blood := Blood sample 3, GIAB := GIAB sample 1.

Across all samples and platforms, methylation values show the expected bimodal distribution, with peaks near 0 and 1, reflecting unmethylated and fully methylated CpGs, respectively. However, the shape and sharpness of these distributions vary substantially between methods and change markedly upon applying the coverage filter. Specifically, lower coverage data (’None’, i.e. unfiltered) exhibit broader, flatter distributions with platform-specific artifacts, such as an overrepresentation of intermediate methylation values in ONT-derived profiles, likely reflecting stochastic noise or alignment uncertainty at low read depths. In contrast, applying a 10× coverage threshold leads to sharper and more clearly separated density peaks, indicating improved precision in methylation calling. The effect is particularly pronounced for WGEC and TWIST, which display high consistency and narrow peak distributions post-filtering, while ONT still retains some residual variability. These findings underline that low-coverage methylation estimates are prone to noise and that a minimum coverage threshold is essential to recover the expected bimodal nature of DNA methylation.

This observation is corroborated by the correlation analysis across platforms (Figure 4), where inter-method Pearson correlation coefficients systematically increase with higher coverage thresholds. The ridge plots visually explain this trend: as the density distributions stabilize, the overlap between methylation profiles measured by different platforms becomes more coherent, thereby enhancing pairwise correlations. The ridge plot also suggests that platforms differ not only in global methylation levels but also in their coverage-dependent precision. For example, while RRBS and TWIST display sharper distributions already at the unfiltered CpG set, ONT shows significant improvements only after filtering, highlighting platform-specific sensitivities to coverage depth. Supplementary Figures S3 and S4 provide the corresponding per-sample density profiles underlying Figure 5: Figure S3 shows the full set of blood and fibroblast samples across ONT, RRBS, TWIST and WGEC before and after 10x filtering, while Figure S4 extends this comparison to the two GIAB samples and additionally overlays EPIC data (Blood3, Fibro4, GIAB1), illustrating that the pronounced intermediate-methylation peak is most evident for ONT in blood.

This pattern reflects the genomic architecture of DNA methylation, in which most CpG sites are either fully unmethylated or highly methylated, while intermediate states are comparatively rare. The presence of a main peak around 94% methylation rather than at 100% suggests that cell populations within each sample are not perfectly homogeneous, consistent with biological variability and cell-type–specific methylation differences. Increased coverage filtering (from 0 to 30 reads) generally reduces noise and slightly sharpens the density peaks, indicating that low-coverage CpGs contribute to broader or spurious intermediate methylation estimates. This effect is most apparent in the unfiltered data (Coverage Filter = ’None’), where distributions show more flattening toward intermediate methylation levels, particularly in lower-quality methods or tissues with higher heterogeneity.

Figure 5B focuses on ONT, PacBio and TWIST, which achieved higher mean coverage and thus permit more stringent filtering. The distributions become increasingly well defined with higher coverage thresholds and inter-method differences diminish, supporting the reliability of methylation estimates at moderate-to-high coverage. The persistent right-hand peak near 0.94–0.98 methylation fraction across methods confirms the biological consistency of the data, while the variable left-hand tail (near 0–0.1) reflects tissue-specific differences in unmethylated regions. Together with the correlation curves, the ridge density visualization emphasizes the dual role of coverage filtering: it mitigates platform-specific biases and enhances biological interpretability by suppressing noise.

### Comparative analysis of methylation profiling methods using principal component analysis underlines correlation concordance

To explore the combined effects of sequencing platform and inclusion of array-based methylation data on inter- and intra-sample variability, we performed a Principal Component Analysis (PCA) on methylation profiles from blood and fibroblast samples across five different measurement technologies: ONT, RRBS, TWIST, WGEC and EPIC (Figure 6). Each panel in the composite PCA figure represents the data after a coverage thresholds of 10× was applied and array inclusion (EPIC vs. No EPIC) across tissue types. Across all settings, PC1 consistently captures the dominant source of variance (*>*94%) and the resulting projections reveal a clear separation of the array-based method (EPIC) from all sequencing-based platforms (WGEC, TWIST, RRBS & ONT) along this axis, indicating that the primary axis for methylation variation in the dataset reflects a technology-class effect rather than sample-specific signatures. This separation along PC1 indicates a dominant platform-class effect, wherein array-based profiling (EPIC) diverges substantially from all sequencing-based methods(WGEC, TWIST, RRBS & ONT). Notably, sequencing-based platforms, cluster in close proximity to one another, suggesting that despite differences in library preparation, conversion chemistry and genomic coverage, they capture a largely concordant methylation signal.

**Figure 6:**
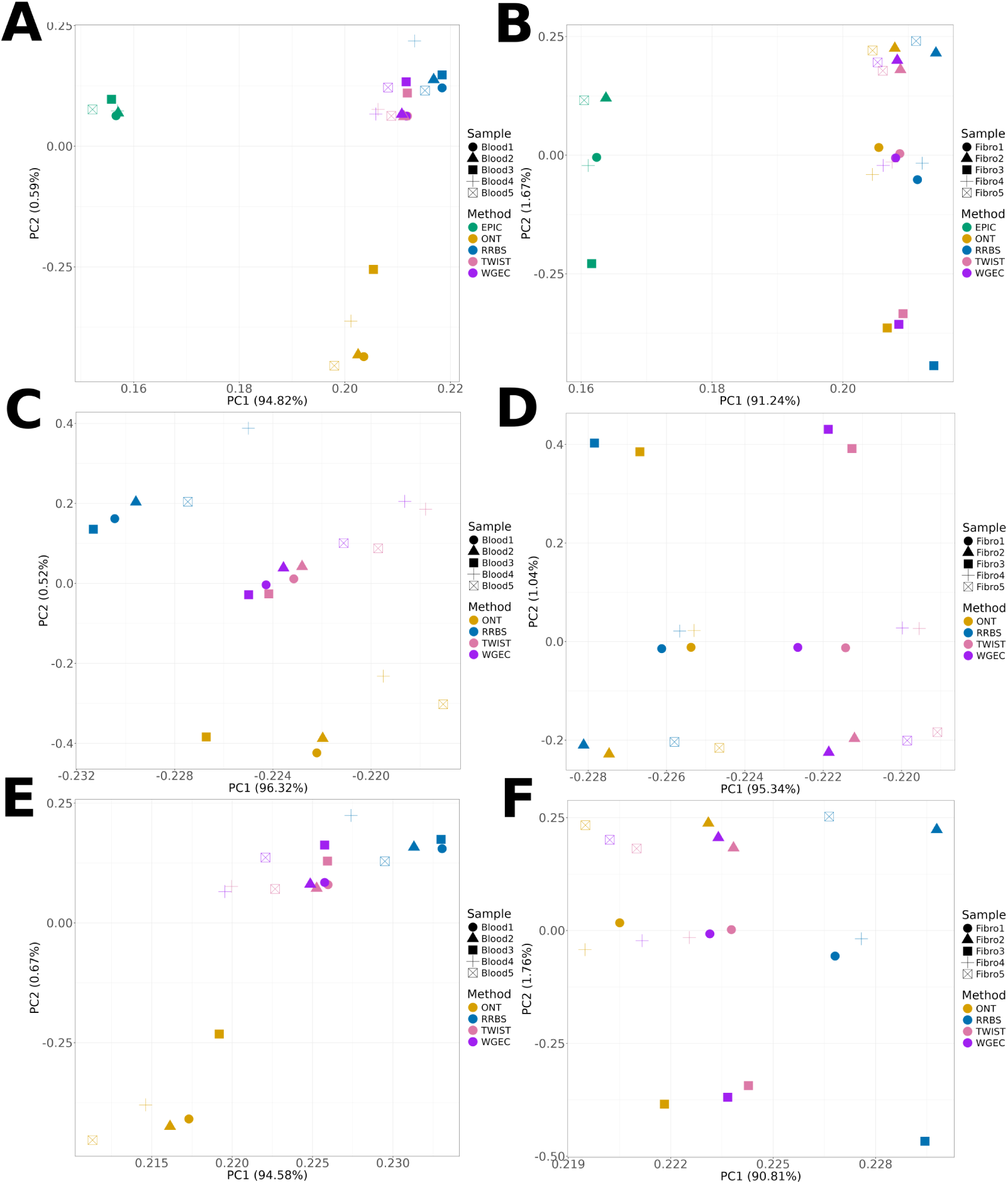
Principal component analysis of methylation profiles across tissues and methods under varying platform inclusion conditions. Principal Component Analysis (PCA) plots showing methylation profiles of five blood (subfigures A, C & E) and five fibroblast (subfigures B,D & F) samples, measured with five different technologies (ONT, RRBS, TWIST, WGEC and EPIC). Panels are organized by tissue type (columns) and inclusion of EPIC data (rows). Each point represents a sample-method combination, colored by platform and shaped by sample ID.

When EPIC data were excluded, blood and fibroblast samples each formed tight clusters(Figure 6C,D,E & F), indicating strong agreement among sequencing-based platforms (WGEC, TWIST, RRBS, ONT). Upon inclusion of EPIC array data (Figure 6A and 6B), the clustering pattern changes markedly. EPIC emerges as a clear outlier, even in blood samples where the overall concordance among sequencing-based platforms remains relatively high. This deviation is even more pronounced in fibroblast samples (Figure 6B), where EPIC-derived methylation profiles cluster separately from all sequencing-based datasets. The distinct position of EPIC in PCA space likely reflects its fixed CpG probe set and array-specific normalization effects, which limit its comparability with genome-wide sequencing approaches.

Interestingly, while some method-specific clustering is visible in blood (Panel A), the fibroblast PCA (Panel B) reveals a stronger separation by biological sample rather than by method, indicating that biological variability dominates over technical differences in this cell type.

These results suggest that while EPIC can provide broadly consistent results in promoter-rich blood-derived samples, its divergence from sequencing-based assays becomes more pronounced in fibroblasts, where intragenic and intergenic methylation differences contribute more strongly to the overall profile. Thus, EPIC’s restricted genome-wide coverage may limit its comparability in certain tissue types, particularly when whole-genome or gene body methylation is of interest.

To determine whether the technology-class separation observed for EPIC in blood and fibroblast samples (Figure 6) extends to the GIAB reference samples, and to assess how the inclusion of PacBio affects this picture, we performed an analogous PCA restricted to the five non-array methods (ONT, PacBio, RRBS, TWIST, WGEC) on the two GIAB samples, before and after applying a 10x coverage filter (Supplementary Figure S1). As in the blood/fibroblast analysis, PC1 captured the large majority of variance (88.1% unfiltered, 90.8% after filtering) and predominantly separated the two GIAB donors rather than methods, consistent with the sequencing-based clustering pattern in Figure 6. PacBio was a notable exception to this donor-driven structure: profiles from both GIAB1 and GIAB2 clustered together and remained offset from the corresponding donor-matched profiles of the other four methods, both before and after coverage filtering. This indicates that PacBio-derived methylation calls carry proportionally more method-specific technical variance relative to donor identity than ONT, RRBS, TWIST or WGEC, and that this separation is not resolved by coverage filtering alone - consistent with the coverage-independent PacBio discrepancy described in the correlation analysis (Figure 4B).

### Differential methylation analysis across platforms

To systematically evaluate cross-platform concordance in differential methylation detection, we compared five methylation profiling technologies: Oxford Nanopore Technologies (ONT) as a representative long-read sequencing approach, TWIST enzymatic conversion followed by capture-based enrichment and whole-genome enzymatic conversion (WGEC) as enzymatic conversion methods, reduced representation bisulfite sequencing (RRBS) as a standard bisulfite-based approach and the Illumina EPIC array, with respect to their ability to detect differentially methylated CpGs (DMCs) between matched blood and fibroblast samples from five individuals. PacBio was not included in this differential methylation analysis, as PacBio data were only available for the GIAB reference samples, which lack a matched blood-versus-fibroblast contrast required for this comparison.

#### Exploratory analysis: platform-dependent sensitivity under standard statistical models

As an initial screen, DMCs were identified on the full set of CpGs covered across all platforms (n=146,704) using two complementary statistical frameworks: limma, a moderated linear model with empirical Bayes variance shrinkage and a variance-robust Wilcoxon rank-sum test. The two approaches yielded markedly different pictures of cross-platform performance. Under limma, EPIC produced the largest number of significant DMCs, followed by TWIST, WGEC and RRBS, while ONT yielded comparatively few (Supplementary Figure S7); however, a substantial fraction of DMCs was shared across platforms, indicating that even under this model biologically consistent signal was recovered.

Under the Wilcoxon test, only EPIC and TWIST reached significance after multiple-testing correction, whereas ONT, RRBS and WGEC did not (Figure 7A). This divergence is not primarily biological but statistical: as shown in Figure 7C and 7D, ONT and the short-read sequencing platforms are characterized by substantially lower per-CpG coverage and higher variance than TWIST and EPIC.

**Figure 7:**
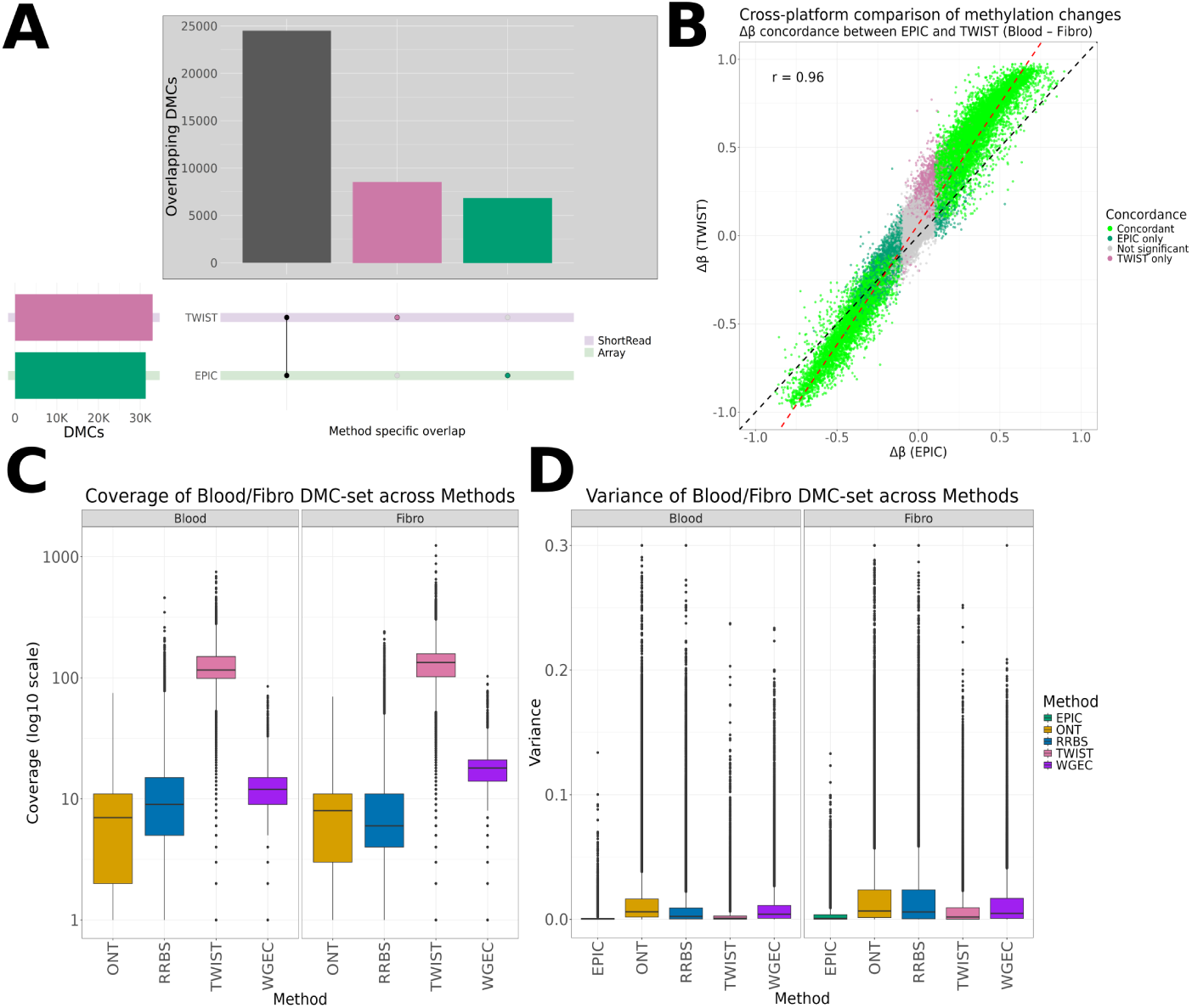
Differential CpG detection, coverage and variance across methylation platforms. (A): UpSet plot showing the overlap of significantly DMCs between blood and fibroblast samples, as identified using a wilcoxon test. Only EPIC and TWIST identify significant DMCs, demonstrating the reduced statistical power of ONT, RRBS and WGEC under non-parametric testing. **(B):** Cross-platform scatter plot comparing Δ*β* values between EPIC and TWIST. CpGs are colored by concordance: green indicates CpGs significant and directionally consistent in both platforms, red and blue mark EPIC- or TWIST-specific DMCs, respectively. The strong correlation (r = 0.96) indicates high cross-platform reproducibility under a variance-robust test. **(C):** Boxplots showing the mean coverage of CpGs included in each platform’s DMC set. Sequencing-based assays display broader coverage variability compared to EPIC, reflecting differences in sequencing depth and genomic sampling. **(D):** Variance of CpG methylation in blood and fibroblast samples across platforms. EPIC exhibits the lowest within-group variance due to array normalization, whereas sequencing-based platforms show higher dispersion linked to read depth and CpG representation.

Reduced coverage lowers the precision of individual *β*-value estimates and elevated variance disproportionately impairs a non-parametric test that, unlike limma, does not borrow statistical strength across CpGs via empirical Bayes shrinkage. limma therefore detects differential methylation across all five platforms, whereas the Wilcoxon test is only sufficiently powered for the two lowest-variance, highest-coverage technologies.

Despite this difference in detection power, the magnitude and direction of methylation change was highly concordant wherever it could be compared directly. Δ*β* values for EPIC and TWIST correlated at *r* = 0.96, with the large majority of CpGs falling close to the identity line (Figure 7B), indicating that both technologies capture essentially the same underlying biological effect and that most of the observed discrepancy in DMC counts reflects technical rather than biological variability. This concordance is corroborated by the near-identical Δ*β* density distributions across all five platforms (Supplementary Figure S6) and by hierarchical clustering of the top 500 and 1000 most variable CpGs, which consistently separates samples by tissue type regardless of platform (Supplementary Figure S8).

The platform-specific differences in coverage and variance underlying these results are themselves attributable to distinct library designs. TWIST’s hybridization-based enrichment, applied to the same underlying libraries as WGEC, yields more uniform, higher-depth coverage of predefined CpG regions and thereby reduces sampling noise, at the cost of a hybridization-driven bias toward expected genomic targets shared with array-based EPIC. WGEC, by design, distributes sequencing effort uniformly across the genome without targeted enrichment, resulting in lower and more variable per-CpG coverage. RRBS further restricts genomic scope to CpG-dense fragments obtained by restriction digestion (typically 4-12% of the genome at *≥* 1*x*), which increases per-CpG variance and reduces the number of CpGs surviving conservative non-parametric testing. EPIC, conversely, benefits from probe-level normalization and large-scale calibration, giving it exceptionally low variance and correspondingly high sensitivity even to modest effect sizes, at the cost of restriction to a fixed, curated probe set. CpGs located within promoters and CpG islands showed lower variance across all methods than those in enhancers or intergenic regions, consistent with differences in chromatin accessibility, GC content and mappability that disproportionately affect bisulfite/enzymatic sequencing-based assays.

#### Primary analysis: count-based DMC detection on a coverage-matched consensus set

The exploratory analysis demonstrates that DMC counts obtained under standard models are strongly shaped by platform-specific coverage and variance structure rather than by biology alone, and that treating *β*-values as continuous, homoscedastic outcomes disadvantages platforms with intrinsically noisier, count-derived measurements. To obtain a statistically appropriate and less confounded estimate of cross-platform concordance, we therefore performed a primary differential methylation analysis using DSS, which models methylated- and total-read counts directly via a Beta-Binomial distribution with empirical Bayes shrinkage of dispersion estimates, rather than treating derived *β*-values as continuous data. Because EPIC does not yield read counts, its differential methylation was instead assessed with limma on M-values (logit-transformed *β*-values), which stabilizes the array’s mean-variance relationship for linear modeling; this necessary difference in statistical treatment between sequencing- and array-based data was applied transparently and consistently throughout.

To remove the confounding effect of differential genomic coverage on DMC counts, this analysis was restricted to a platform-specific consensus CpG set, defined per platform as sites with *≥* 10*x* coverage (or, for EPIC, a non-missing *β*-value) in at least four of five samples in each tissue independently. Intersecting these per-platform sets yielded two tiers: Tier 1, comprising the four sequencing-based platforms (ONT, WGEC, TWIST, RRBS; n = 6,599 CpGs), used for sequencing-only concordance analysis; and Tier 2, additionally intersected with EPIC-covered sites (n = 6,598 CpGs), used for direct array-versus-sequencing comparison. DMCs were called after Gaussian-kernel smoothing (bandwidth = 500 bp) at FDR *<* 0.05 and |Δ*β| ≥* 0.1 (mean group difference for EPIC).

Figure 8 summarizes DMC concordance across the four sequencing-based platforms on the Tier 1 consensus set. Pairwise Δ*β* comparisons (Figure 8A) show tight clustering along the identity line across all six platform pairs, confirming that, once coverage and sample completeness are matched, sequencing-based technologies estimate near-identical effect sizes for blood-versus-fibroblast methylation differences. The UpSet plot (Figure 8B) shows that the majority of significant DMCs are shared across all four platforms, with comparatively small platform-exclusive fractions; this is quantified in Figure 8C, where the majority of each platform’s DMCs fall into the consensus category, and only a minority are detected by a subset of platforms or exclusively by one. The corresponding per-platform DMC counts, stratified by direction of methylation change and by genic annotation context, are provided in Supplementary Figure S14 and are consistent with the region-level patterns shown in Figure 9B–C. Figure 8D confirms that, by construction of the consensus set, coverage on the Tier 1

**Figure 8:**
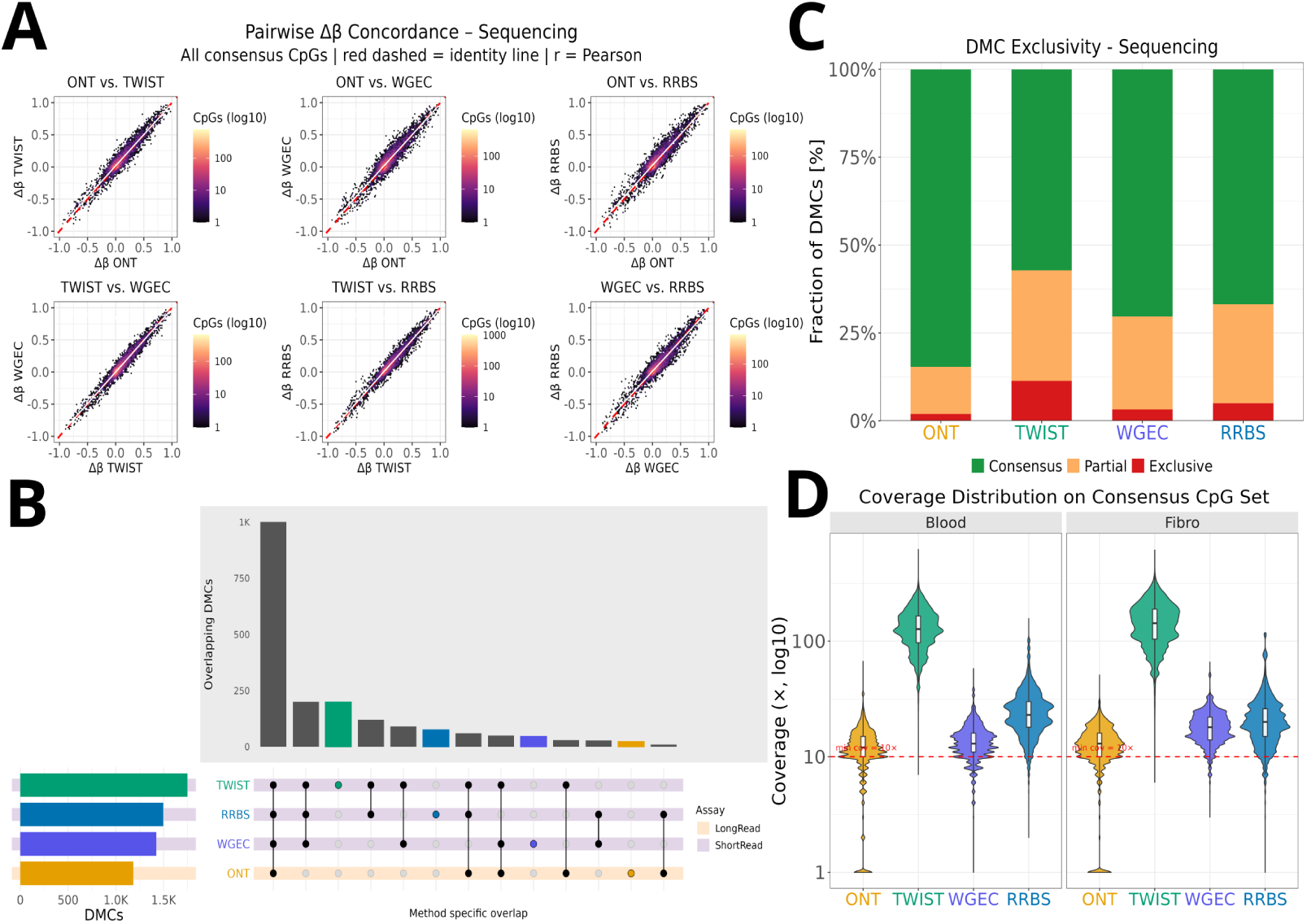
Differential methylation concordance across sequencing-based platforms at the CpG level. All analyses were performed on a consensus CpG set comprising sites with *≥* 10*x* sequencing coverage in *≥*4 of 5 samples per tissue across all four sequencing platforms (ONT, WGEC, TWIST, RRBS; n = 6,599 CpGs). DMCs were identified using DSS with a Beta-Binomial model (FDR *<* 0.05, |Δ*β| ≥* 0.1, smoothing bandwidth = 500bp). **(A):** Pairwise Δ*β* concordance across all consensus CpGs. Each panel shows the methylation difference (Blood - Fibroblast) estimated by one platform against another, with point density represented on a log10 color scale. The red dashed line indicates the identity line (slope = 1). **(B):** UpSet plot depicting the overlap structure of significant DMCs across platforms. Intersection bars indicate the number of DMCs shared by the indicated combination of methods; set size bars (left) show the total number of DMCs per platform. Row background shading indicates assay category (long-read: ONT; short-read: WGEC, TWIST, RRBS). **(C):** DMC exclusivity per platform. Stacked bars show the proportion of each platform’s DMCs that are detected by all four methods simultaneously (Consensus), by a subset of two to three methods (Partial), or exclusively by a single method (Exclusive). **(D):** Sequencing coverage distribution across platforms on the consensus CpG set, stratified by tissue. Each violin represents the per-CpG coverage distribution across all samples of a given platform and tissue; the red dashed line marks the minimum coverage threshold of 10x. Coverage is shown on a log10 scale.

**Figure 9:**
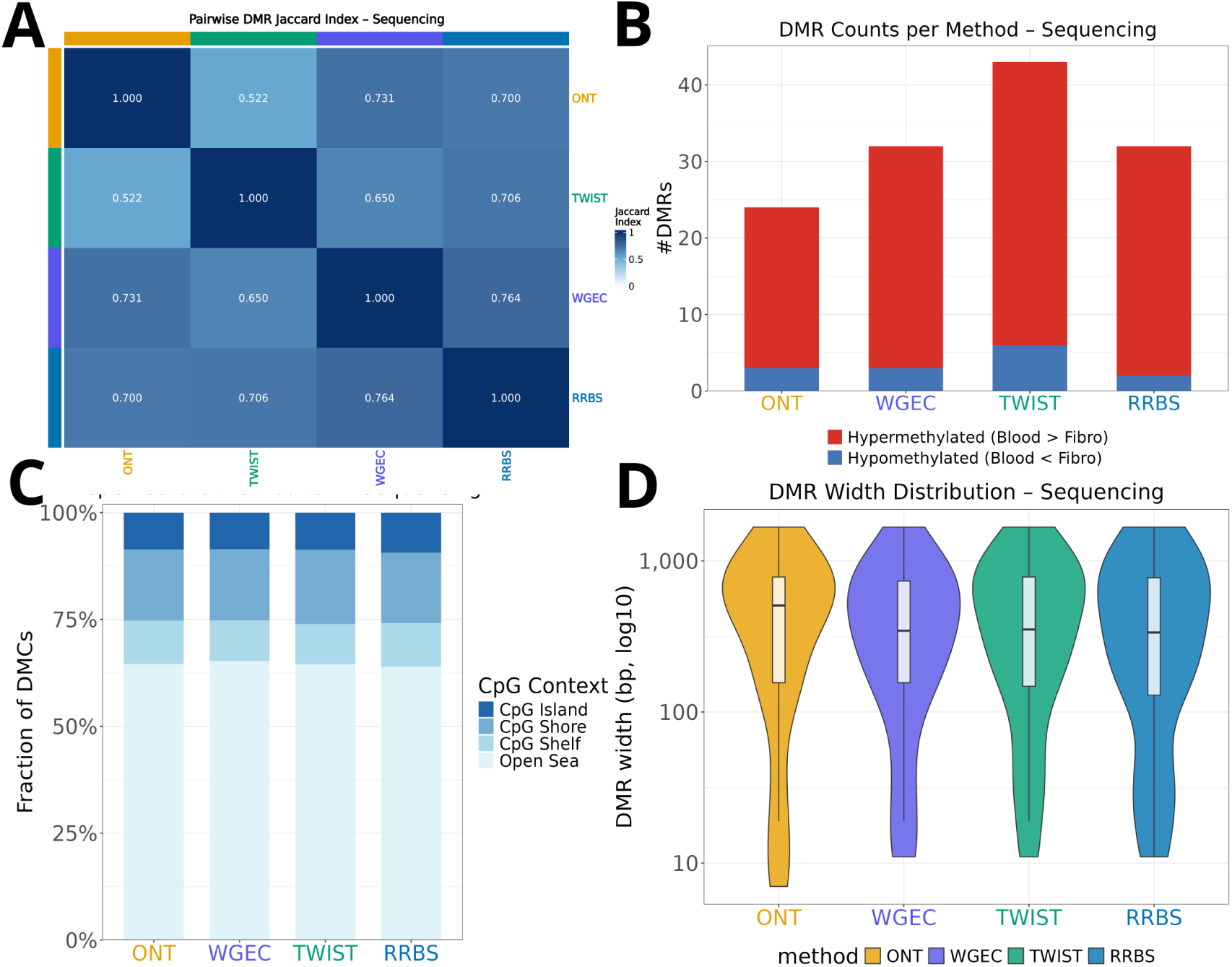
Differential methylation concordance across sequencing-based platforms at the genomic region level. DMRs were identified using DMRcate applied to CpG-level test statistics derived from DSS, with a kernel bandwidth of *λ* = 1,000bp, a minimum inter-CpG spacing of C = 2, and a minimum of three CpGs per DMR. All analyses were restricted to the only-sequencing consensus CpG set (n = 6,599 CpGs; see Figure 8). **(A):** Pairwise base-pair-weighted Jaccard similarity index between DMR sets across platforms. Values represent the ratio of intersecting to total base pairs across all pairwise platform combinations, quantifying genomic overlap of identified DMRs independently of region count or boundary precision. color scale ranges from 0 (no overlap) to 1 (complete overlap); row and column labels are color-coded by platform identity. **(B):** Number of DMRs identified per platform, stratified by direction of differential methylation. Red bars indicate regions hypermethylated in blood relative to fibroblasts (Δ*β >* 0); blue bars indicate hypomethylated regions (Δ*β <* 0). **(C):** Genomic CpG context distribution of identified DMRs per platform. Stacked bars show the proportion of DMRs annotated to CpG islands, shores, shelves, and open sea regions (hg38 reference). The absence of platform-specific enrichment in any CpG structural context indicates that inter-platform differences in DMR counts reflect differences in statistical sensitivity rather than systematic genomic coverage bias. **(D):** Distribution of DMR widths per platform shown on a log10 scale. Each violin represents the empirical width distribution of all identified DMRs; the embedded boxplot indicates the median and interquartile range.

CpGs is comparable and consistently above the 10x threshold across platforms and tissues, indicating that the residual differences in DMC number between platforms reflect remaining differences in statistical sensitivity rather than unequal access to the underlying CpGs. This concordance was independently corroborated using a threshold-free rank-recovery approach, in which the continuous DMC score from each platform was evaluated for its ability to recover the significant DMCs of every other platform as a reference (Supplementary Figure S15). AUC values ranged consistently between 0.77 and 0.81 irrespective of the chosen reference platform, confirming that cross-platform agreement is not an artifact of the specific significance thresholds applied and that ONT’s comparatively lower concordance reflects its higher underlying measurement variance (Figure 7D) rather than a threshold effect. Extending the CpG-level comparison to the EPIC-inclusive Tier 2 consensus set (Supplementary Figure S12) showed a qualitatively similar but somewhat attenuated pattern: pairwise Jaccard indices among the four sequencing-based platforms ranged from 0.64-0.74, whereas all EPIC-involving pairs were lower (0.57–0.65), and EPIC showed a markedly higher fraction of platform-exclusive DMCs (*≈*18%) than any sequencing-based technology (*≤*7%), mirroring its distinct clustering in the PCA (Figure 6).

#### Region-level validation: DMR concordance

CpG-level significance calls remain sensitive to local coverage gaps and to the stochastic nature of single-site detection, particularly given the still-heterogeneous per-CpG variance across platforms evident in Figure 8D. To assess whether cross-platform agreement is robust at a coarser, more biologically interpretable scale, we therefore aggregated the Tier 1 CpG-level test statistics into differentially methylated regions (DMRs) using DMRcate (Gaussian kernel bandwidth *λ* = 1000 bp, minimum inter-CpG spacing C = 2, minimum of three CpGs per DMR).

Figure 9 reports the resulting region-level concordance. Pairwise base-pair-weighted Jaccard indices (Figure 9A) -which quantify the overlap of DMR sets independent of region count or boundary precision - ranged from 0.52 to 0.76 across platform pairs, with WGEC, TWIST and RRBS showing the highest mutual overlap and ONT showing comparatively lower, but still substantial, overlap with each of the short-read platforms. The number of DMRs detected per platform (Figure 9B) mirrored the CpG-level sensitivity ranking (TWIST *>* WGEC *≈* RRBS *>* ONT), with a consistent predominance of hypermethylated regions (blood relative to fibroblast) across all four platforms. Genomic context annotation of DMRs (Figure 9C) showed no platform-specific enrichment for any CpG-structural category (island, shore, shelf, open sea), indicating that the observed differences in DMR count reflect differences in statistical power rather than a systematic genomic coverage bias of any single platform. DMR width distributions (Figure 9D) were broadly similar across platforms on a log-scale, with ONT showing a somewhat narrower interquartile range, consistent with its lower local CpG density on the consensus set.

Extending the DMR-level analysis to the EPIC-inclusive Tier 2 set (Supplementary Figure S13) revealed a more pronounced divergence than at the CpG level. Pairwise DMR Jaccard indices between EPIC and the four sequencing-based platforms (0.38-0.46) were consistently lower than the corresponding CpG-level Jaccard indices (Supplementary Figure S12: 0.57-0.65) and than the sequencing-only pairwise values (0.52-0.76), indicating that region aggregation amplifies boundary- and clustering-related discordance for the array-based platform. More notably, EPIC-derived DMRs were predominantly hypomethylated in blood relative to fibroblast (25 of 41 DMRs), in contrast to all four sequencing-based platforms, which were consistently hypermethylation-dominant (ONT: 21/24; WGEC: 29/32; TWIST: 37/43; RRBS: 30/32 hypermethylated). This directional asymmetry was already detectable at the CpG level (Supplementary Figure S14B: 36% of EPIC DMCs hypomethylated versus 14–20% for sequencing-based platforms), indicating that it reflects a genuine platform-specific measurement property rather than an artifact of DMR calling.

Taken together, the three-tiered analysis - exploratory CpG-level testing, primary count-based DMC detection on a coverage-matched consensus set, and region-level DMR validation - converges on the same conclusion from progressively more stringent and less confounded angles: differences in the number of significant methylation differences detected across platforms are predominantly attributable to platform-specific coverage and variance structure and to the assumptions of the applied statistical model, whereas the underlying biological methylation signal - its magnitude, direction and genomic localization - is highly concordant across all sequencing-based technologies and, where directly comparable, between sequencing and array-based profiling.

## Discussion

In this benchmarking study, we systematically compared six methylation profiling technologies - EPIC, TWIST, WGEC, RRBS, ONT, and PacBio - with respect to coverage, variance structure, CpG representation, and differential methylation detection. Despite substantial differences in assay chemistry, resolution, and genomic scope, all technologies consistently identified differentially methylated CpGs (DMCs) in promoter and intronic regions (Supplementary Figure S9). This convergence underscores that core biological methylation signatures are robustly captured across fundamentally distinct technologies. Promoter-associated DMCs likely reflect canonical transcriptional regulation, whereas the substantial fraction of intronic DMCs suggests broader epigenetic mechanisms related to splicing regulation, chromatin accessibility, and transcriptional elongation. This pattern held not only at the aggregate, redundancy-collapsed level (Supplementary Figure S9) but also when genic context was resolved per platform and per Tier (Supplementary Figure S14C–D), with intronic and promoter enrichment remaining stable across all five technologies irrespective of assay chemistry or genomic scope.

Our results also highlight how annotation redundancy can skew downstream interpretation. CpG-island–based categories (islands, shores, shelves) frequently overlapped with promoter or gene-dense regions, and their apparent enrichment largely disappeared when annotations were collapsed to unique genomic categories (Supplementary Figure S9). This points to the importance of harmonized annotation strategies when comparing platforms that differ in CpG sampling density or genomic coverage.

The methodological comparison revealed clear platform-specific strengths. Sequencing-based assays, particularly WGEC and TWIST, provided broad genomic coverage and were effective in detecting intronic and intergenic methylation changes. ONT and PacBio further offer unique advantages, including long-read phasing[25][38] of methylation and the potential for simultaneous detection of additional DNA modifications; however, these capabilities were beyond the scope of the present study and were therefore not explored here. A particularly notable finding of this benchmark concerns PacBio’s coverage-independent discordance with other platforms. Contrary to the general expectation that higher sequencing depth improves cross-platform agreement, GIAB1 - sequenced at the highest PacBio coverage observed in this study (92x) - showed the lowest concordance with the remaining platforms (*r ≈* 0.83 *−* 0.84), whereas GIAB2, sequenced at substantially lower coverage (38x), showed markedly better agreement (*r ≈* 0.92; Figure 4B). This inverse relationship between coverage and concordance indicates that the residual discrepancy is not primarily a sampling-noise phenomenon and is therefore not expected to resolve with further increases in sequencing depth. One methodological consideration that should temper a purely technology-level interpretation of this finding is that the GIAB1 and GIAB2 PacBio datasets used here originate from two distinct sequencing efforts that differ not only in nominal chemistry but in instrument software generation: GIAB1 (HG001) was obtained from the Platinum Pedigree long-read benchmarking resource, generated using standard Revio SPRQ chemistry, whereas GIAB2 (HG002) was generated using the newer SPRQ-Nx multi-use chemistry with three acquisitions per SMRT Cell and an updated on-instrument base-calling model that additionally supports 5hmC and 6mA detection alongside improved 5mC calling. Because only 5mC calls were used in the present analysis, the functional comparison between samples is nominally restricted to this single modification; however, the underlying basecalling model differs between the two datasets and improvements to 5mC calling accuracy specifically noted for the SPRQ-Nx release cannot be excluded as a contributing factor to the higher concordance observed for GIAB2 relative to GIAB1 (Figure 4B). This cross-cohort heterogeneity in chemistry and basecalling model generation cannot be fully disentangled from a genuine, coverage-independent PacBio measurement property within the scope of this study, and should be considered when interpreting the magnitude of the observed discordance between GIAB1 and GIAB2, even though it does not affect the central conclusion that increasing coverage alone did not resolve cross-platform disagreement for either sample. Notably, native sequencing on ONT is more susceptible to impurities than other platforms and improvements in sequencing chemistry and basecalling algorithms may further enhance CpG coverage; yet, such refinements were not specifically addressed in the current benchmarking. However, these methods also exhibited higher variance and more heterogeneous coverage, which can reduce statistical sensitivity. Notably, WGEC libraries included internal positive and negative methylation controls in each reaction, enabling per-sample estimation of enzymatic conversion efficiency during downstream analysis. While such controls cannot fully account for sample-specific effects such as DNA damage or chromatin accessibility, they provide an additional layer of technical validation and confidence in the robustness of the WGEC workflow.

In contrast, the EPIC array showed highly stable probe-level measurements, reflected by consistently low methylation variance across samples. Although limited to a predefined probe set, EPIC reliably captured promoter-associated differences and produced a large number of significant DMCs. Extending the concordance analysis to the EPIC-inclusive Tier 2 consensus set showed that this comes with a measurable, systematic cost to cross-platform agreement: pairwise DMC Jaccard indices between EPIC and the four sequencing-based platforms (0.57–0.65) were consistently lower than those among sequencing-based platforms themselves (0.64–0.74; Supplementary Figure S12B), and EPIC displayed the highest proportion of platform-exclusive DMCs of any technology (*≈*18%, versus *≤*7% for all sequencing-based assays; Supplementary Figure S12C). This divergence was substantially more pronounced at the level of aggregated regions than at individual CpGs: DMR-level Jaccard indices between EPIC and sequencing-based platforms fell to 0.38–0.46 (Supplementary Figure S13A), indicating that boundary effects introduced during region aggregation compound, rather than average out, EPIC’s platform-specific measurement properties. More strikingly, EPIC-derived DMCs and DMRs showed a pronounced directional asymmetry relative to all sequencing-based platforms. Whereas ONT, WGEC, TWIST and RRBS consistently identified predominantly hypermethylated regions in blood relative to fibroblast (proportion hypermethylated: 88–93% of DMRs per platform), EPIC-derived DMRs were majority hypomethylated (61%; Supplementary Figure S13B), and this asymmetry was already present, albeit less pronounced, at the level of individual DMCs (36% hypomethylated for EPIC versus 14–20% across sequencing-based platforms; Supplementary Figure S14B). Because this directional bias emerges specifically at the level of called significant sites rather than in the underlying continuous effect-size estimates - which remained highly concordant between EPIC and TWIST (r = 0.96; Figure 7B) - it is unlikely to reflect a true biological difference and more plausibly arises from EPIC’s fixed probe set and array-specific normalization procedure differentially affecting the detectability of hypo- versus hypermethylation events at significance. This distinction between concordance in continuous effect size and concordance in binary significance calls has direct practical implications: analyses relying on discrete EPIC-derived DMR/DMC calls to infer the direction of tissue-specific methylation differences should be interpreted with this asymmetry in mind, and, where feasible, validated using orthogonal measurements such as targeted sequencing or expression-based correlation analyses.

A central finding of this work concerns the interaction between platform-specific measurement properties and statistical modeling. limma’s empirical Bayes variance shrinkage is well suited for array data but disproportionately favors methods with stable, low-variance measurements. Accordingly, EPIC produced the highest number of significant DMCs under limma, whereas sequencing-based assays, which intrinsically exhibit broader variance due to heterogeneous read depth, yielded fewer significant hits. This imbalance reflects not only biological and technical differences but also the assumptions embedded in the statistical model.

To mitigate this bias, we applied a variance-robust Wilcoxon rank-sum test. Unlike limma, the Wilcoxon test makes no assumptions about variance structure or normality and thereby equalizes sensitivity across heterogeneous datasets. Under this framework, cross-platform agreement improved substantially: sequencing-based methods, previously dispersed under limma, clustered closely and showed highly concordant Δ*β* patterns. EPIC still formed a distinct cluster, likely reflecting its predefined CpG set and array-specific normalization, yet its Δ*β* values were directionally aligned with sequencing methods on overlapping CpGs. This demonstrates that most biological methylation differences between blood and fibroblasts are consistently detected across platforms when statistical tests are chosen appropriately. This conclusion was further corroborated using a threshold-free rank-recovery analysis, in which the ability of each sequencing-based platform’s continuous DMC score to recover another platform’s significant DMCs was quantified independently of any specific significance cutoff (Supplementary Figure S15). AUC values remained stable within a narrow range (0.77–0.81) across all twelve pairwise reference–comparison combinations, confirming that the cross-platform concordance reported above is not contingent on the particular FDR and effect-size thresholds applied and providing an internally consistent, threshold-independent estimate of platform agreement to complement the Beta-Binomial and Jaccard-based analyses.

Several limitations must be considered. First, our study includes only ten biological samples across two tissue types and two GIAB references. Although sufficient for benchmarking, larger and more diverse cohorts would allow more robust characterization of platform-specific variance and DMC reproducibility. Second, GIAB samples are deeply characterized and may not reflect coverage and noise profiles observed in routine sequencing studies, potentially biasing comparisons toward sequencing platforms. Third, we focused on overlapping CpGs to ensure comparability; however, the biological relevance of platform-specific DMCs - particularly those unique to EPIC or ONT - remains unresolved and warrants validation through integrative omics approaches such as RNA-seq or ATAC-seq. Fourth, our analysis primarily examined CpG-level methylation; emerging platforms allow quantification of non-CpG methylation and additional DNA modifications (e.g. 5hmC), which may reveal further layers of cross-platform variability. Fifth, the directional asymmetry observed for EPIC-derived DMCs and DMRs (Supplementary Figures S13, S14) could not be further dissected within the scope of this study; disentangling contributions of probe-level normalization, background correction and the curated, promoter-biased nature of the EPIC probe design would require dedicated spike-in or simulation-based experiments beyond the present benchmarking framework. Sixth, the two PacBio GIAB datasets originate from distinct sequencing efforts that differ not only in donor but in chemistry and instrument software generation (standard Revio SPRQ chemistry with Platinum Pedigree processing for GIAB1 versus SPRQ-Nx multi-use chemistry with an updated basecalling model for GIAB2); this cross-cohort heterogeneity cannot be fully disentangled from a genuine, coverage-independent PacBio measurement property within the scope of this study and should be considered when interpreting the magnitude, though not the direction, of the observed discordance. Seventh, blood/fibroblast samples were profiled using the Infinium MethylationEPIC BeadChip version 1, whereas GIAB samples were profiled using version 2, as version 1 was no longer commercially available at the time of GIAB processing; because the two array versions differ in probe content and design, this represents an analogous cross-cohort confound to the PacBio chemistry discrepancy described above and should be considered when comparing EPIC-specific results between the GIAB and blood/fibroblast cohorts.

Despite these limitations, our findings emphasize that promoter and intronic regions represent robust methylation hotspots consistently captured across all technologies. For practical applications, EPIC arrays remain a cost-effective solution for large promoter-focused cohort studies, TWIST and WGEC provide comprehensive genome-wide coverage, and ONT enables phasing and multimodal measurements but currently requires deeper coverage for optimal DMC detection. Ultimately, the choice of platform should be driven by study objectives: genome-wide discovery, targeted interrogation, or haplotype-resolved epigenomics.

Taken together, this study shows that careful consideration of coverage thresholds, genomic annotation, and statistical modeling is essential for unbiased interpretation of methylation data across platforms. Variance-robust approaches such as the Wilcoxon test, and threshold-free rank-recovery analyses more broadly, can substantially improve cross-platform agreement and reveal the underlying biological consistency masked by model assumptions. However, this consistency is not unconditional: array-based profiling in particular can diverge from sequencing-based consensus specifically at the level of discrete significance calls, even when underlying continuous effect sizes remain concordant, underscoring that platform-comparison studies should assess concordance at multiple levels of analysis — continuous effect size, called significant sites, and aggregated regions — rather than any single one in isolation. Future benchmarking integrating larger cohorts, multiple tissues, and functional readouts will further refine these conclusions and guide the design of reliable methylation studies.

## Conclusion

This comprehensive benchmark demonstrates that DNA methylation profiling technologies differ markedly in genome-wide coverage, CpG representation and variance structure, yet converge on highly consistent biological signals when appropriate statistical frameworks are used. limma, while highly effective for microarray data, inflates sensitivity for low-variance platforms such as EPIC and underestimates differential methylation in sequencing-based assays. In contrast, the Wilcoxon rank-sum test harmonizes sensitivity across heterogeneous datasets and reveals strong cross-platform reproducibility of Δ*β* estimates, including near-perfect agreement between EPIC and TWIST.

Sequencing-based methods provide in general unparalleled genomic breadth, capturing intronic and intergenic methylation dynamics inaccessible to arrays, whereas EPIC remains a robust and cost-efficient solution for promoter-focused cohort studies. This robustness applies most clearly to EPIC’s continuous methylation estimates and their concordance with sequencing-based effect sizes; at the level of discrete DMC and DMR calls, EPIC showed both reduced overlap with sequencing-based platforms and a disproportionate tendency toward hypomethylation calls relative to all four sequencing-based technologies, a discrepancy not attributable to coverage and warranting cautious interpretation when EPIC-derived significant calls are used in isolation to infer the direction of tissue-specific methylation change. Across all platforms, coverage emerged as a key determinant of reproducibility: a pragmatic minimum CpG coverage threshold of approximately 10× consistently reduced noise and improved cross-platform agreement while retaining sufficient genomic representation. However, the benefit of stricter filtering was method dependent - short-read assays were largely robust, ONT required higher and more uniform coverage to mitigate variance, and PacBio exhibited residual discrepancies that were not fully resolved by increasing coverage alone - highlighting that coverage thresholds represent technology-specific quality constraints rather than universal guarantees of concordance. A summary of the recommended minimum CpG coverage thresholds per technology, together with the empirical rationale underlying each recommendation, is provided in Table S1. This was most evident in GIAB1, where the highest PacBio coverage observed in this study (92x) coincided with the lowest cross-platform concordance (*r ≈* 0.83–0.84; Figure 4B), directly illustrating that coverage alone does not guarantee cross-platform agreement. Conversely, GIAB2, sequenced at less than half the PacBio coverage of GIAB1 (38x), showed markedly better agreement with the other platforms (*r ≈* 0.92), underscoring that cross-platform concordance for PacBio in this dataset was not predictable from coverage alone. Overall, our findings underscore the importance of coverage-aware filtering, redundancy-aware genomic annotation and variance-robust statistical modeling for reliable and integrative methylation analyses. The framework presented here provides practical guidance for selecting suitable methylation technologies based on research aims, biological context, and resource constraints.

## Materials & Methods

### Sample collection and study design

Matched blood and fibroblast samples from five human individuals were analyzed using six DNA methylation profiling technologies: whole-genome enzymatic conversion (WGEC) using the NEBNext Enzymatic Methyl-seq Kit (New England Biolabs, Ipswich, USA), Twist Targeted Methylation Sequencing Workflow (Twist Bioscience, South San Francisco, USA), reduced representation bisulfite sequencing (RRBS), Illumina EPIC array and Oxford Nanopore Technologies (ONT). Two of the samples (HG001 and HG002)[18] were additionally obtained from the Genome in a Bottle (GIAB) consortium and included PacBio methylation data to enable multi-platform validation. All samples were processed using standardized workflows and platform-specific pipelines were applied to generate single-CpG resolution methylation calls.

### Sample collection and DNA extraction

DNA from EDTA blood of 5 probands was extracted using the Flexigene kit (Qiagen) according to manufacturer’s instructions. Fibroblasts of the same 5 individuals were cultured under standard conditions (5% CO2 and 100% humidity) in 75cm2 flasks and harvested when 80% confluent. DNA was extracted using the Monarch® HMW DNA Extraction Kit for Cells & Blood (New England Biolabs).

DNA concentration was measured using the Qubit Fluorometer and dsDNA High sensitivity assay (both Thermo Fisher Scientific, Waltham, USA). Genomic integrity was assessed using pulsed-field capillary electrophoresis with the Genomic DNA 165 kb Analysis Kit on a FemtoPulse (Agilent Technologies, Santa Clara, USA) instrument.

### DNA Preparation and Sequencing

#### Next Generation Sequencing (WGEC and Twist Methylome)

Enzymatic Conversion was performed using 200 ng of DNA. Per sample, internal conversion controls pUC and Lambda, provided in the NEBNext® Enzymatic Methyl-seq Kit (New England Biolabs, Ipswich, USA) were diluted and added according to the vendor’s protocol. Subsequently, samples were sheared using a Covaris E220 ultrasonicator (Covaris, Woburn, MA, USA) and subjected to enzymatic conversion and NGS library preparation with the NEBNext® Enzymatic Methyl-seq Kit (New England Biolabs, Ipswich, USA). All libraries were divided by volume into two similar fractions: one to be sequenced without further treatment and one to be enriched by using the Twist Targeted Methylation Sequencing Workflow (Twist Bioscience, South San Francisco, USA). In brief, 250 ng of the above-mentioned libraries was used to perform the hybridization-based enrichment with the Twist Human Methylome Panel and the corresponding workflow provided by the manufacturer (Twist Bioscience, South San Francisco, USA). Enriched libraries were amplified and a cleanup was performed.

All enzymatically converted libraries (WGEC and Twist) were quantified using the Qubit Fluorometer and dsDNA High sensitivity assay (both Thermo Fisher Scientific, Waltham, USA) and average library lengths were determined using the Tapestation 4200 and the High Sensitivity D1000 ScreenTape Assay (Agilent Technologies, Santa Clara, USA). From these data, libraries were pooled in an equimolar fashion. All WGEC and Twist libraries were sequenced on the same sequencing run and on the same S2 Flow Cell on an Illumina NovaSeq 6000 (Illumina, San Diego, CA, USA) in paired-end mode with 2x150 base pairs (bp).

#### Illumina EPIC array

From each sample, 500 ng of genomic DNA was treated with bisulfite using the EZ DNA Methylation kit (D5002, Zymo Research), according to the manufacturer’s recommendations and with a final elution volume of 10 *µ*l of elution buffer. For each sample, 4 *µ*l of bisulfite converted DNA was used as input for the Infinium MethylationEPIC BeadChip (Illumina, San Diego) according to the vendor’s protocol and the microarrays were scanned on the Illumina HiScan platform. For the blood/fibro samples EPIC version 1 was used and for GIAB samples EPIC version 2 was used (as version 1 was not available for purchase anymore).

#### RRBS

RRBS was performed as described previously (Mohan et al. 2023[28]). Briefly, 200 ng of genomic DNA from each sample was subjected to restriction with 1*µ*l HaeIII enzyme (50 U/*µ*l; New England Biolabs; NEB) and 3*µ*l 10x Cutsmart buffer (NEB) at 37° C for 18 hours. A-tailing was performed with 1*µ*l Klenow fragment (3’ *→* 5’exo-, 5 U/*µ*l, NEB) and 1*µ*l dATP (10 mmol/l, NEB) at 37°C for 30 minutes, followed by enzyme inactivation at 75° C for 20 minutes. Fragmented DNA samples were purified using 50 *µ*l AMPure XP Beads (Beckman Coulter) following the manufacturer’s protocol and a final elution with 36.5 *µ*l of nuclease-free water (saving 34.5 *µ*l to a new tube). NGS adapter ligation was done by adding 1*µ*l of NGS adapter (10*µ*mol/l; TruSeq Single Index Set B; Illumina), 0.5*µ*l of T4 Ligase (2000 U/*µ*l, NEB) and 4 *µ*l of 10x T4 ligation buffer (NEB) and incubated at 16° C for 18 hours, followed by enzyme inactivation at 65° C for 20 minutes. Bisulfite conversion and subsequent cleanup were performed using EZ-DNA Methylation Gold Kit (Zymo Research) according to the manufacturer’s instructions with a final elution in 24 *µ*l nuclease-free water. Each library was then PCR-amplified by combining 23 *µ*l of library, 1 *µ*l of each primer (10 *µ*mol/l, primer i5: AATGATACGGCGACCACCGAGATCTACAC, primer i7: CAAGCAGAAGACGGCATACGAGAT) and 25 *µ*l of the KAPA HiFiHotStart Uracil+ ReadyMix (2x; Roche) with the following thermo-protocol: initial 95° C for 3 minutes, then 15 cycles of 98° C for 20 s, 58° C for 30 s, 72° C for 1 minute, and then 72° C for 12 minutes, and final hold at 4° C. The amplified RRBS libraries were purified using 0.8x volume (40 *µ*l) of AMPure XP Beads according to the manufacturer’s instruction and eluted in 12 *µ*l of 0.1x Tris-HCl buffer. Library concentrations were measured using the Qubit HS DNA kit (Life Technologies) and the NEBnext library quant kit for Illumina (NEB). Sequencing was performed on an Illumina NextSeq500 platform (1x 75nt) with additional sequencing on the Aviti24 platform (Element Biosciences).

#### LR-GS ONT

A total of 3.25 *µ*g of genomic DNA was sheared with Megaruptor 3 (Diagenode) for 5 blood and 5 fibroblasts. Libraries were prepared with the 1D Ligation SQK-LSK109-XL Sequencing kit (Oxford Nanopore Technologies) and assessed with the Genomic DNA 165-kb Analysis Kit on a FemtoPulse (Agilent) instrument. A total of 600 ng (50 fmol) of each library was loaded on each PromethION R9 flow cell. For blood DNA two flow cells were used whereas for fibroblasts a single flow cell was required to reach equivalent sequencing yield. The GIAB samples were prepared using 1D Ligation SQK-LSK114-XL Sequencing kit and sequenced using two PromethION R10 flow cells for GIAB1 and a single flow cell for GIAB2.

#### LR-GS PacBio

PacBio HiFi whole-genome sequencing data for GIAB2 (HG002) were obtained from the publicly available PacBio SPRQ-Nx dataset[34]. Libraries were sheared by pipette-tip shearing on a Hamilton NGS STAR MOA 96 system and prepared using the HiFi prep kit 96 on the same automation platform[34]. Sequencing was performed on the Revio system using SPRQ-Nx multi-use chemistry with three acquisitions per SMRT Cell and 24-hour movie collection[34]. Base and methylation calling was performed on-instrument using the corresponding release instrument software; secondary analysis, including read alignment, was performed using the standard PacBio HiFi WGS variant-calling pipeline[34]. Of the three modification channels available on-instrument, only 5mC calls were retained for the analyses presented here, consistent with the CpG-methylation scope of this benchmark; 5hmC and 6mA calls were not further analyzed.

PacBio HiFi data for GIAB1 (HG001/NA12878) were obtained from the Platinum Pedigree dataset, generated as part of a long-read benchmarking effort across the four-generation CEPH-1463 pedigree, of which HG001 is a member[20]. This dataset combines deep PacBio HiFi, Oxford Nanopore, and Illumina sequencing across the pedigree to construct a haplotype-resolved, Mendelian-inheritance-validated variant benchmark[19]. HG001 HiFi data used in this study were downloaded from the Platinum Pedigree Amazon Open Data repository as documented in the associated GitHub repository[20]. Chemistry and kit version details for the Platinum Pedigree HG001 PacBio HiFi data are as documented in [19][20]; direct chemistry-generation matching with the GIAB2 dataset could not be confirmed and is noted as a limitation (see Discussion).

### Methylation Calling and Preprocessing

All sequencing data were aligned to the GRCh38 human reference genome.

For ONT datasets, methylation calls were extracted from BAM files using modbam2bed[12] and processed using the modkit[30] suite, with default parameters, to generate per-CpG methylation summaries. Bismark[21] was used for alignment and methylation extraction for WGEC, RRBS and TWIST datasets. PacBio data was processed using Pacific Biosciences pb-CpG-tools software[33] with default settings. For the EPIC, the obtained raw data was preprocessed using the RnBeads R/Bioconductor package (v2.12.2)[1][29]. For the fibroblast samples 3,691 probes (GIAB: 15,941) were removed using the Greedycut algorithm, based on a detection p-value threshold of 0.05, as implemented in the RnBeads package. Also, 17,371 (GIAB: 0) CpG sites were filtered as their detection probes overlapped with 3 or more annotated SNPs (RnBeads setting ’filtering.snp = 3’) and 2,985 (GIAB: 3,724) methylation sites from non-CpG contexts were removed (RnBeads setting ’filtering.context.removal = CC,CAG,CAH,CTG,CTH,Other’). In addition, 144 (GIAB: 0) CpGs were removed because they were missing in more than 80% of samples (RnBeads setting ’filtering.missing.value.quantile *>* 0.8’). As a final outcome of the filtering procedures 24.191 sites (GIAB: 19,665) and 0 samples were removed, resulting in 823,649 (GIAB: 910,475) methylation sites in the data set for subsequent analysis. For each CpG site, a beta-value was calculated, which represents the fraction of methylated cytosines across all detected molecules (0 = unmethylated, 1 = fully methylated). Subsequently, beta-values were normalized using the watermelon dasen[36][43] normalization method as implemented in RnBeads. Differential pairwise comparisons were done using a two-sided Student’s t-test. Ranking of statistical signals was done with the combined rank method as implemented in RnBeads, which uses a combinatorial ranking based on the nominal p-value from the t-test and the absolute delta of the average methylation of the two compared groups. All plots were done in R.

To harmonize methylation measurements across platforms, CpG sites were filtered to include only those with unambiguous genomic coordinates. CpG-level beta values (methylated reads / total reads) were calculated per sample. Data from all technologies were then merged into common matrices using data.table[2] in R.

### Quality control and coverage filtering

Quality control metrics included read length, mean CpG coverage per sample, number of unique reads and fraction of CpGs with sufficient coverage. To assess the impact of sequencing depth, coverage filters (*≥*10×, *≥*15×,

*≥*20×, *≥*30×, *≥*40×) were applied to the merged methylation tables. Coverage distributions were visualized using boxplots and ridge density plots (ggridges[45]).

### Principal component and correlation analyses

Principal component analysis (PCA) was conducted using the prcomp function in base R after filtering for CpGs commonly covered across platforms. Pairwise Pearson correlation coefficients were computed per sample on matched CpG sites using cor with complete pairwise observations. For additional robustness, correlation analyses were repeated under multiple coverage thresholds.

### Differential methylation analysis

#### Exploratory Analysis: limma-based differential methylation and Wilcoxon Rank-Sum Test

Initial differential methylation analysis was performed using CpG-level methylation matrices generated from each platform (ONT, RRBS, WGEC, EPIC and TWIST). Each matrix contained genomic coordinates (chromosome, start position) and per-sample methylation proportions. For comparability, all analyses were restricted to the common set of CpG sites covered across platforms (n=146,704 CpGs).

Statistical testing was carried out in R using the limma package[39], which applies an empirical Bayes approach to moderated linear models. For each CpG, methylation values were modeled as a function of experimental condition (e.g., Blood vs. Fibroblast), with batch information accounted for when applicable. For comparisons involving more than two biological conditions (e.g., Blood, Fibroblast, GIAB), a design matrix reflecting group membership was constructed and contrasts were extracted for relevant pairwise comparisons.

Significance thresholds were set at an adjusted p-value *<* 0.05 (Benjamini–Hochberg correction) and an absolute Δ*β ≥* 0.1 unless otherwise specified, consistent with the effect-size threshold applied in the primary DSS-based analysis (see below). Results were visualized using UpSet plots (UpSetR[8], ComplexUpset[22] & ggplot2[44]) to assess both the magnitude and overlap of differential methylation across platforms.

As a variance-robust alternative, differential methylation was also assessed using a Wilcoxon rank-sum test applied independently to each CpG. This non-parametric test does not rely on normality or equal variances and is therefore more appropriate for comparing platforms with heterogeneous measurement noise. For each CpG, the Wilcoxon test evaluated whether *β*-values in blood and fibroblasts originated from the same distribution and the resulting p-values were again corrected using Benjamini–Hochberg FDR. CpGs with FDR *<* 0.05 were classified as significant, with Δ*β* used to assess effect size and directionality. Because the Wilcoxon test is unaffected by platform-specific variance inflation or shrinkage, it provides a more balanced statistical sensitivity across sequencing- and array-based assays. This approach led to noticeably improved cross-platform concordance of effect sizes and was therefore used as the primary basis for downstream integrative comparisons.

#### Primary differential methylation analysis: Beta-Binomial modelling on a consensus CpG set

To address the statistical limitations of applying linear models to count-derived methylation proportions, the primary differential methylation analysis was performed using the DSS package(Version 2.54.0)[46], which implements a Beta-Binomial model with empirical Bayes shrinkage of dispersion estimates. Unlike approaches that treat *β*-values as continuous outcomes, DSS directly models the underlying read counts, methylated reads *M* and total coverage *N* , thereby accounting for the binomial sampling variance inherent to sequencing-based methylation measurements. This is particularly relevant in a multi-platform benchmarking context, where coverage depth varies substantially across methods and inflates the apparent precision of high-coverage platforms when proportion-based tests are used.

For array-based methylation profiling (EPIC), which does not yield read-count data, differential methylation was assessed using the limma package[39] with empirical Bayes moderation applied to M-values (logit transformed *β*-values). M-values transformation stabilizes the mean-variance relationship of array-derived methylation data, rendering it more suitable for linear modeling, rather than raw *β*-values. The statistical framework thus differed by necessity between sequencing-based (DSS, Beta-Binomial) and array-based (limma, linear model on M-values) platforms. This distinction is intrinsic to the measurement modality of each technology and was handled transparently throughout the analysis.

To ensure that inter-platform differences in DMC detection reflected methodological sensitivity rather than differential genomic coverage, all analyses were restricted to a consensus CpG set defined as sites fulfilling platform-specific data quality criteria in both tissue types simultaneously. For sequencing-based methods, a site was retained if it exhibited a minimum coverage of 10*x* in at least four of five samples per tissue (Blood and Fibroblast independently), a threshold established as a compromise between statistical reliability of the Beta-Binomial estimator and genomic breadth. For the EPIC array, the equivalent criterion required a valid, non-missing *β*-value in at least four of five samples per tissue. The consensus set was then defined as the intersection of all per-platform passing site sets across both tissues, yielding two analytical tiers:

- **Tier1** comprised CpG sites passing quality criteria across all four sequencing-based platforms (ONT, WGEC, RRBS and TWIST), yielding n=6,599 CpGs. This set constituted the primary analytical substrate for sequencing-based DMC detection and inter-platform concordance assessment.
- **Tier2** further intersected the Tier1 set with EPIC-covered sites, yielding n=6,598 CpGs and served as the substrate for direct assay-versus-sequencing comparison. Given that RRBS by design enriches for CpG island-proximal regions and that EPIC targets a curated subset of functionality relevant CpG loci, the Tier2 set is inherently enriched for promoter-associated and CpG island-resident sites. This genomic context bias is reported explicitly in the annotation analyses.

For all sequencing-based comparisons, smoothing was applied prior to testing using a Gaussian kernel with a bandwidth of 500bp (DSS::DMLtest, smoothing=TRUE), which pools information across neighbouring CpGs to improve power in regions of high local correlation. DMCs were called at FDR *<* 0.05 with a minimum absolute methylation difference of |Δ*β| ≥* 0.1. The latter serving as a biologically meaningful effect size filter independent of statistical significance. For EPIC, the equivalent filter was applied to the mean *β*-value difference between groups (Blood - Fibroblast), ensuring directional consistency with DSS-derived estimates.

#### Genomic annotation and region-level analysis

To contextualize DMC calls within functionally relevant genomic features, all significant CpGs were annotated using the annotatr package[6] against two complementary reference sets: CpG-structural features (CpG islands, shores, shelves and open sea regions) and gene-centric features (promoters, 5’UTRs, exons, introns, 3’UTRs and intergenic regions), both derived from the hg38 reference assembly. Annotation proportions were compared across platforms to assess whether observed inter-method differences in DMC counts reflected genuine methylation biology or were confounded by platform-specific genomic coverage biases, in particular the known enrichment of RRBS for CpG island-proximal loci and the targeted nature of the EPIC array design.

To assess concordance at the level of genomic regions rather than individual CpG sites, which is more robust to local coverage gaps and to the stochastic nature of single-CpG detection, differential methylation region (DMR) analysis was performed using DMRcate(Version 3.2.1)[35]. DMRcate integrates CpG-level test statistics from the preceding DSS(sequencing) and limma(EPIC) analyses via kernel smoothing, aggregating spatially proximal significant CpGs into contiguous DMRs. A Gaussian kernel bandwidth of *λ* = 1,000 bp was applied with a minimum inter-CpG spacing of *C* = 2 and a minimum of three CpGs per DMR, parameters chosen to balance sensitivity against the CpG density constraints imposed by the consensus site set. Inter-platform concordance was quantified using a base-pair-weighted Jaccard similarity index, defined as the ratio of the total intersecting base pairs to the total union of base pairs across all pairwise DMR set comparisons. This metric penalises boundary discrepancies between platforms while rewarding overlap in the core methylated region, and is therefore more informative than simple DMR count overlap for benchmarking purposes.

Overlap of significant DMCs and DMRs across platforms was visualized using UpSet plots (ComplexUpset)[22], with intersection bars and set membership indicators color-coded by platform identity and assay-type background shading (Long-read sequencing: ONT; Short-read sequencing: WGEC, TWIST, RRBS; Array: EPIC) to facilitate visual stratification by technological category. Pairwise platform concordance was additionally summarized using Jaccard similarity heatmaps at both the CpG and DMR levels.

## Abbreviations

5hmC: 5-Hydroxymethylcytosine
5mC: 5-Methylcytosine
6mA: N6-Methyladenine
ATAC-seq: Assay for Transposase-Accessible Chromatin sequencing
BAM: Binary Alignment Map
CDS: Coding Sequence
CGI: CpG Island
CpG: Cytosine-phosphate-Guanine dinucleotide
DAC: Data Access Committee
DFG: Deutsche Forschungsgemeinschaft
DMC: Differentially Methylated Cytosine
DNA: Deoxyribonucleic Acid
EGA: European Genome-phenome Archive
EPIC: Infinium MethylationEPIC BeadChip
FASTQ: Fast All quality
FDR: False Discovery Rate
GIAB: Genome in a Bottle
HMW: High Molecular Weight
LR-GS: Long-Read Genome Sequencing
NGS: Next Generation Sequencing
ONT: Oxford Nanopore Technologies
PacBio: Pacific Biosciences
PCA: Principal Component Analysis
PBMC: Peripheral Blood Mononuclear Cell
RNA-seq: RNA Sequencing
RRBS: Reduced Representation Bisulfite Sequencing
SMRT: Single-Molecule Real-Time
SNP: Single Nucleotide Polymorphism
TE: Transposable Element
TSS: Transcription Start Site
TWIST: Twist Targeted Methylation Panel
UTR: Untranslated Region
WGBS: Whole-Genome Bisulfite Sequencing
WGEC: Whole-Genome Enzymatic Conversion

## Acknowledgments

This project was funded by the DFG (German Research Foundation) through a grant to JSH (SCHU 2693/7-1) and RB (BU 3602/1-1).

OR was funded by the Ministry of Science, Research and Art of Baden-Württemberg through the BEGIN project (MWK25-0123-139/4/1), and through joint funding of the “Genomes of Europe” project by the Federal Ministry of Research, Technology and Space of Germany and the European Commission.

NGS methods were performed with the support of the DFG-funded NGS Competence Center Tubingen (INST 37/1049-1), funded by the Deutsche Forschungsgemeinschaft (DFG, German Research Foundation) – Project-ID 286/2020B01-428994620.

## Author Contributions

J.S.-H., R.B., S.O., T.H. and O.R. conceived the study. J.S.-H., T.H. and R.B. designed the study. L.L. was responsible for formal analysis, data interpretation, and writing the original draft. G.G. performed preprocessing and quality control of the EPIC array data. T.H. conducted preprocessing and quality control of the RRBS data. J.A., N.C., and T.B.H. performed sequencing and preprocessing of ONT, WGEC, and TWIST data. E.B.-A., M.P., L.S. and R.B. carried out the experimental laboratory procedures, including DNA extraction and library preparation. L.L., E.B.-A., M.P. and R.B. contributed to writing, and all authors contributed to revising the manuscript.

## Data Availability

GIAB reference samples (HG001, HG002) including PacBio methylation data are publicly available via the PacBio website (HG002 SPRQ-Nx dataset: https://downloads.pacbcloud.com/public/2026Q2/HG002-SPRQ- Nx/; HG001 Platinum Pedigree dataset: see [19][34]) Processed methylation matrices are available from the corresponding author upon reasonable request.

## Code Availability

The respective code is available in the Github repo: https://github.com/epigenetics-sb/MethylBench.

## Ethics Statement

Informed consent was obtained for all probands and studies were conducted in accordance with the local ethics committee (066/2021BO2).

## Supplementary Figures

**Figure S1:**
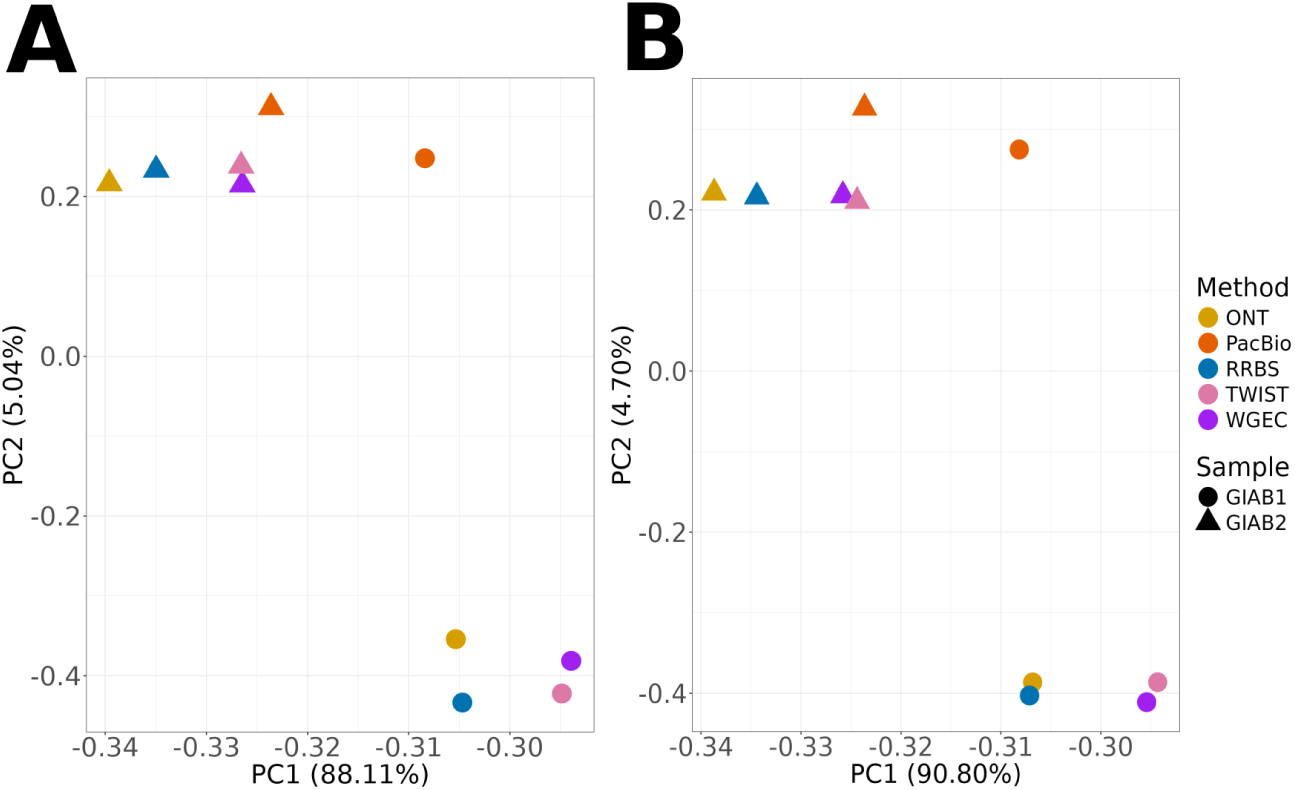
Principal component analysis of genome-wide methylation profiles across quantification methods for GIAB samples. PCA of CpG methylation profiles from ONT, PacBio, RRBS, TWIST, and WGEC before **(A)** and after **(B)** 10x coverage filtering. PC1 explained most variance (88.1%/90.8%) and primarily separated the two GIAB donors rather than methods. PacBio was the exception: both donor profiles clustered together and remained distinct from the other methods before and after filtering, indicating systematic method-specific variation beyond coverage effects.

**Figure S2:**
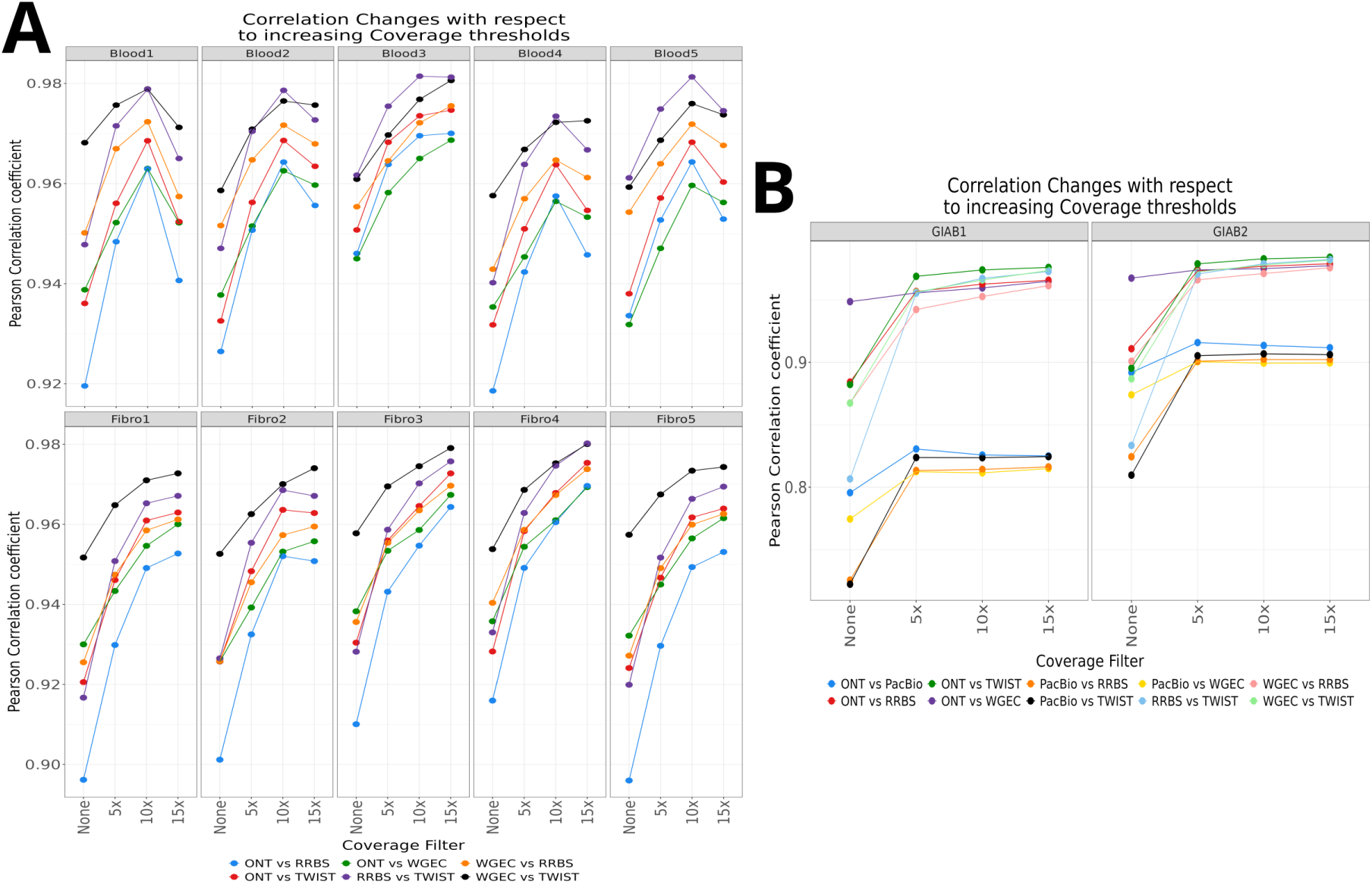
Pairwise methylation correlation across methods at increasing coverage thresholds for five blood, fibroblast and two GIAB samples. (A): Pearson correlation coefficients for blood & fibroblast samples. **(B):** Pearson correlation coefficients for GIAB samples. The lack of increased correlation in blood and fibroblast samples likely results from the substantial reduction of the CpG space under stringent coverage filtering (see Figure 3C), which limits correlation robustness.

**Figure S3:**
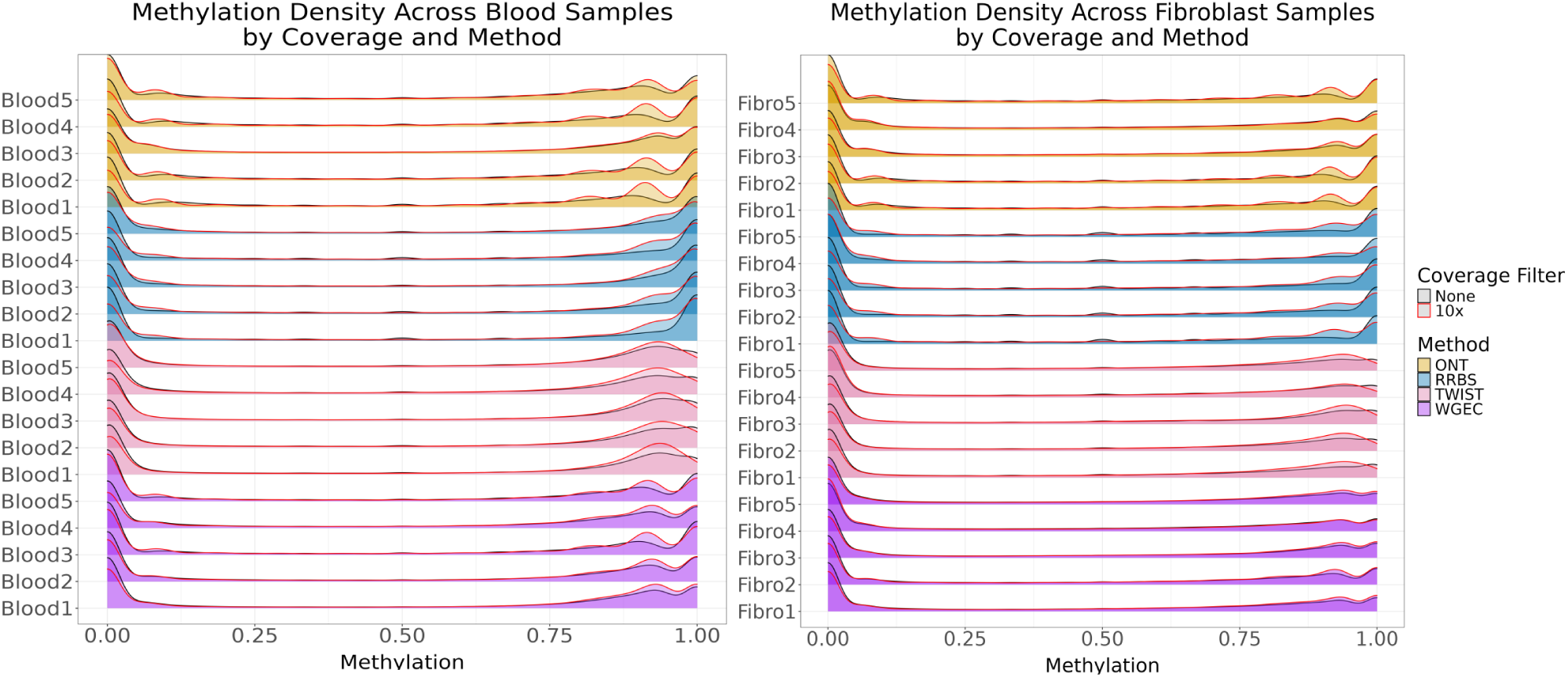
Methylation density distributions across samples, methods and coverage thresholds for five blood and fibroblast samples.(Left): Blood samples. **(Right):** Fibroblast samples.

**Figure S4:**
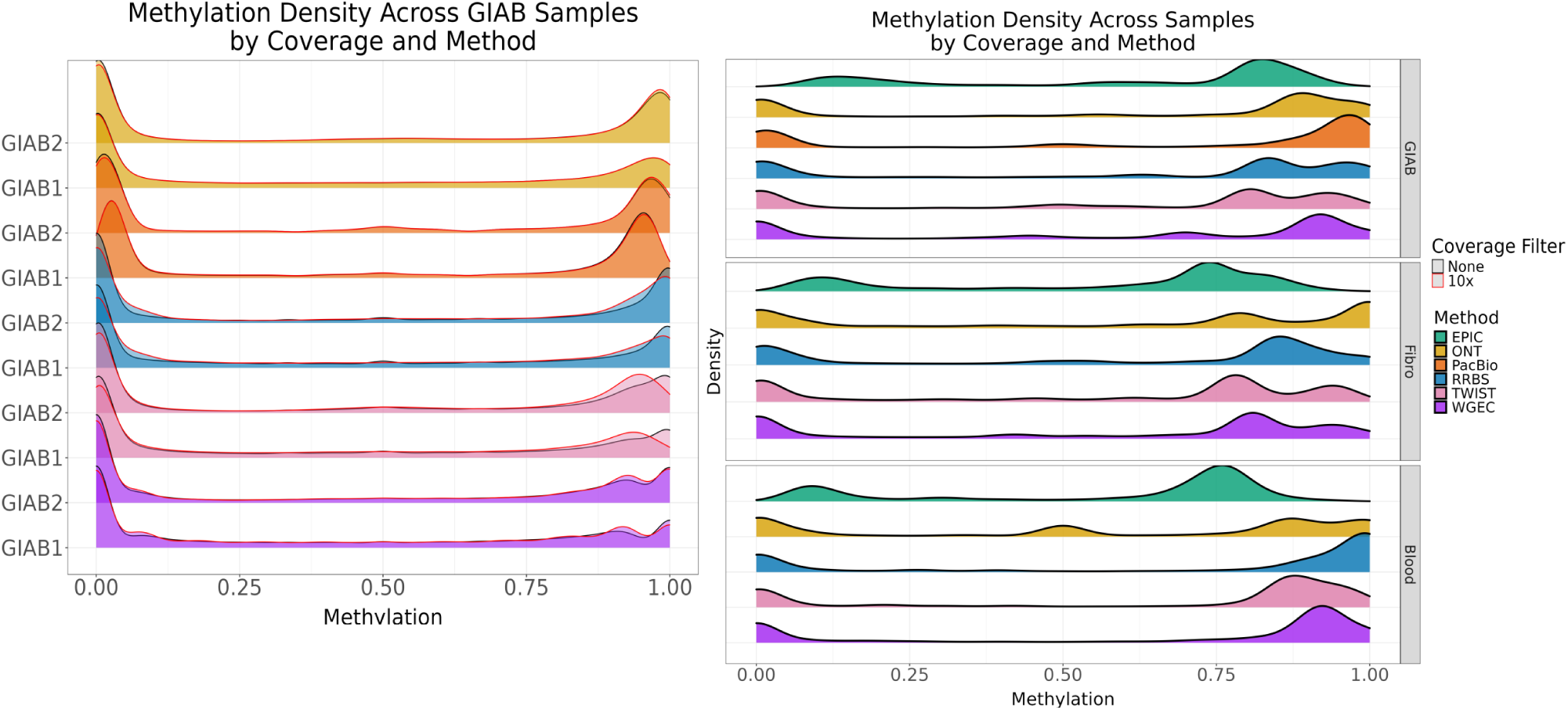
Methylation density distributions across samples, methods and coverage thresholds. (Left): Methylation density profiles for two GIAB samples. **(Right):** Methylation density distributions across samples and methods for the samples Blood3, Fibro4 and GIAB1, with EPIC included. The coverage filter was set to 10x for all methods and then overlapped with the EPIC data. The pronounced intermediate methylation peak observed exclusively for ONT in blood likely reflects its single-molecule measurement principle combined with the high cellular heterogeneity of blood, resulting in genuine intermediate methylation states that are attenuated or discretized by aggregation-based array and bisulfite sequencing approaches.

**Figure S5:**
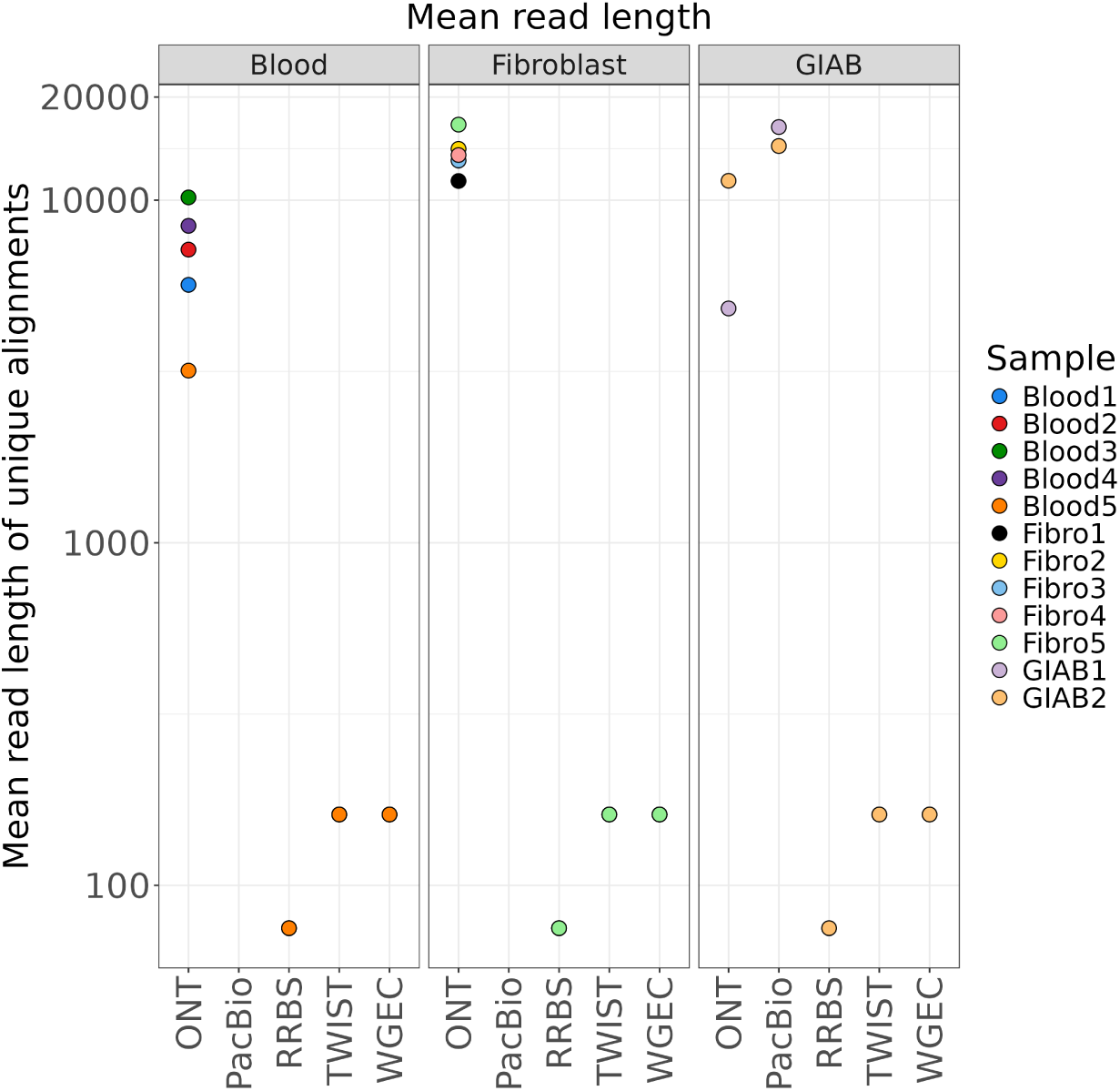
Mean read length of uniquely aligned reads, highlighting differences in read size distribution across technologies. For TWIST, WGEC and RRBS read lengths are fixed by library construction and sequencing design, resulting in complete overlap of values and a single visible data point per sample set.

**Figure S6:**
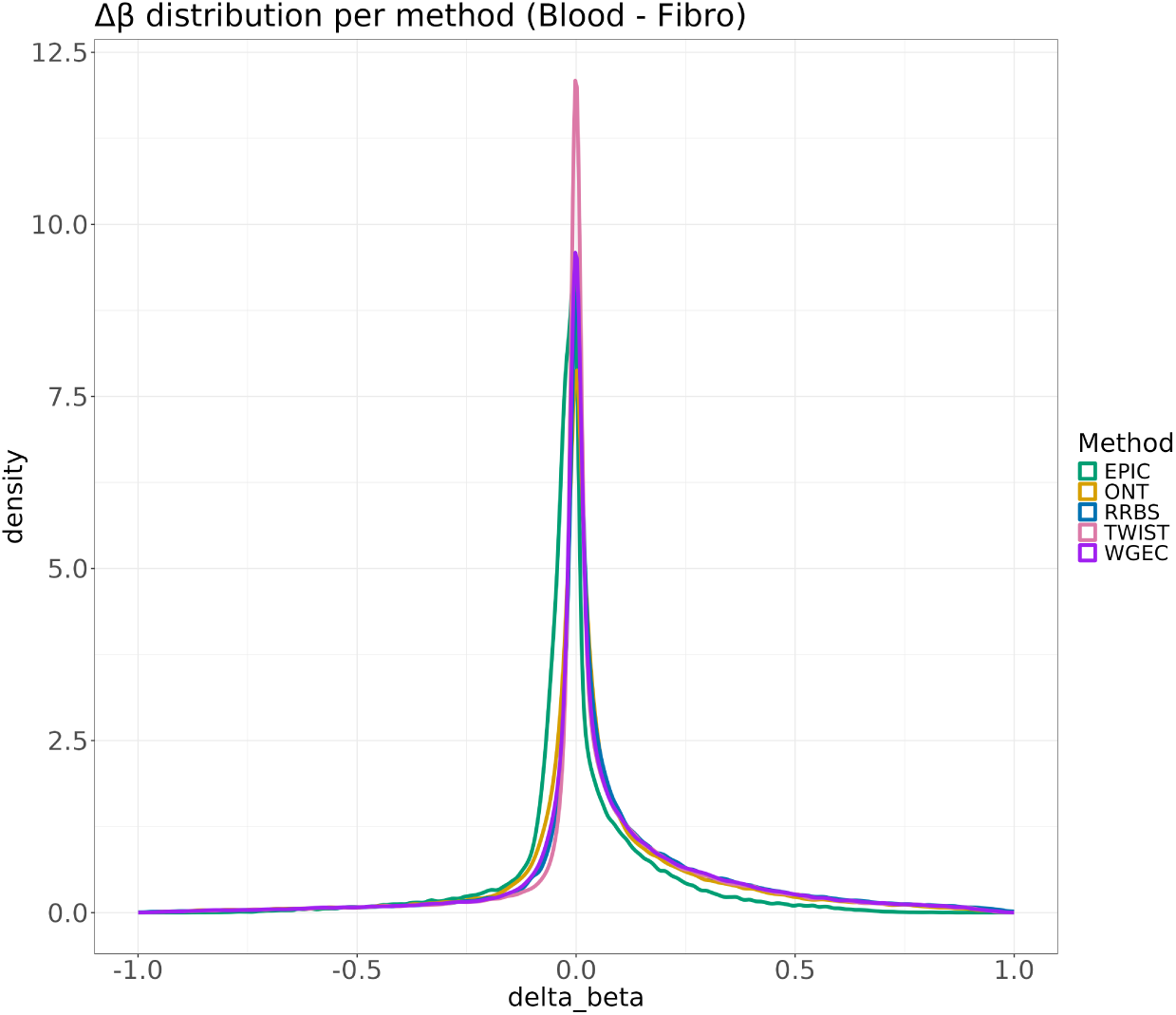
Platform-specific. Δ*β* distribution per method (Fibroblast vs. Blood).

**Figure S7:**
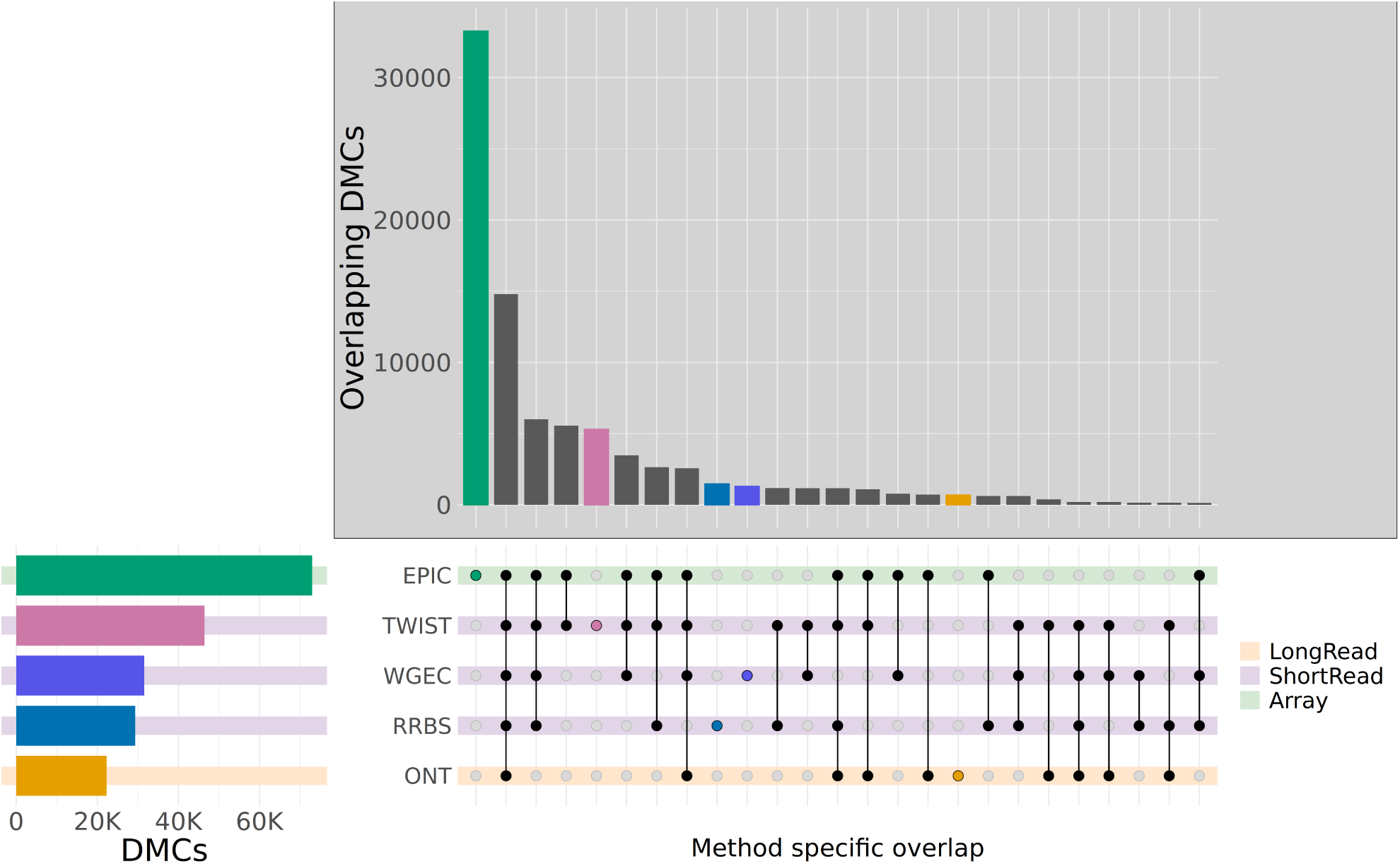
**UpSet plot showing the overlap of significantly differentially methylated CpG sites (DMCs) between blood and fibroblast samples across EPIC, TWIST, WGEC, RRBS and ONT datasets, as identified using the limma model.**

**Figure S8:**
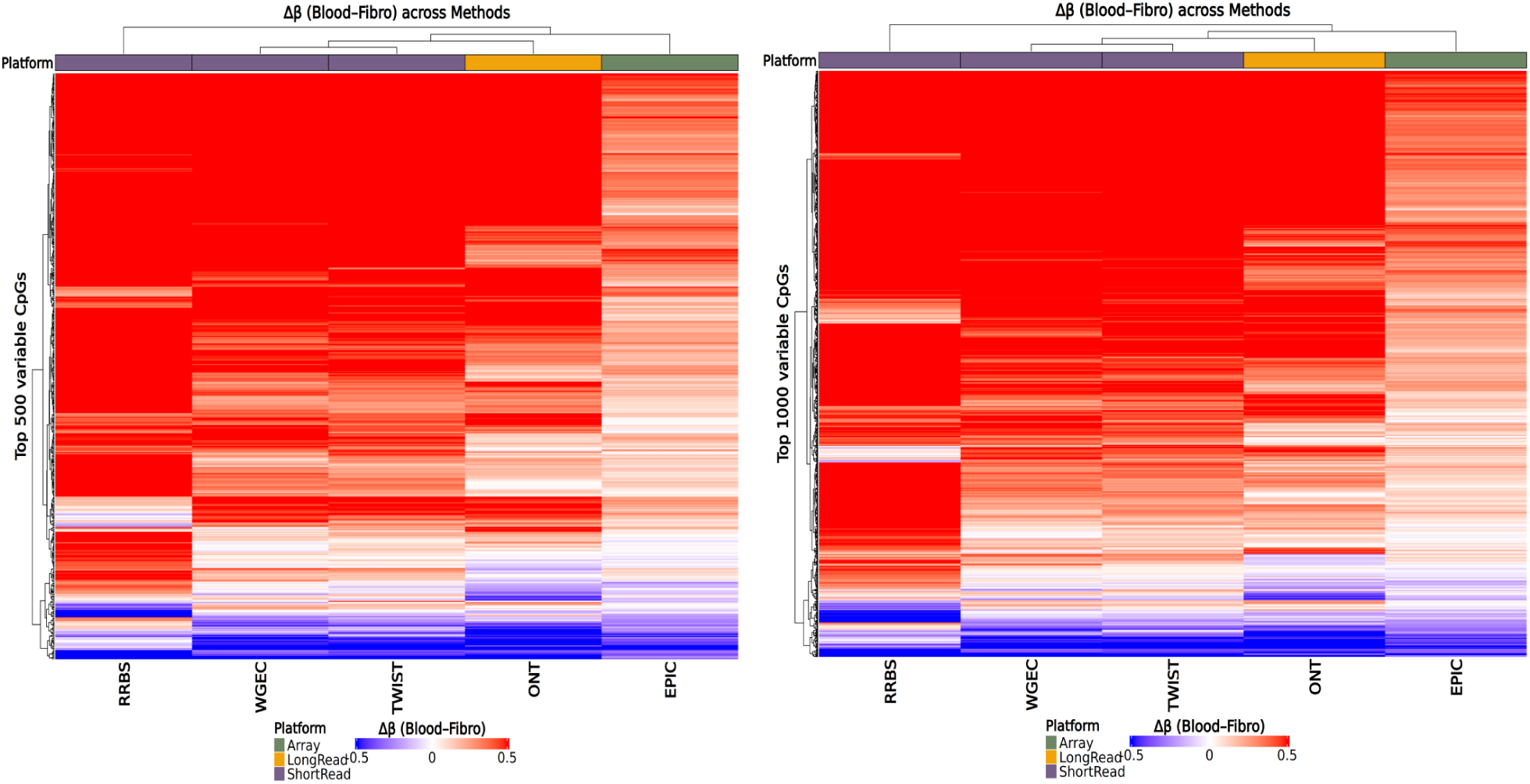
Heatmaps of the most variable CpGs. (Left): Heatmap of the top 500 most variable CpGs, based on Δ*β* value and ranked according to p-value. **(Right):** Heatmap of the top 1000 most variable CpGs, based on Δ*β* value and ranked according to p-value.

**Figure S9:**
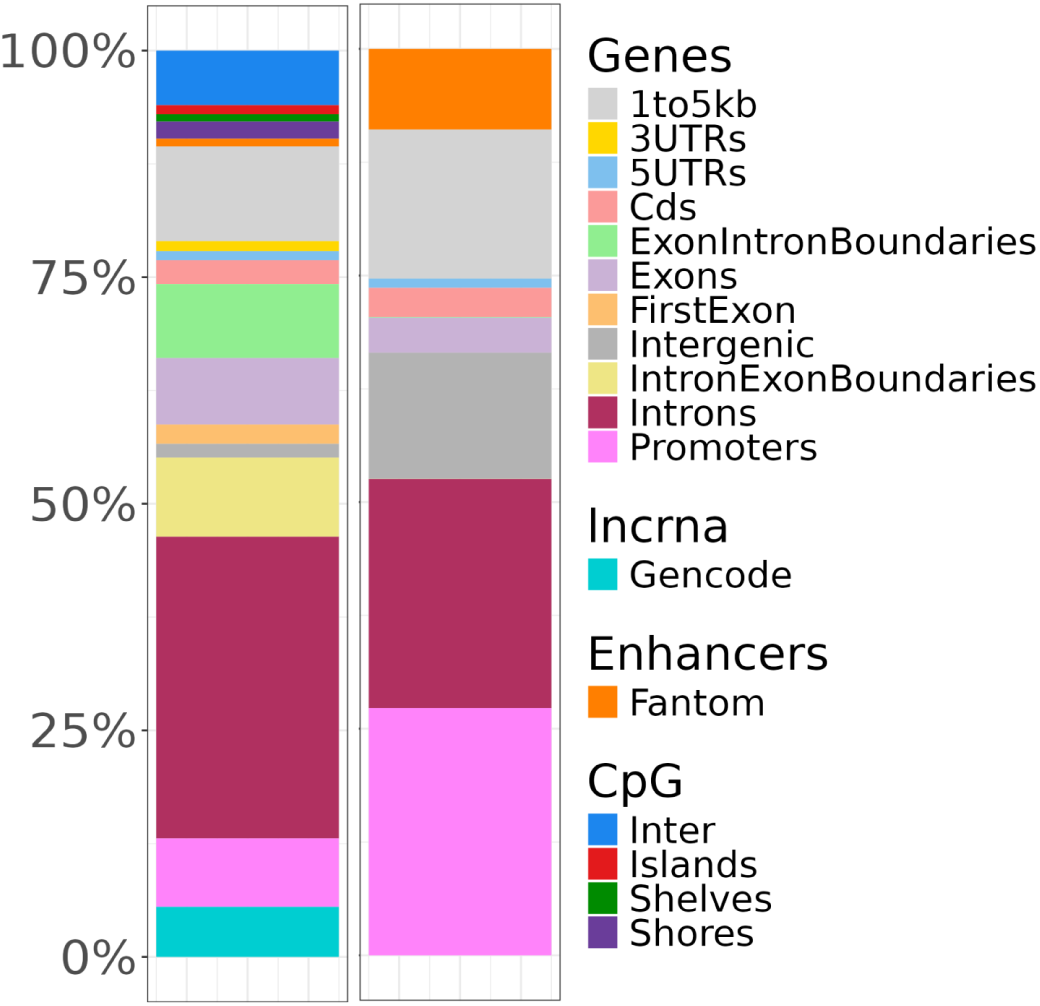
Annotation of the significant identified DMCs(left) and reduced annotation distribution, after reducing DMC annotation space to unique annotations(right). All annotations are derived from the R-package `annotatr`[6].

**Figure S10:**
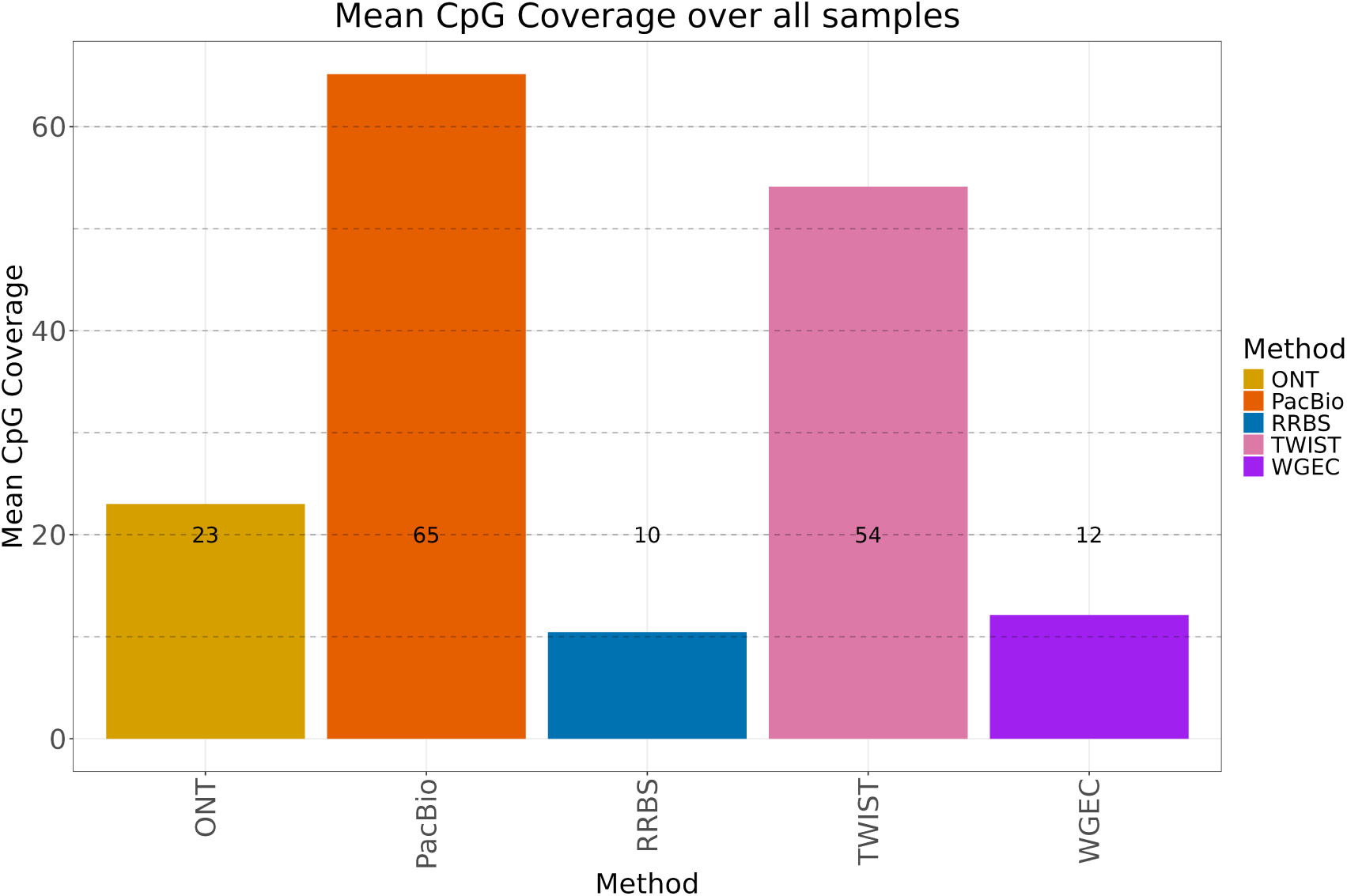
Barplot showing the mean genomic coverage per method over all samples. Dashed lines indicate multiples of a 10× coverage threshold & numbers represent the rounded mean genomic coverage. For ONT, RRBS, WGEC & TWIST, *n_Samples_* = 12 & PacBio *n_Samples_* = 2.

**Figure S11:**
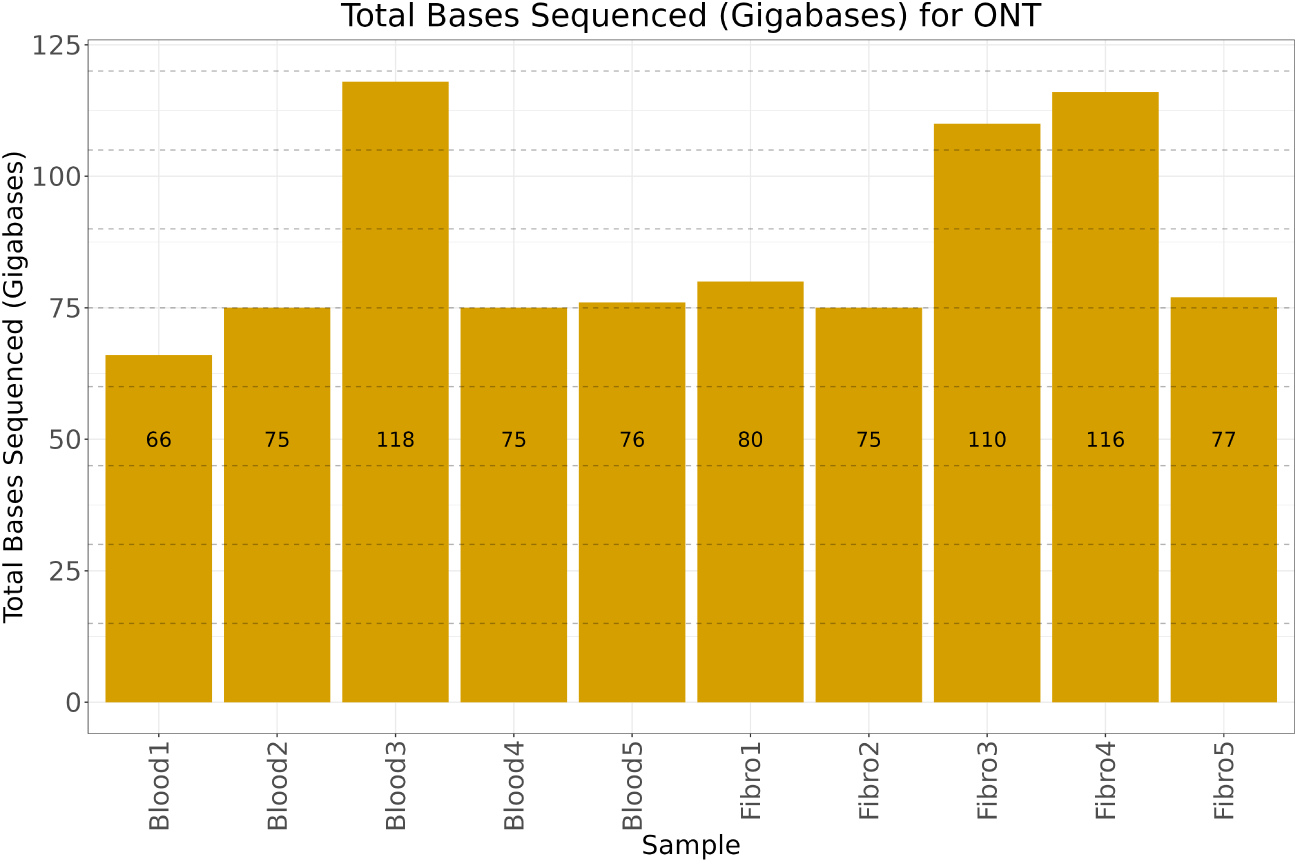
Barplot showing the total bases sequenced per sample, expressed in gigabases (Gb) for ONT. Dashed lines indicate multiples of a 15× threshold & numbers represent the rounded total bases sequenced in Gb.

**Figure S12:**
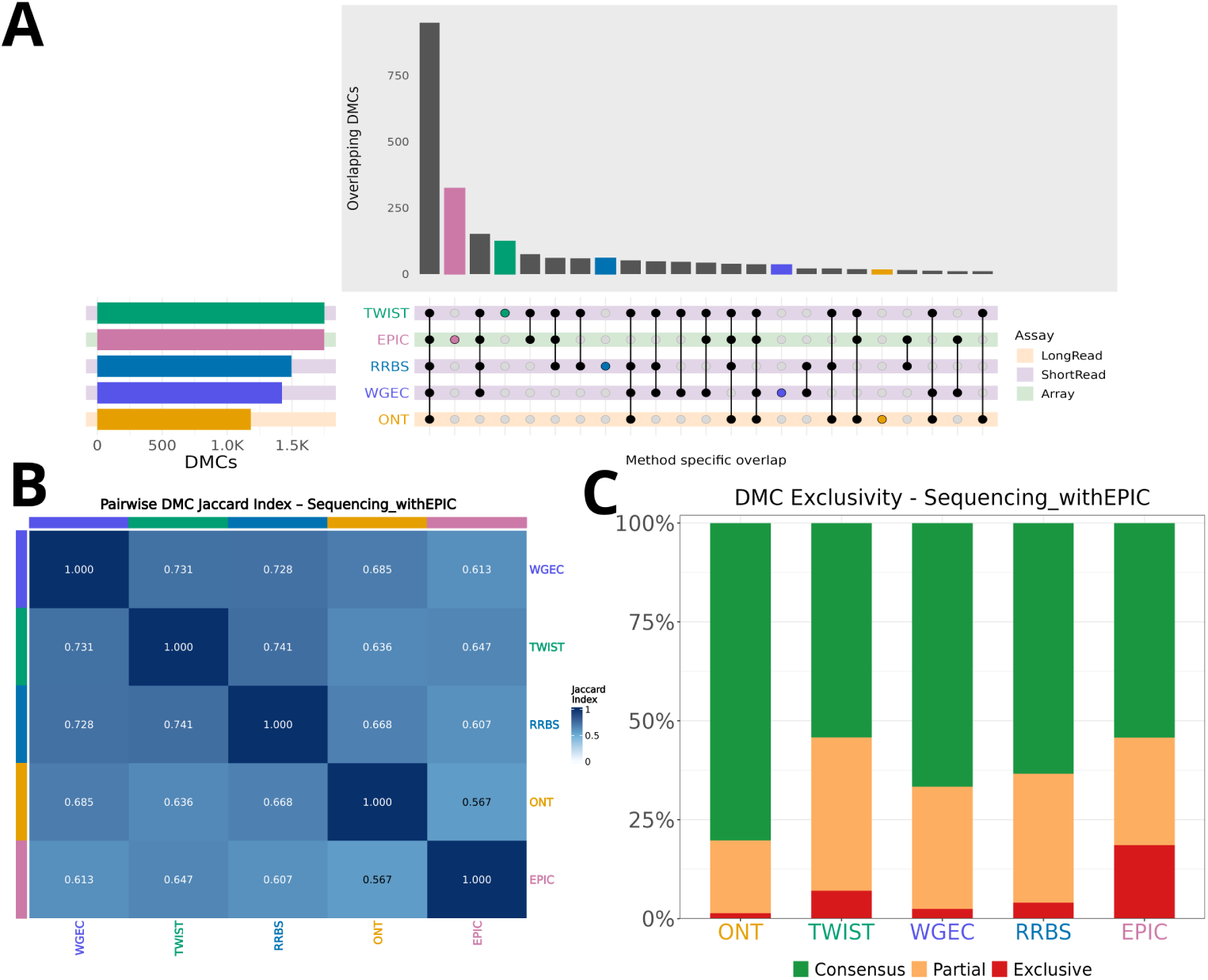
Differential methylation concordance across all five platforms, including EPIC, at the CpG level (Tier 2 consensus set). Analyses were performed on the Tier 2 consensus CpG set (n = 6,598 CpGs; EPIC DMCs called via limma on M-values, see Methods). **(A):** UpSet plot of significant DMC overlap across TWIST, EPIC, RRBS, WGEC and ONT; intersection bars are colored by assay category (long-read, short-read, array) as in Figure 8B. **(B):** Pairwise DMC Jaccard similarity index across all five platforms. **(C):** DMC exclusivity per platform, showing the proportion of each platform’s DMCs detected by all five methods (Consensus), a subset of methods (Partial) or exclusively by one method (Exclusive).

**Figure S13:**
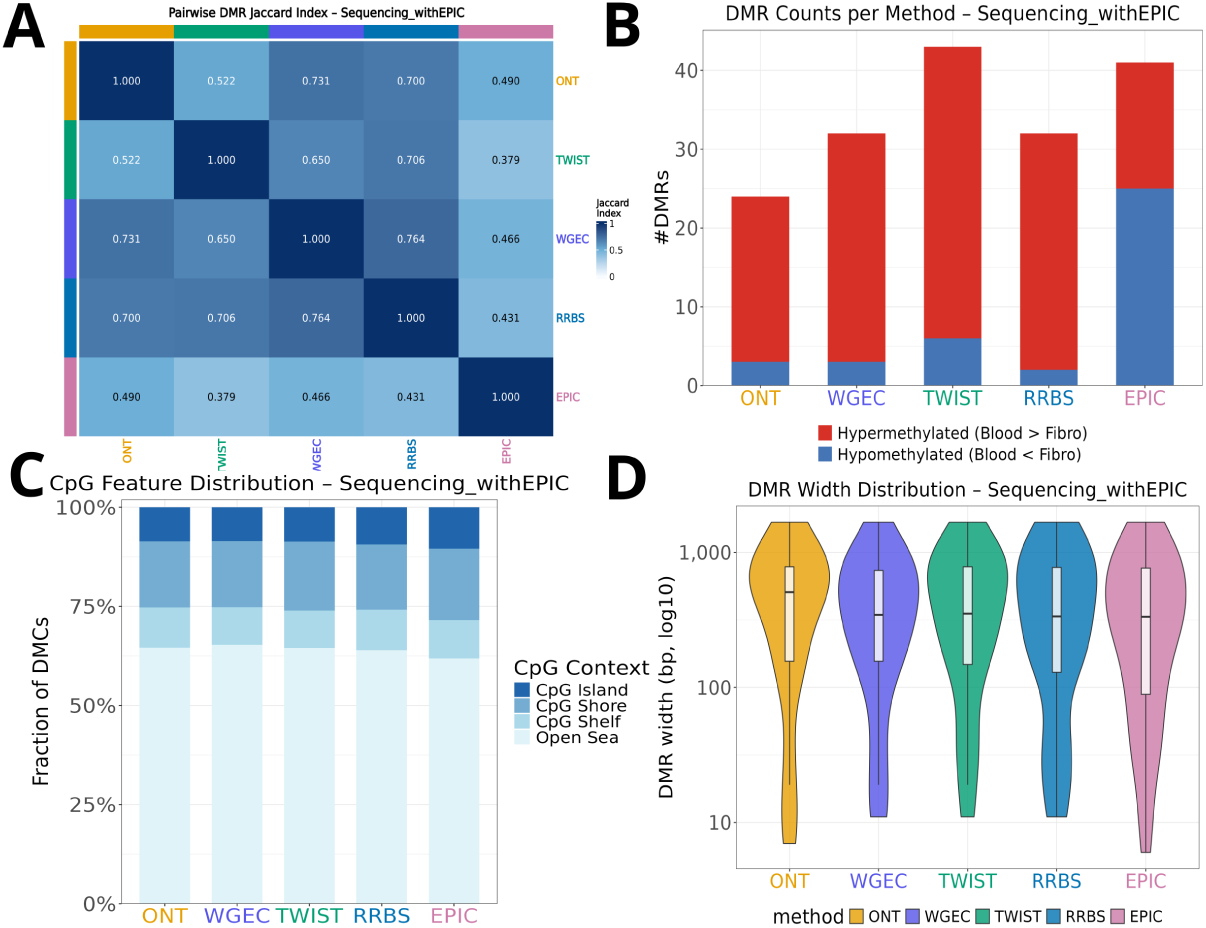
Differential methylation region (DMR) concordance across all five platforms, including EPIC (Tier 2 consensus set). DMRs for EPIC were identified using DMRcate applied to limma-derived CpG-level statistics on M-values, using the same smoothing and minimum-CpG parameters as for the sequencing-based platforms (see Methods and Figure 9). **(A):** Pairwise base-pair-weighted Jaccard similarity index between DMR sets across all five platforms. **(B):** Number of DMRs identified per platform, stratified by direction of differential methylation (red: hypermethylated in blood; blue: hypomethylated in blood). **(C):** Genomic CpG context distribution (CpG islands, shores, shelves, open sea) of DMRs per platform. **(D):** Distribution of DMR widths per platform (log10 scale).

**Figure S14:**
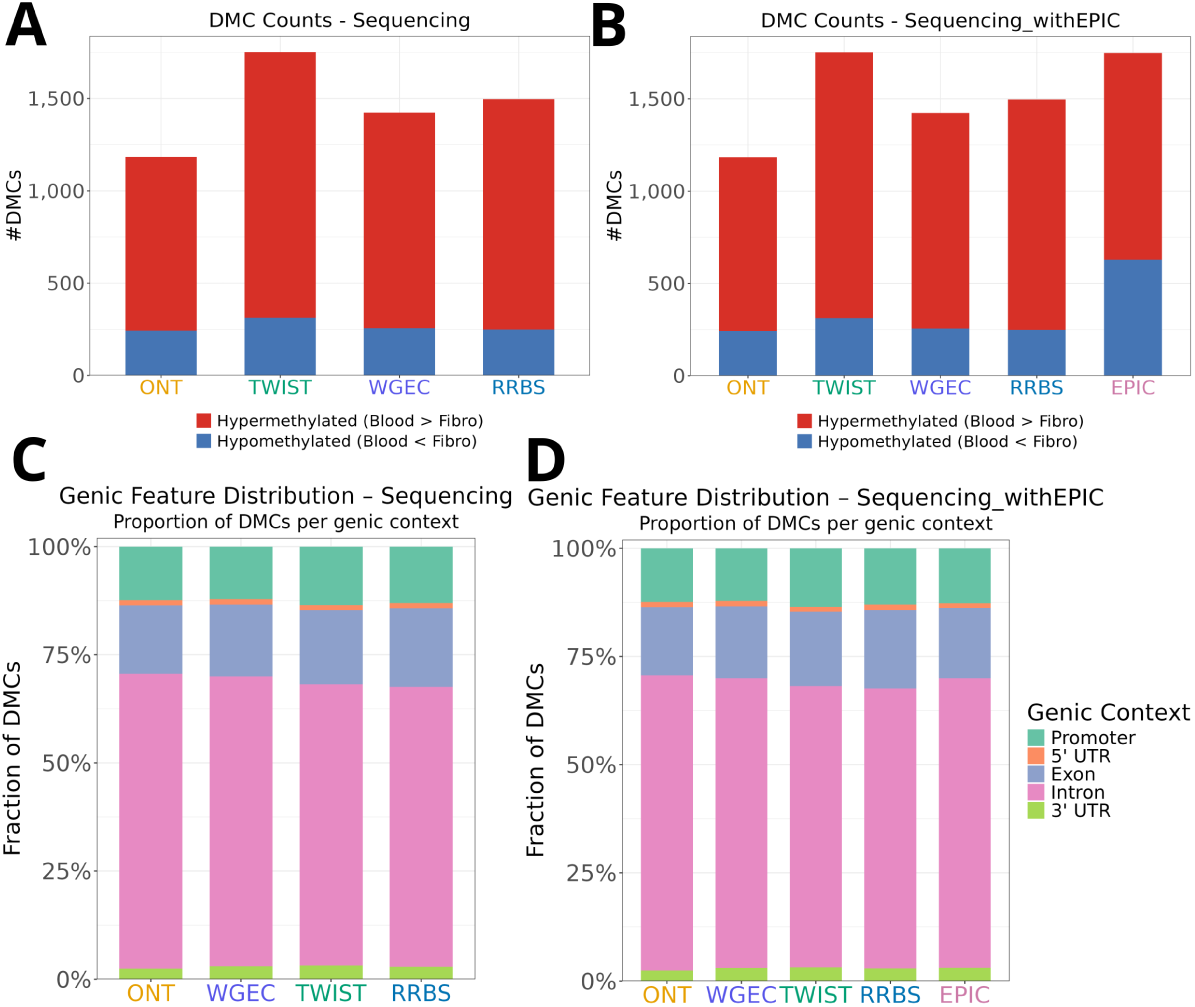
Direction and genic context of differentially methylated cytosines (DMCs) across platforms. (A): Number of significant DMCs per platform for the four sequencing-based technologies (Tier 1 consensus set), stratified by direction of methylation change. **(B):** As in (A), including EPIC (Tier 2 consensus set). **(C):** Proportion of DMCs per platform annotated to genic features (promoter, 5’ UTR, exon, intron, 3’ UTR) for the four sequencing-based technologies. **(D):** As in (C), including EPIC. Gene-centric annotations were derived using annotatr (hg38 reference), as in Figure S9.

**Figure S15:**
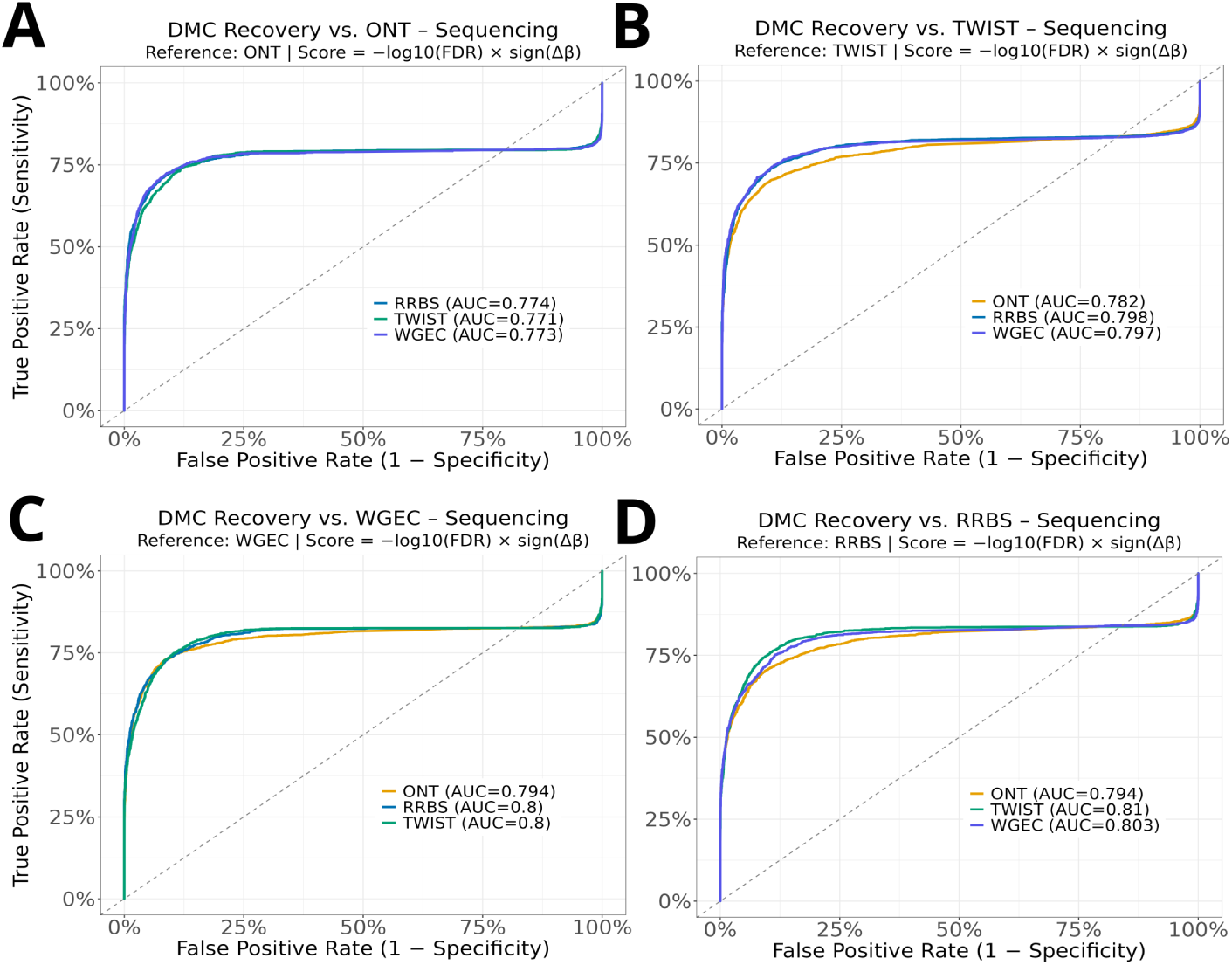
Threshold-free recovery of platform-specific DMCs across sequencing technologies. For each of the four sequencing-based platforms (ONT, TWIST, WGEC, RRBS) as reference, ROC curves show how well the ranked score (*−log*10(*FDR*) x *sign*(Δ*β*)) from each of the remaining three platforms recovers the reference platform’s significant DMCs (Tier 1 consensus set). **(A):** Reference: ONT. **(B):** Reference: TWIST. **(C):** Reference: WGEC. **(D):** Reference: RRBS. AUC values are given per comparison.

**Table S1:**
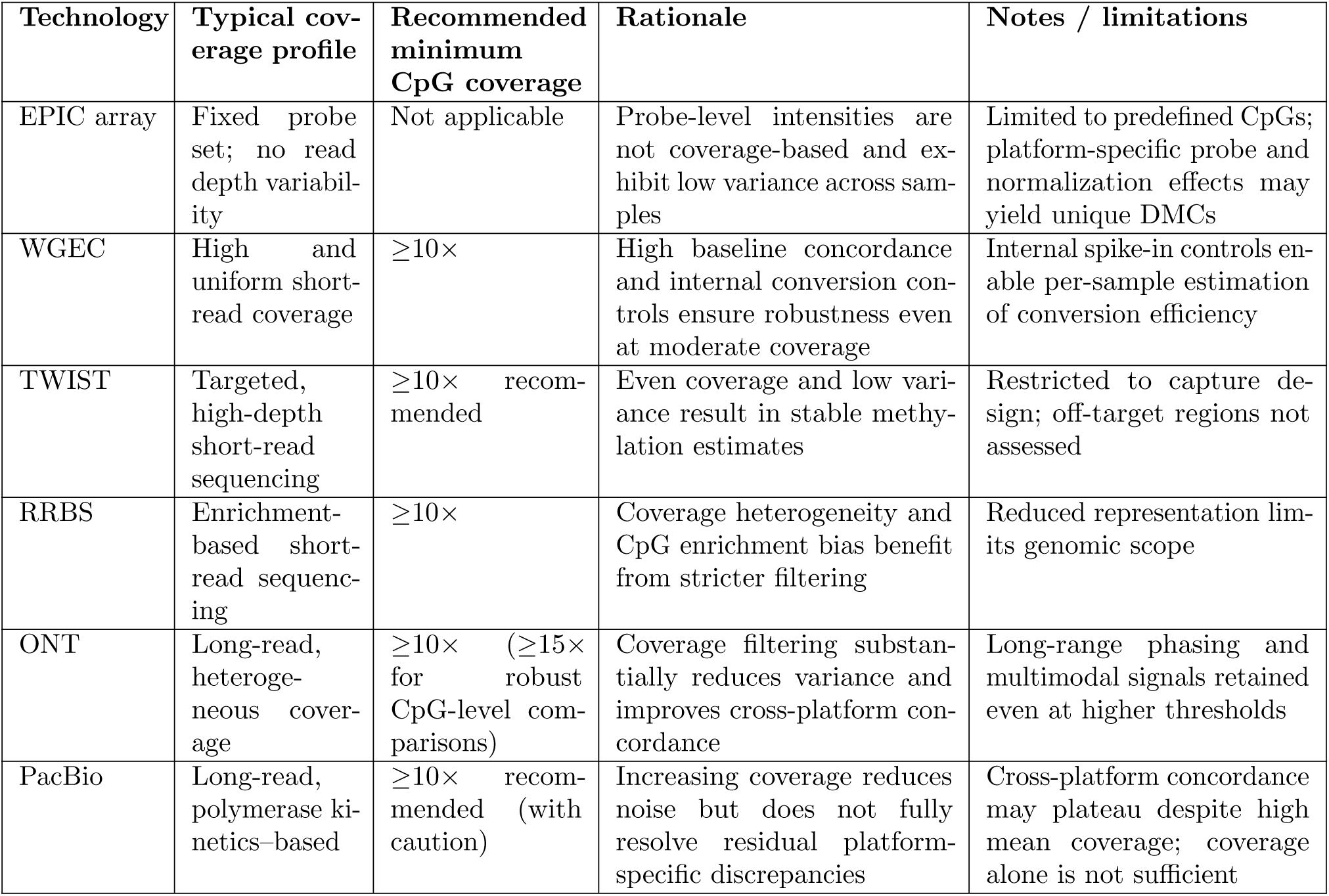
Recommended Minimum CpG Coverage Thresholds for Integrative DNA Methylation Analyses Based on Variance, Cross-Platform Concordance, and CpG Retention. Recommended minimum CpG coverage thresholds derived from empirical evaluation of coverage-dependent variance, cross-platform correlation, and CpG retention across six DNA methylation profiling technologies. Thresholds represent pragmatic guidelines for integrative analyses rather than universal optima and should be adapted to study design, sample quality and analytical objectives.

## References

[1] Y. Assenov, F. Müller, P. Lutsik, J. Walter, T. Lengauer, and C. Bock. Comprehensive analysis of dna methylation data with rnbeads. Nature methods, 11(11):1138–1140, 2014.

[2] T. Barrett, M. Dowle, A. Srinivasan, J. Gorecki, M. Chirico, T. Hocking, B. Schwendinger, and I. Krylov. data.table: Extension of ‘data.frame*‘*, 2025. R package version 1.17.99.

[3] S. B. Baylin and P. A. Jones. A decade of exploring the cancer epigenome—biological and translational implications. Nature Reviews Cancer, 11(10):726–734, 2011.

[4] J. Beltran-Garcıa, R. Osca-Verdegal, S. Mena-Molla, and J. L. García-Giménez. Epigenetic ivd tests for personalized precision medicine in cancer. Frontiers in genetics, 10:621, 2019.

[5] A. Breiling and F. Lyko. Epigenetic regulatory functions of dna modifications: 5-methylcytosine and beyond. Epigenetics & chromatin, 8:1–9, 2015.

[6] R. G. Cavalcante and M. A. Sartor. Annotatr: genomic regions in context. Bioinformatics, 33(15):2381–2383, 2017.

[7] J. Clarke, H.-C. Wu, L. Jayasinghe, A. Patel, S. Reid, and H. Bayley. Continuous base identification for single-molecule nanopore dna sequencing. Nature nanotechnology, 4(4):265–270, 2009.

[8] J. R. Conway, A. Lex, and N. Gehlenborg. Upsetr: an r package for the visualization of intersecting sets and their properties. Bioinformatics, 33(18):2938–2940, 2017.

[9] P. M. d. S. Costa, S. L. A. Sales, D. P. Pinheiro, L. Q. Pontes, S. S. Maranhao, C. d. Ó. Pessoa, G. P. Furtado, and C. L. M. Furtado. Epigenetic reprogramming in cancer: From diagnosis to treatment. Frontiers in Cell and Developmental Biology, 11:1116805, 2023.

[10] L. Di Stefano. All quiet on the te front? the role of chromatin in transposable element silencing. Cells, 11(16):2501, 2022.

[11] A. R. Elhamamsy. Role of dna methylation in imprinting disorders: an updated review. Journal of assisted reproduction and genetics, 34(5):549–562, 2017.

[12] epi2me labs/modbam2bed. online, Nov. 2024.

[13] B. A. Flusberg, D. R. Webster, J. H. Lee, K. J. Travers, E. C. Olivares, T. A. Clark, J. Korlach, and S. W. Turner. Direct detection of dna methylation during single-molecule, real-time sequencing. Nature methods, 7(6):461–465, 2010.

[14] J. Foox, J. Nordlund, C. Lalancette, T. Gong, M. Lacey, S. Lent, B. W. Langhorst, V. K. C. Ponnaluri, L. Williams, K. R. Padmanabhan, R. Cavalcante, A. Lundmark, D. Butler, C. Mozsary, J. Gurvitch, J. M. Greally, M. Suzuki, M. Menor, M. Nasu, A. Alonso, C. Sheridan, A. Scherer, S. Bruinsma, G. Golda, A. Muszynska, P. P. Labaj, M. A. Campbell, F. Wos, A. Raine, U. Liljedahl, T. Axelsson, C. Wang, Z. Chen, Z. Yang, J. Li, X. Yang, H. Wang, A. Melnick, S. Guo, A. Blume, V. Franke, I. Ibanez de Caceres, C. Rodriguez-Antolin, R. Rosas, J. W. Davis, J. Ishii, D. B. Megherbi, W. Xiao, W. Liao, J. Xu, H. Hong, B. Ning, W. Tong, A. Akalin, Y. Wang, Y. Deng, and C. E. Mason. The seqc2 epigenomics quality control (epiqc) study. Genome Biology, 22(1), Dec. 2021.

[15] H. Gu, C. Bock, T. S. Mikkelsen, N. Jager, Z. D. Smith, E. Tomazou, A. Gnirke, E. S. Lander, and A. Meissner. Genome-scale dna methylation mapping of clinical samples at single-nucleotide resolution. Nature methods, 7(2):133–136, 2010.

[16] M. Gutierrez-Arcelus, H. Ongen, T. Lappalainen, S. B. Montgomery, A. Buil, A. Yurovsky, J. Bryois, I. Padioleau, L. Romano, A. Planchon, et al. Tissue-specific effects of genetic and epigenetic variation on gene regulation and splicing. PLoS genetics, 11(1):e1004958, 2015.

[17] J. D. Hollister and B. S. Gaut. Epigenetic silencing of transposable elements: a trade-off between reduced transposition and deleterious effects on neighboring gene expression. Genome research, 19(8):1419–1428, 2009.

[18] G. in a bottle consortium. Genome in a bottle - a human dna standard. Nature Biotechnology, 33(7):675–675, July 2015.

[19] Z. Kronenberg, C. Nolan, D. Porubsky, T. Mokveld, W. J. Rowell, S. Lee, E. Dolzhenko, P.-C. Chang, J. M. Holt, C. T. Saunders, N. D. Olson, C. J. Steely, S. McGee, A. Guarracino, N. Koundinya, W. T. Harvey, W. S. Watkins, K. M. Munson, K. Hoekzema, K. P. Chua, X. Chen, C. Fanslow, C. Lambert, H. Dashnow, E. Garrison, J. D. Smith, P. M. Lansdorp, J. M. Zook, A. Carroll, L. B. Jorde, D. W. Neklason, A. R. Quinlan, E. E. Eichler, and M. A. Eberle. The platinum pedigree: a long-read benchmark for genetic variants. Nature Methods, 22(8):1669–1676, Aug. 2025.

[20] Z. Kronenberg, C. Nolan, D. Porubsky, T. Mokveld, W. J. Rowell, S. Lee, E. Dolzhenko, P.-C. Chang, J. M. Holt, C. T. Saunders, N. D. Olson, C. J. Steely, S. McGee, A. Guarracino, N. Koundinya, W. T. Harvey, W. S. Watkins, K. M. Munson, K. Hoekzema, K. P. Chua, X. Chen, C. Fanslow, C. Lambert, H. Dashnow, E. Garrison, J. D. Smith, P. M. Lansdorp, J. M. Zook, A. Carroll, L. B. Jorde, D. W. Neklason, A. R. Quinlan, E. E. Eichler, and M. A. Eberle. Platinum-pedigree-datasets. online, 2025.

[21] F. Krueger and S. R. Andrews. Bismark: a flexible aligner and methylation caller for bisulfite-seq applications. *bioinformatics*, 27(11):1571–1572, 2011.

[22] A. Lex, N. Gehlenborg, H. Strobelt, R. Vuillemot, and H. Pfister. Upset: visualization of intersecting sets. IEEE transactions on visualization and computer graphics, 20(12):1983–1992, 2014.

[23] R. Lister, M. Pelizzola, R. H. Dowen, R. D. Hawkins, G. Hon, J. Tonti-Filippini, J. R. Nery, L. Lee, Z. Ye, Q.-M. Ngo, et al. Human dna methylomes at base resolution show widespread epigenomic differences. *nature*, 462(7271):315–322, 2009.

[24] N. Loyfer, J. Magenheim, A. Peretz, G. Cann, J. Bredno, A. Klochendler, I. Fox-Fisher, S. Shabi-Porat, M. Hecht, T. Pelet, et al. A dna methylation atlas of normal human cell types. Nature, 613(7943):355–364, 2023.

[25] M. Martin, M. Patterson, S. Garg, S. O Fischer, N. Pisanti, G. W. Klau, A. Schoenhuth, and T. Marschall. Whatshap: fast and accurate read-based phasing. BioRxiv, page 085050, 2016.

[26] A. Meissner, A. Gnirke, G. W. Bell, B. Ramsahoye, E. S. Lander, and R. Jaenisch. Reduced representation bisulfite sequencing for comparative high-resolution dna methylation analysis. Nucleic acids research, 33(18):5868–5877, 2005.

[27] H. Meng, Y. Cao, J. Qin, X. Song, Q. Zhang, Y. Shi, and L. Cao. Dna methylation, its mediators and genome integrity. International journal of biological sciences, 11(5):604, 2015.

[28] K. Mohan, G. Gasparoni, A. Salhab, M. M. Orlich, R. Geffers, S. Hoffmann, R. H. Adams, J. Walter, and A. Nordheim. Age-associated changes in endothelial transcriptome and epigenetic landscapes correlate with elevated risk of cerebral microbleeds. Journal of the American Heart Association, 12(17):e031044, 2023.

[29] F. Müller, M. Scherer, Y. Assenov, P. Lutsik, J. Walter, T. Lengauer, and C. Bock. Rnbeads 2.0: comprehensive analysis of dna methylation data. Genome biology, 20(1):55, 2019.

[30] O. N. T. nanoporetech/modkit. Modkit, 2024.

[31] V. Nobile, C. Pucci, P. Chiurazzi, G. Neri, and E. Tabolacci. Dna methylation, mechanisms of fmr1 inactivation and therapeutic perspectives for fragile x syndrome. Biomolecules, 11(2):296, 2021.

[32] N. N. Olova and S. Andrews. Whole genome methylation sequencing via enzymatic conversion (em-seq): Protocol, data processing, and analysis. In High Throughput Gene Screening: Methods and Protocols, pages 73–98. Springer, 2024.

[33] PacificBiosciences. pb-cpg-tools. online, Jan. 2025.

[34] PacificBiosciences. Pacbio public sprq-nx multi-use chemistry hg002 download page. online, 2026.

[35] T. J. Peters, M. J. Buckley, A. L. Statham, R. Pidsley, K. Samaras, R. V Lord, S. J. Clark, and P. L. Molloy. De novo identification of differentially methylated regions in the human genome. Epigenetics &amp; Chromatin, 8(1), Jan. 2015.

[36] R. Pidsley, C. C. Y Wong, M. Volta, K. Lunnon, J. Mill, and L. C. Schalkwyk. A data-driven approach to preprocessing illumina 450k methylation array data. BMC genomics, 14(1):293, 2013.

[37] L. Poeta, D. Drongitis, L. Verrillo, and M. G. Miano. Dna hypermethylation and unstable repeat diseases: a paradigm of transcriptional silencing to decipher the basis of pathogenic mechanisms. Genes, 11(6):684, 2020.

[38] M. O. Pollard, D. Gurdasani, A. J. Mentzer, T. Porter, and M. S. Sandhu. Long reads: their purpose and place. Human molecular genetics, 27(R2):R234–R241, 2018.

[39] M. E. Ritchie, B. Phipson, D. Wu, Y. Hu, C. W. Law, W. Shi, and G. K. Smyth. limma powers differential expression analyses for rna-sequencing and microarray studies. Nucleic acids research, 43(7):e47–e47, 2015.

[40] B. D. Sigurpalsdottir, O. A. Stefansson, G. Holley, D. Beyter, F. Zink, M. P. Hardarson, S. P. Sverrisson, N. Kristinsdottir, D. N. Magnusdottir, O. P. Magnusson, D. F. Gudbjartsson, B. V. Halldorsson, and K. Stefansson. A comparison of methods for detecting dna methylation from long-read sequencing of human genomes. Genome Biology, 25(1), Mar. 2024.

[41] K. Skvortsova, C. Stirzaker, and P. Taberlay. The dna methylation landscape in cancer. Essays in biochemistry, 63(6):797–811, 2019.

[42] M. B. Stadler, R. Murr, L. Burger, R. Ivanek, F. Lienert, A. Scholer, E. v. Nimwegen, C. Wirbelauer, E. J. Oakeley, D. Gaidatzis, et al. Dna-binding factors shape the mouse methylome at distal regulatory regions. Nature, 480(7378):490–495, 2011.

[43] Y. Wang, E. Hannon, O. A. Grant, T. J. Gorrie-Stone, M. Kumari, J. Mill, X. Zhai, K. D. McDonald-Maier, and L. C. Schalkwyk. Dna methylation-based sex classifier to predict sex and identify sex chromosome aneuploidy. BMC genomics, 22(1):484, 2021.

[44] H. Wickham. ggplot2. Wiley interdisciplinary reviews: computational statistics, 3(2):180–185, 2011.

[45] C. O. Wilke and M. C. O. Wilke. Package ‘ggridges’. Ridgeline Plots in “ggplo*t2*, 2022.

[46] H. Wu, C. Wang, and Z. Wu. A new shrinkage estimator for dispersion improves differential expression detection in rna-seq data. Biostatistics, 14(2):232–243, 2012.

[47] M. J. Ziller, H. Gu, F. Müller, J. Donaghey, L. T.-Y. Tsai, O. Kohlbacher, P. L. De Jager, E. D. Rosen, D. A. Bennett, B. E. Bernstein, et al. Charting a dynamic dna methylation landscape of the human genome. Nature, 500(7463):477–481, 2013.

