## Supplementary Figures 1 to 15 for "MethylBench: A comprehensive benchmark of DNA methylation profiling methods across diverse sequencing platforms"

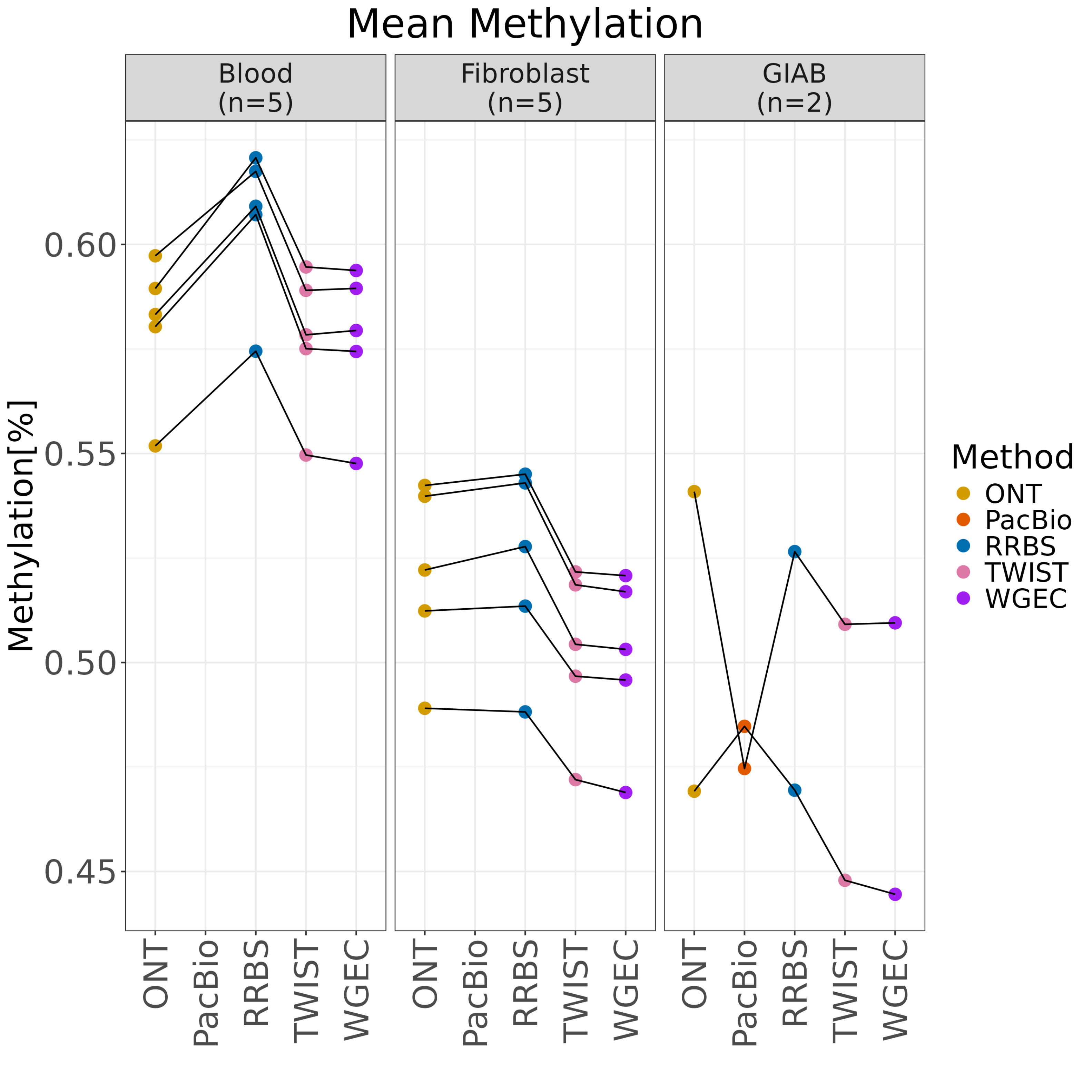
Mean Methylation
Blood (n=5)
Fibroblast (n=5)
GIAB (n=2)
Methylation[%]
0.60
0.55
0.50
0.45
ONT
PacBio
RRBS
TWIST
WGEC
Method
ONT
PacBio
RRBS
TWIST
WGEC

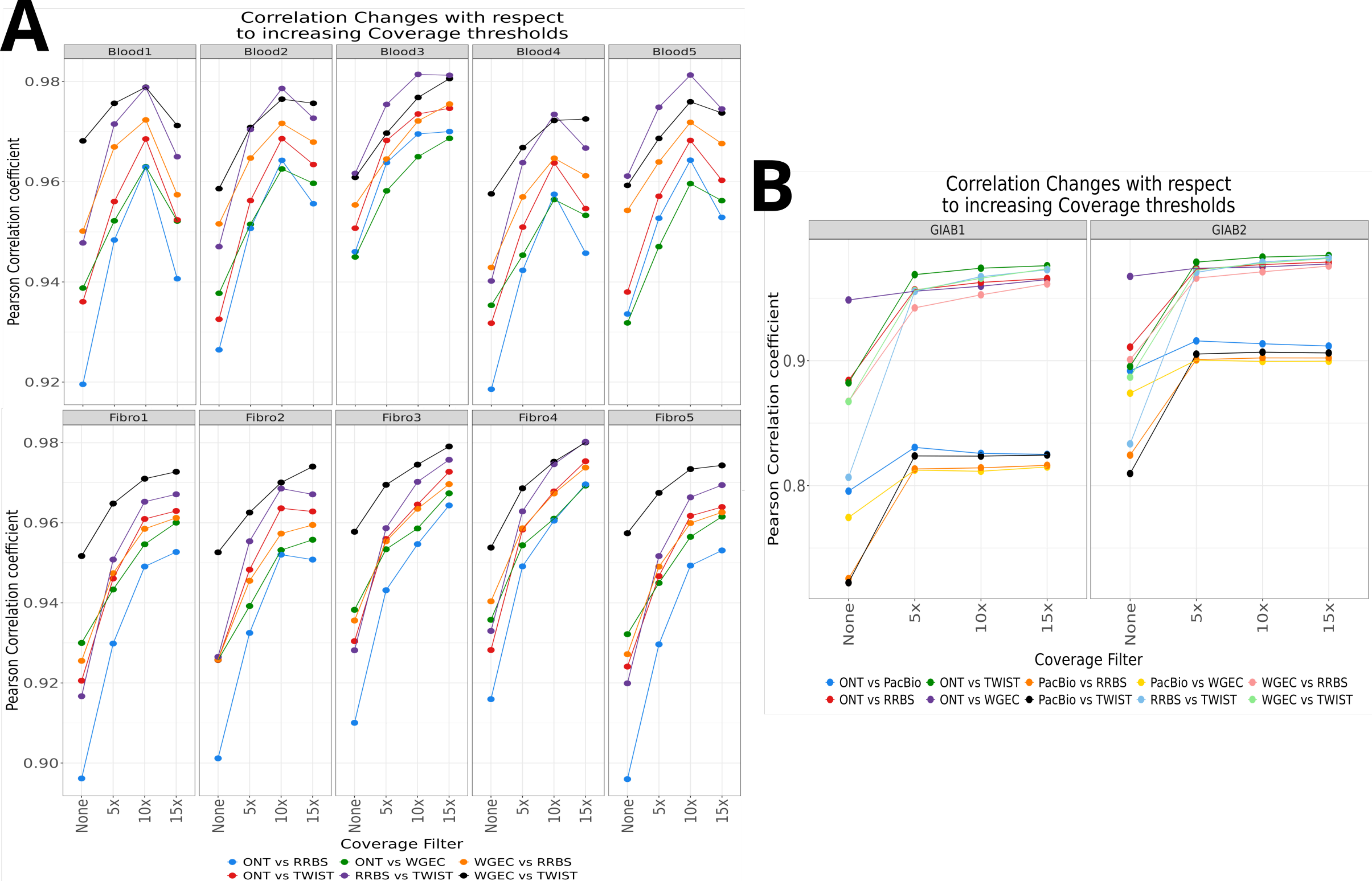
A
Correlation Changes with respect to increasing Coverage thresholds
Blood1
Blood2
Blood3
Blood4
Blood5
Fibro1
Fibro2
Fibro3
Fibro4
Fibro5
Pearson Correlation coefficient
0.98
0.96
0.94
0.92
0.90
None
5x
10x
15x
Coverage Filter
ONT vs RRBS
ONT vs TWIST
ONT vs WGEC
RRBS vs TWIST
WGEC vs RRBS
WGEC vs TWIST
B
Correlation Changes with respect to increasing Coverage thresholds
GIAB1
GIAB2
Pearson Correlation coefficient
0.9
0.8
None
5x
10x
15x
Coverage Filter
ONT vs PacBio
ONT vs RRBS
ONT vs TWIST
ONT vs WGEC
PacBio vs RRBS
PacBio vs TWIST
PacBio vs WGEC
RRBS vs TWIST
WGEC vs RRBS
WGEC vs TWIST

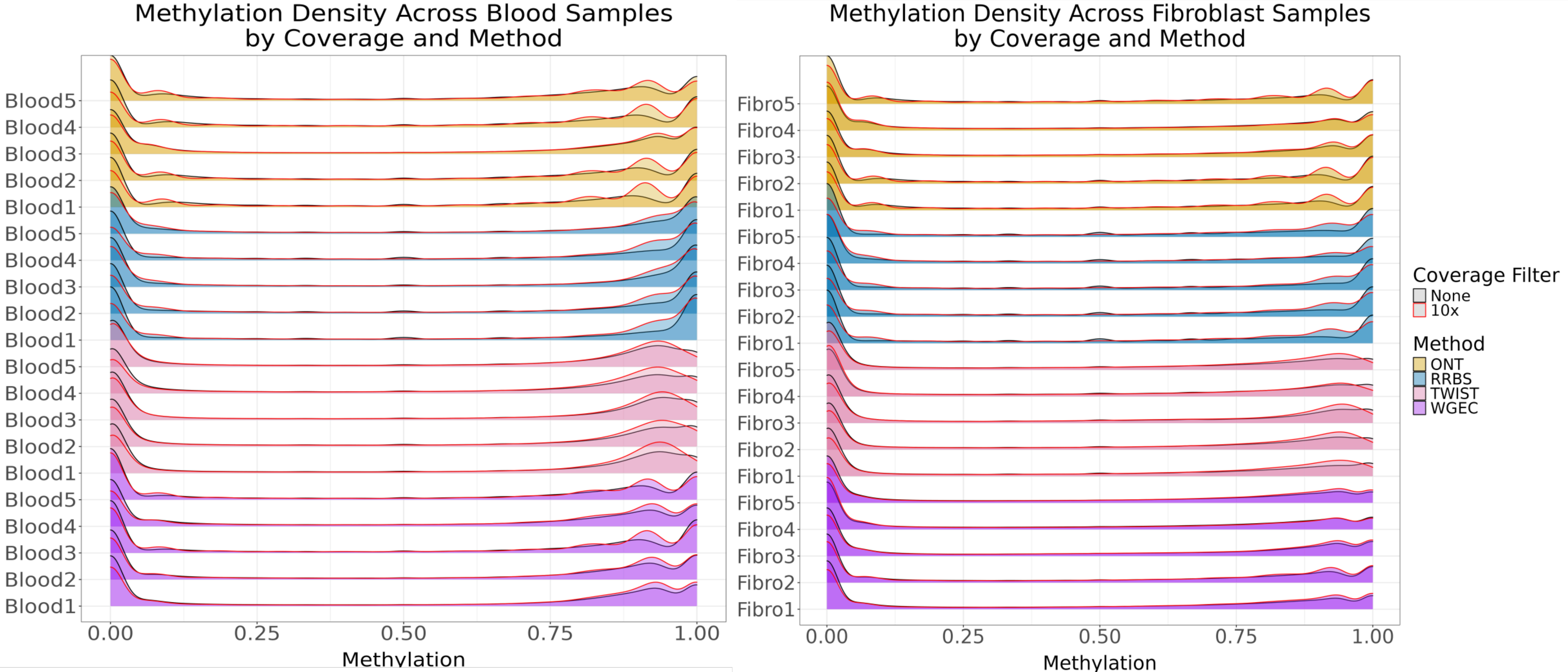

Methylation Density Across Blood Samples by Coverage and Method
Blood5
Blood4
Blood3
Blood2
Blood1
Blood5
Blood4
Blood3
Blood2
Blood1
Blood5
Blood4
Blood3
Blood2
Blood1
Blood5
Blood4
Blood3
Blood2
Blood1
0.00
0.25
0.50
0.75
1.00
Methylation
Methylation Density Across Fibroblast Samples by Coverage and Method
Fibro5
Fibro4
Fibro3
Fibro2
Fibro1
Fibro5
Fibro4
Fibro3
Fibro2
Fibro1
Fibro5
Fibro4
Fibro3
Fibro2
Fibro1
Fibro5
Fibro4
Fibro3
Fibro2
Fibro1
0.00
0.25
0.50
0.75
1.00
Methylation
Coverage Filter
None
10x
Method
ONT
RRBS
TWIST
WGEC

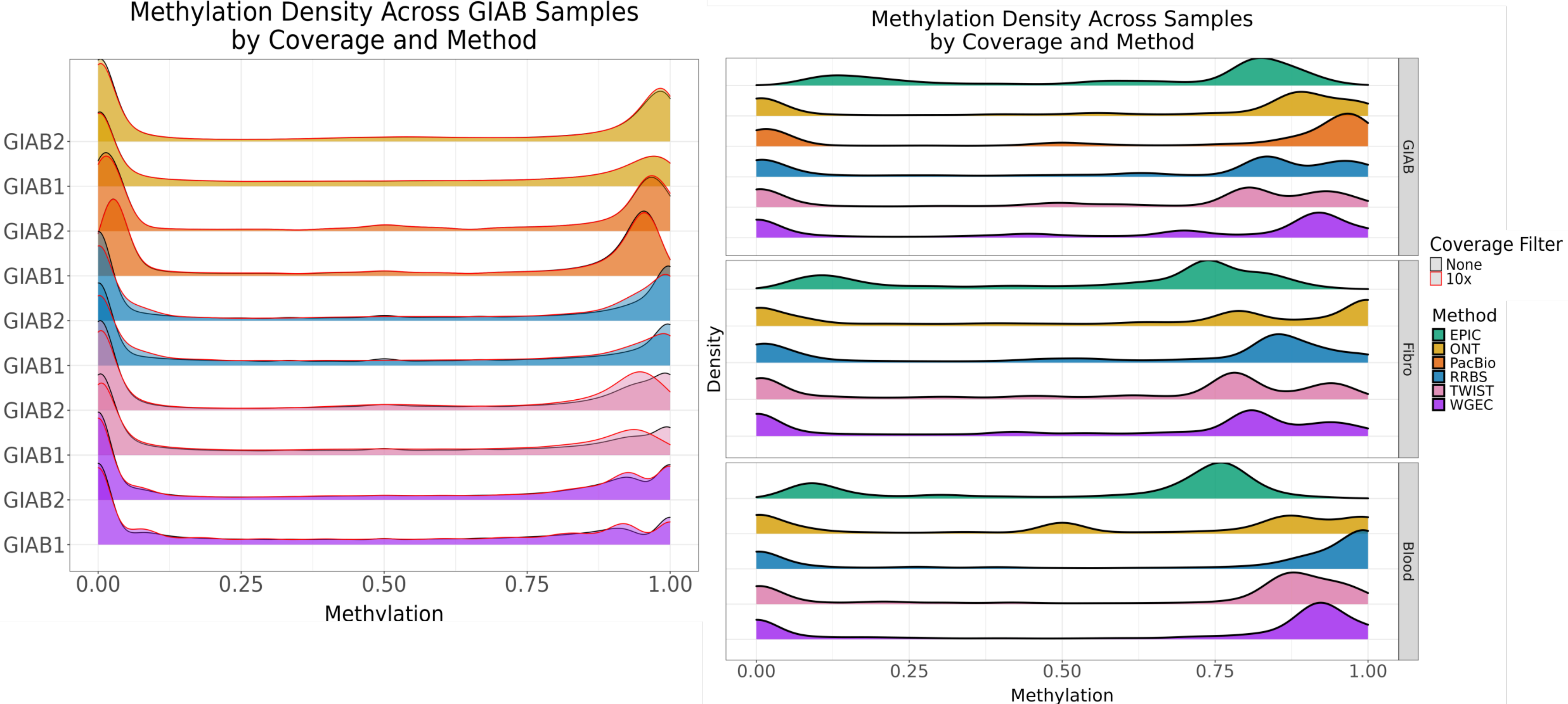

Methylation Density Across GIAB Samples
by Coverage and Method
GIAB2
GIAB1
GIAB2
GIAB1
GIAB2
GIAB1
GIAB2
GIAB1
GIAB2
GIAB1
0.00
0.25
0.50
0.75
1.00
Methylation
Methylation Density Across Samples
by Coverage and Method
Density
GIAB
Fibro
Blood
0.00
0.25
0.50
0.75
1.00
Methylation
Coverage Filter
None
10x
Method
EPIC
ONT
PacBio
RRBS
TWIST
WGEC

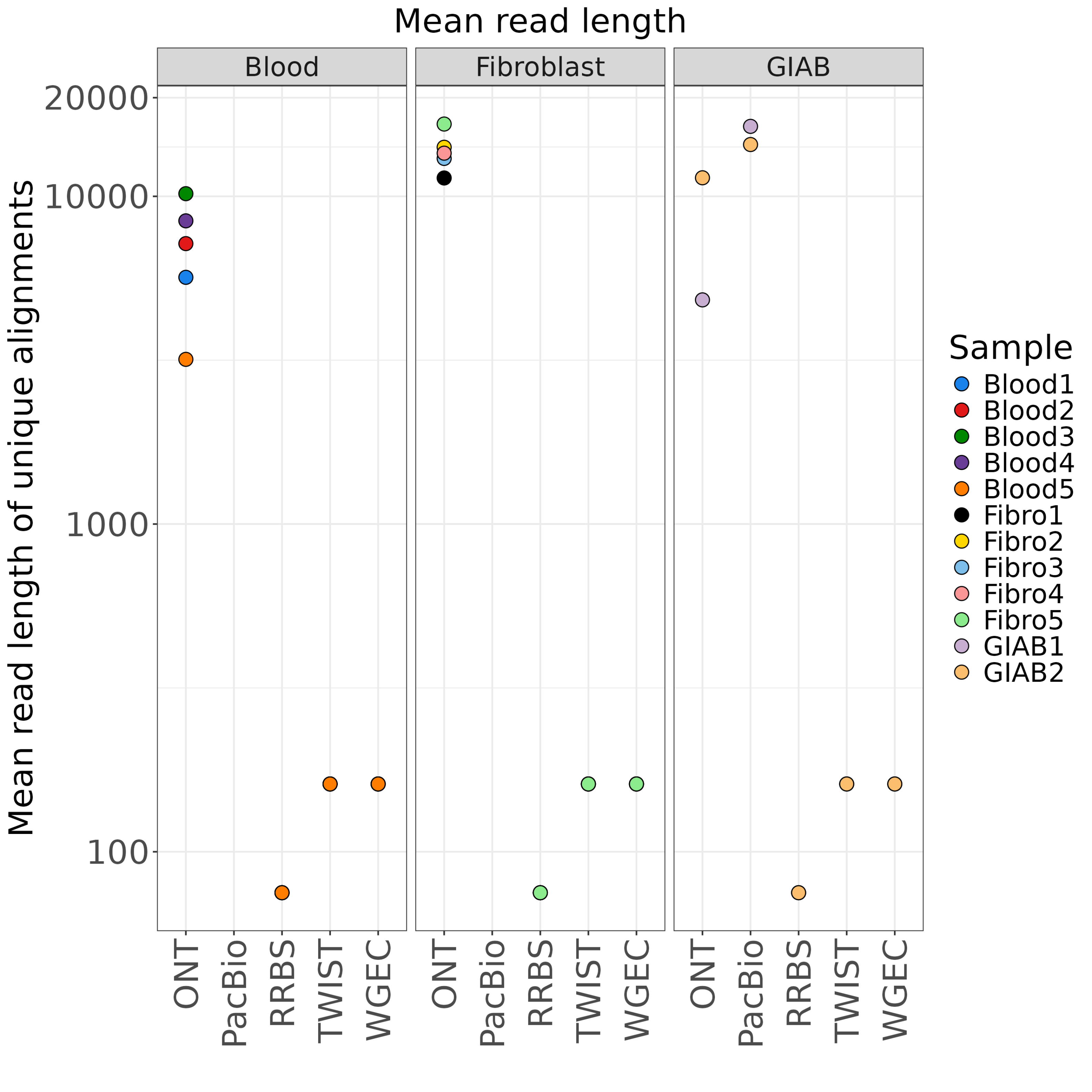

Mean read length
Blood
Fibroblast
GIAB
Mean read length of unique alignments
20000
10000
1000
100
ONT
PacBio
RRBS
TWIST
WGEC
Sample
Blood1
Blood2
Blood3
Blood4
Blood5
Fibro1
Fibro2
Fibro3
Fibro4
Fibro5
GIAB1
GIAB2

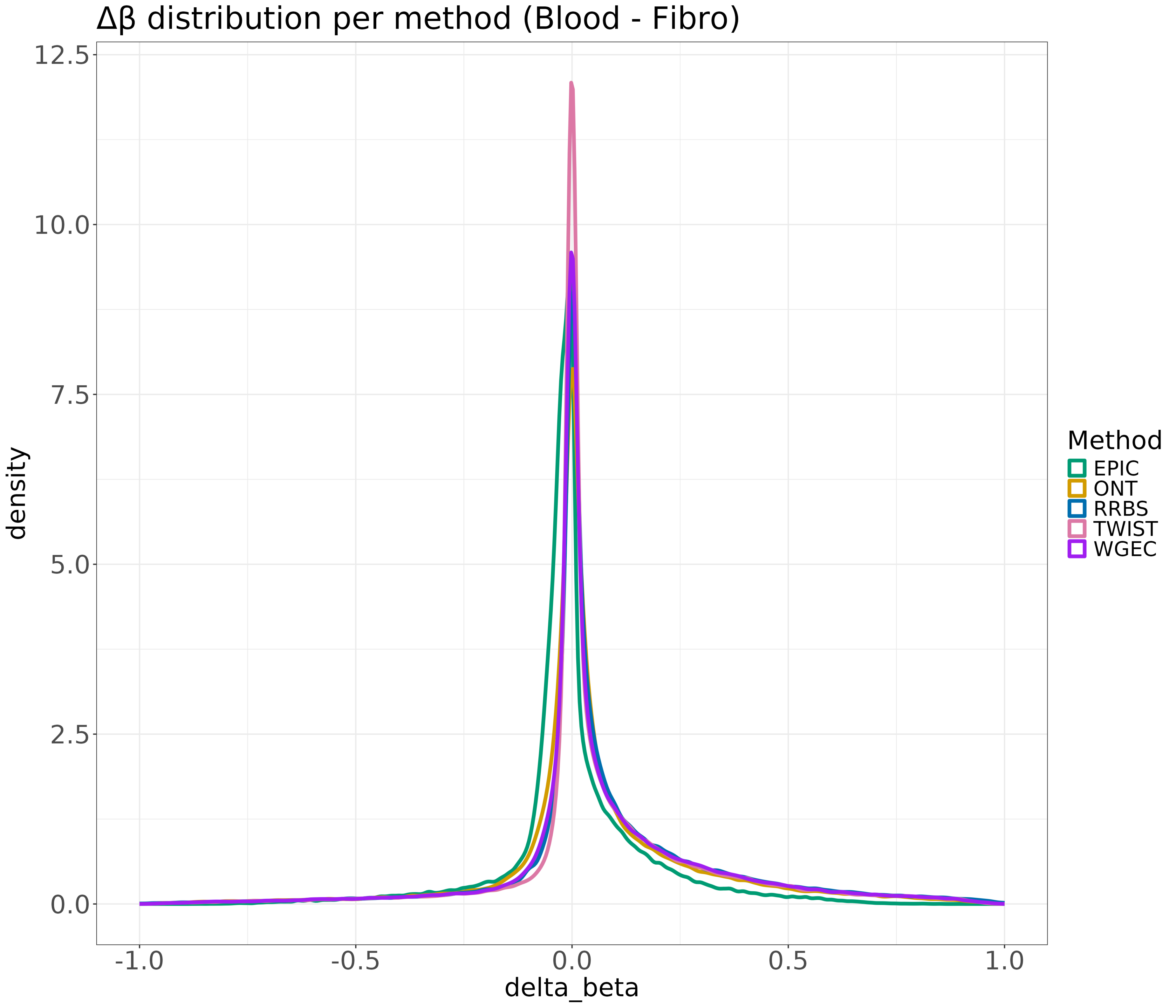
Δβ distribution per method (Blood - Fibro)
density
delta_beta
0.0
2.5
5.0
7.5
10.0
12.5
-1.0
-0.5
0.0
0.5
1.0
Method
EPIC
ONT
RRBS
TWIST
WGEC

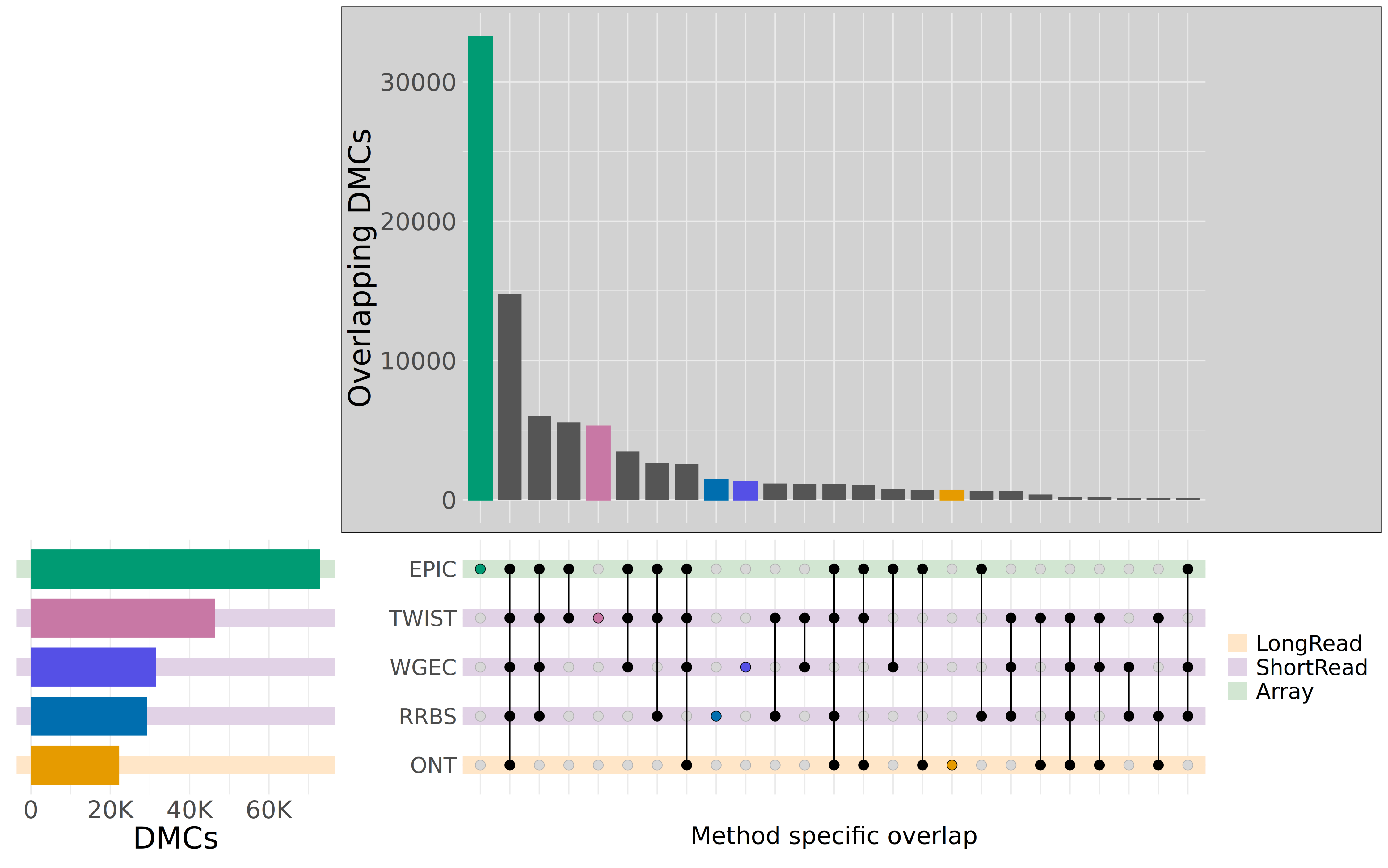

Overlapping DMCs
30000
20000
10000
0
EPIC
TWIST
WGEC
RRBS
ONT
0
20K
40K
60K
DMCs
Method specific overlap
LongRead
ShortRead
Array

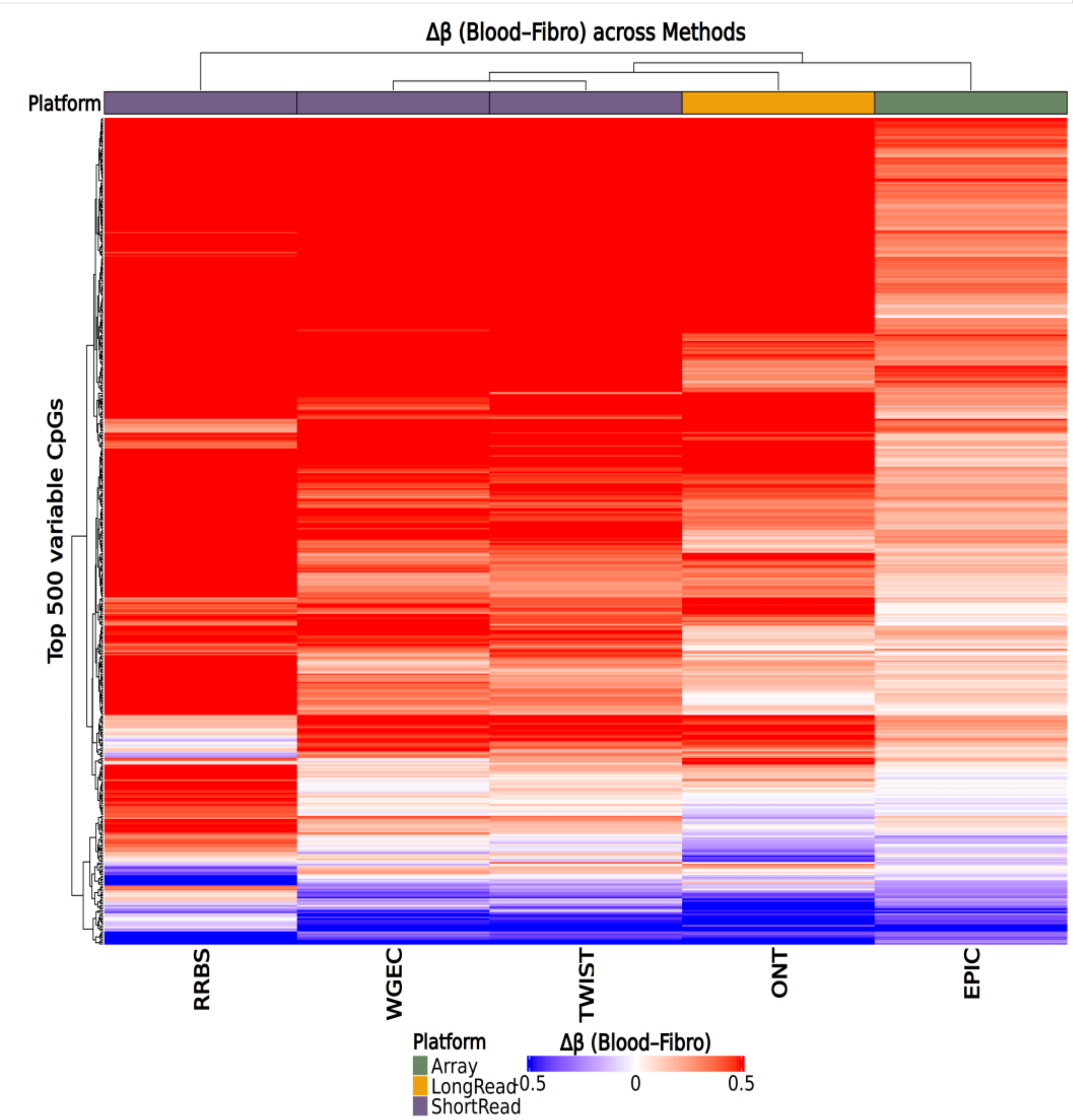
Δβ (Blood–Fibro) across Methods
Platform
Top 500 variable CpGs
RRBS
WGEC
TWIST
ONT
EPIC
Platform
Array
LongRead
ShortRead
Δβ (Blood–Fibro)
-0.5
0
0.5
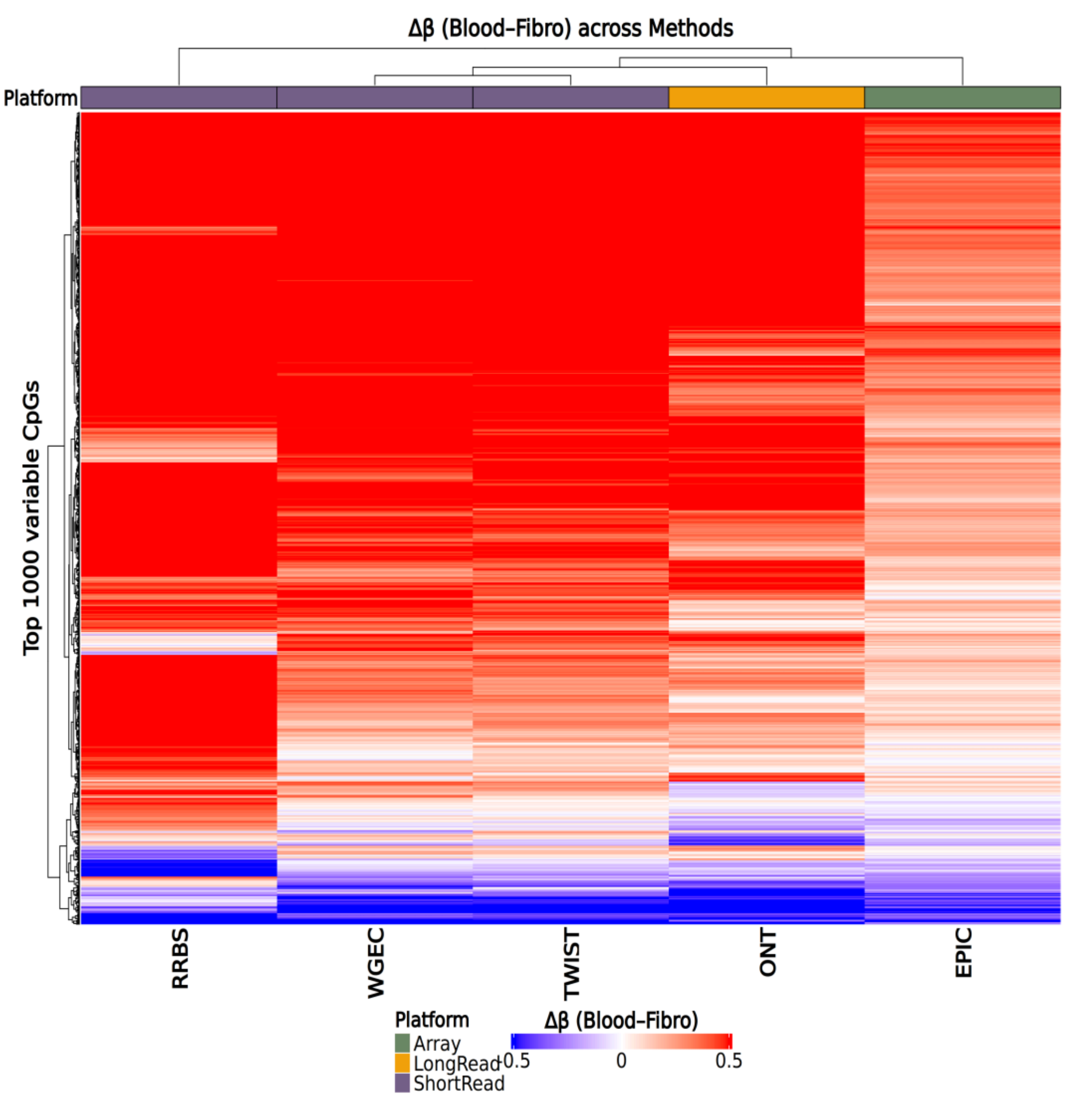
Δβ (Blood–Fibro) across Methods
Platform
Top 1000 variable CpGs
RRBS
WGEC
TWIST
ONT
EPIC
Platform
Array
LongRead
ShortRead
Δβ (Blood–Fibro)
-0.5
0
0.5

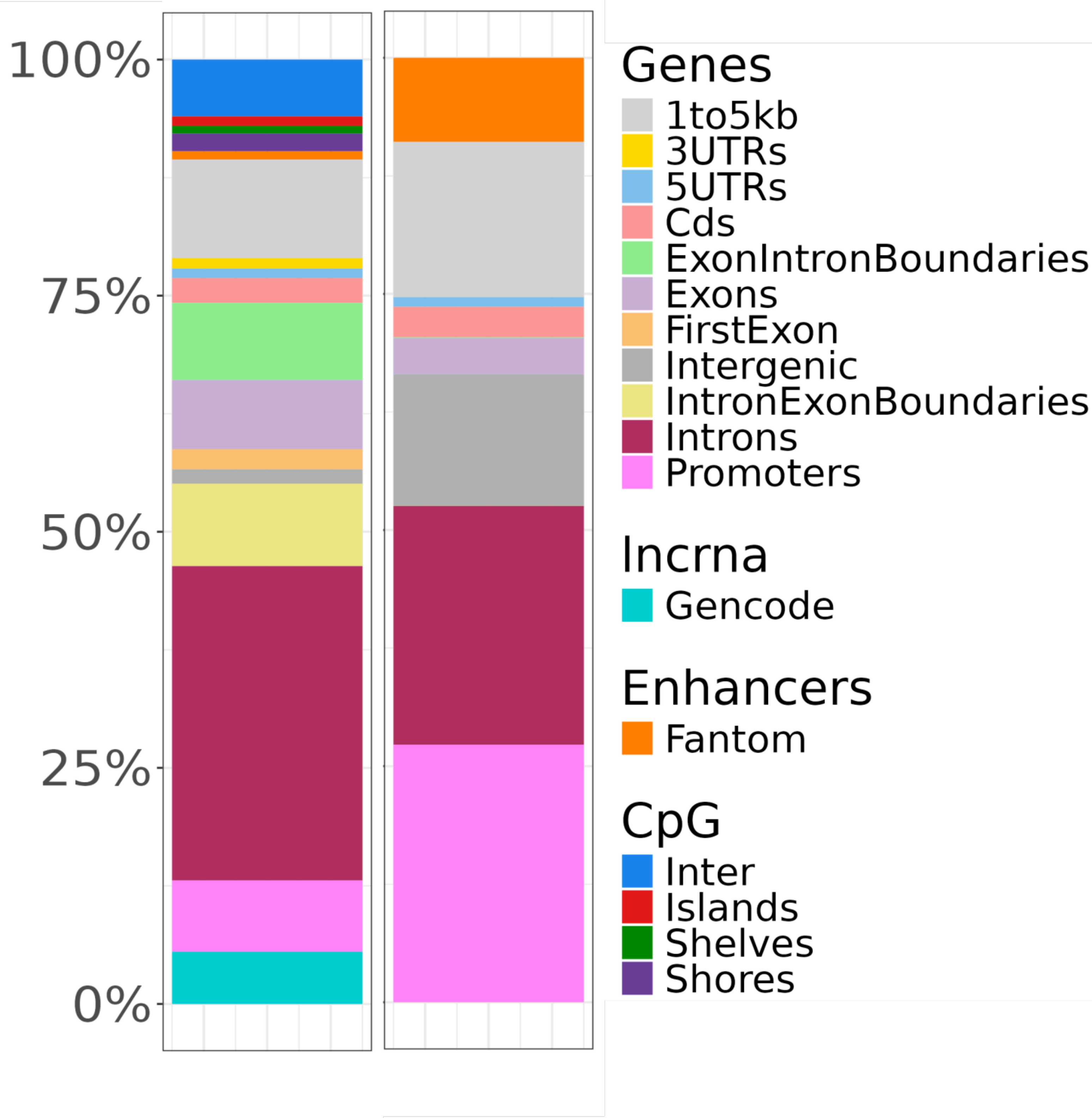
100%
75%
50%
25%
0%
Genes
1to5kb
3UTRs
5UTRs
Cds
ExonIntronBoundaries
Exons
FirstExon
Intergenic
IntronExonBoundaries
Introns
Promoters
Incrna
Gencode
Enhancers
Fantom
CpG
Inter
Islands
Shelves
Shores

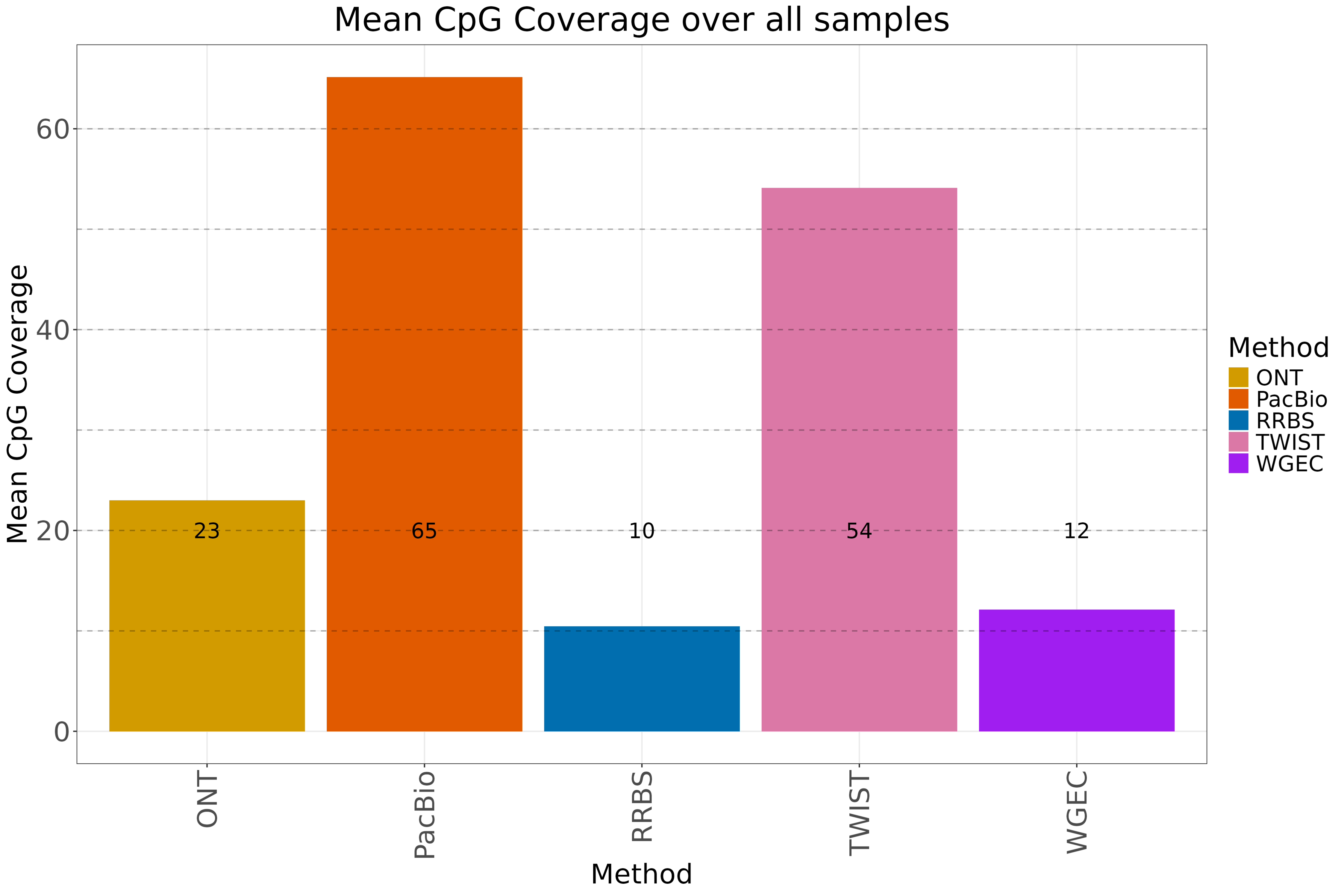

Mean CpG Coverage over all samples
Mean CpG Coverage
0
20
40
60
23
65
10
54
12
ONT
PacBio
RRBS
TWIST
WGEC
Method
Method
ONT
PacBio
RRBS
TWIST
WGEC

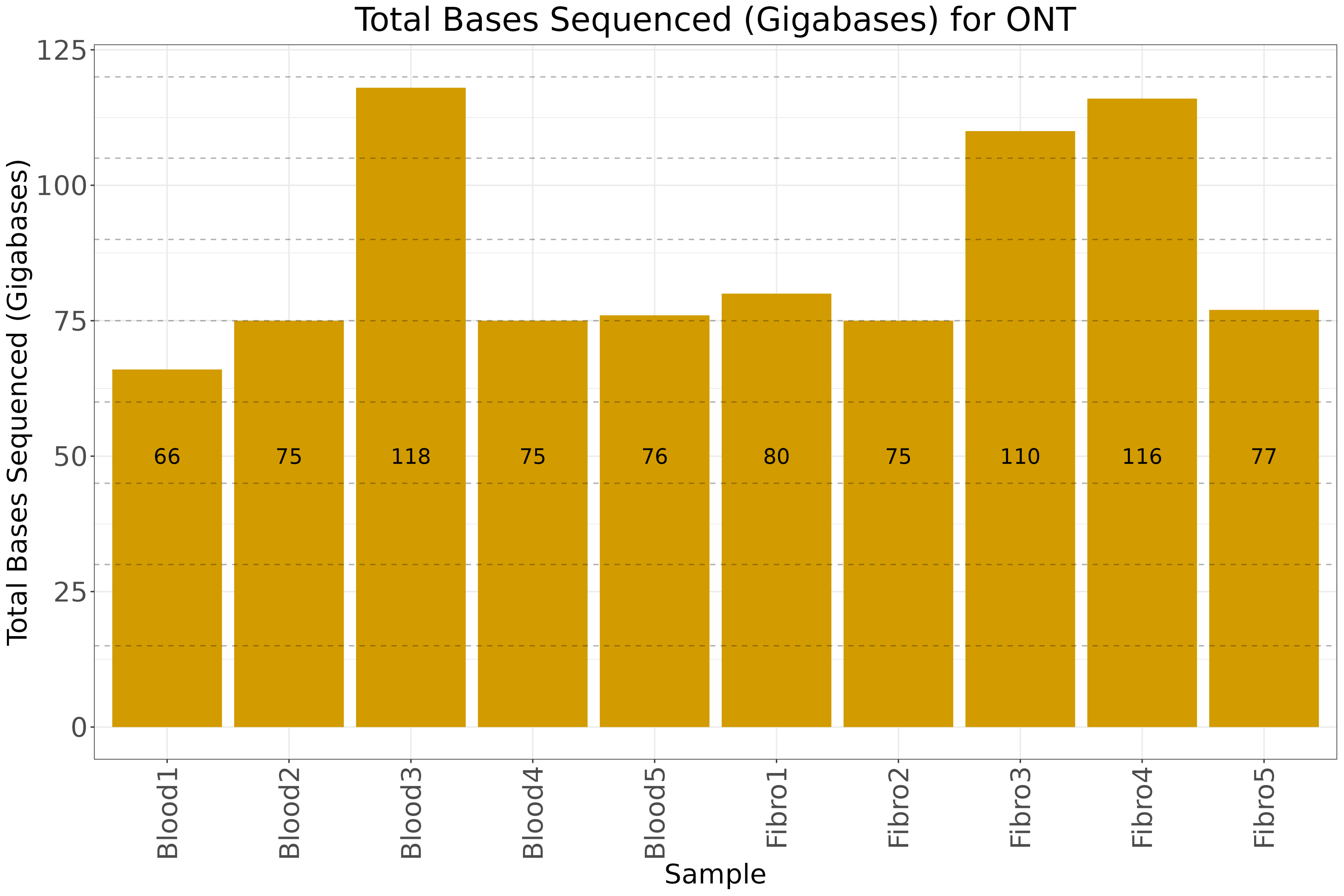
Total Bases Sequenced (Gigabases) for ONT
Total Bases Sequenced (Gigabases)
125
100
75
50
25
0
66
75
118
75
76
80
75
110
116
77
Blood1
Blood2
Blood3
Blood4
Blood5
Fibro1
Fibro2
Fibro3
Fibro4
Fibro5
Sample

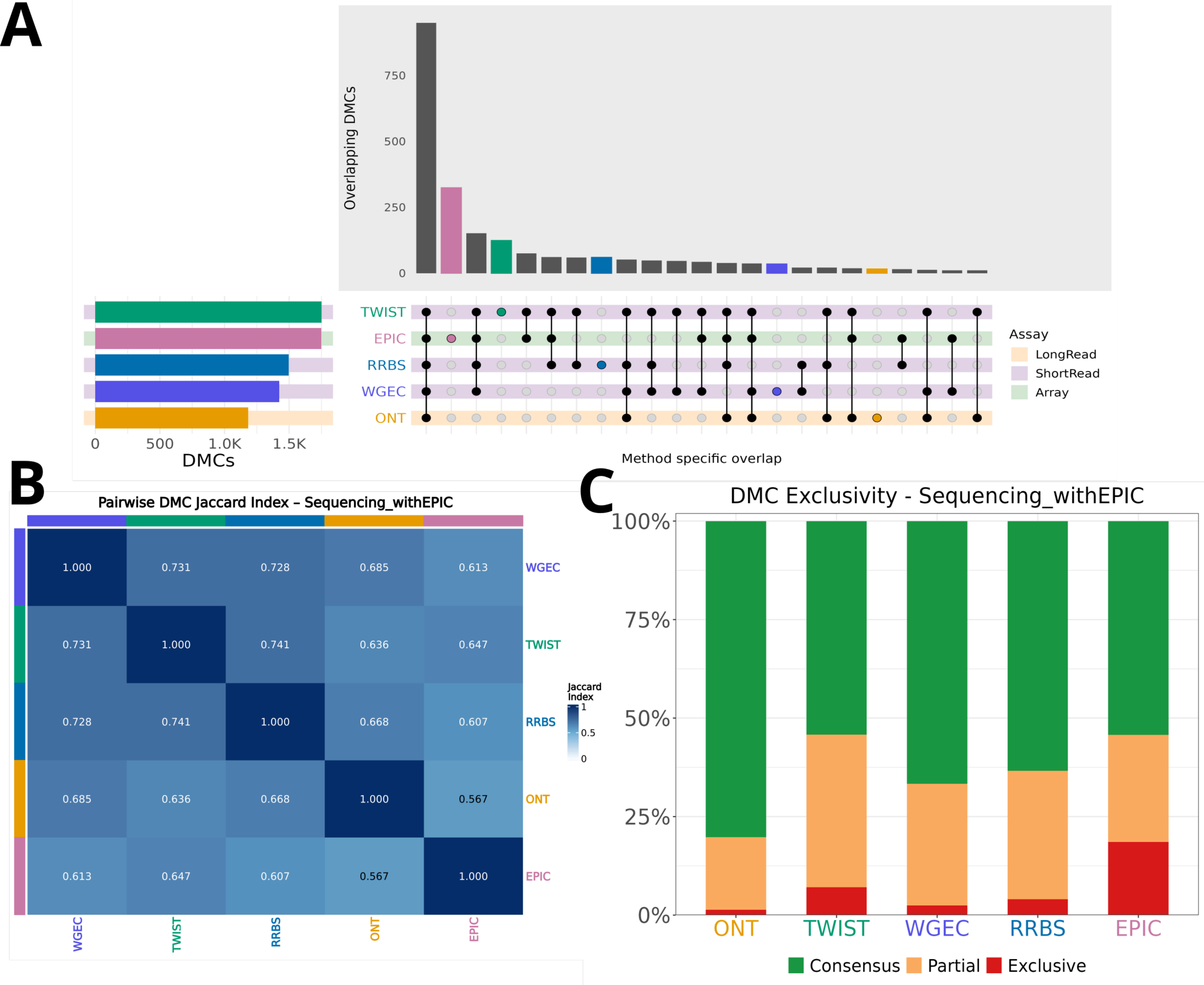

A
Overlapping DMCs
750
500
250
0
TWIST
EPIC
RRBS
WGEC
ONT
0
500
1.0K
1.5K
DMCs
Method specific overlap
Assay
LongRead
ShortRead
Array
B
Pairwise DMC Jaccard Index – Sequencing_withEPIC
WGEC
TWIST
RRBS
ONT
EPIC
Jaccard Index
1
0.5
0
1.000 0.731 0.728 0.685 0.613
0.731 1.000 0.741 0.636 0.647
0.728 0.741 1.000 0.668 0.607
0.685 0.636 0.668 1.000 0.567
0.613 0.647 0.607 0.567 1.000
C
DMC Exclusivity - Sequencing_withEPIC
100%
75%
50%
25%
0%
ONT
TWIST
WGEC
RRBS
EPIC
Consensus
Partial
Exclusive

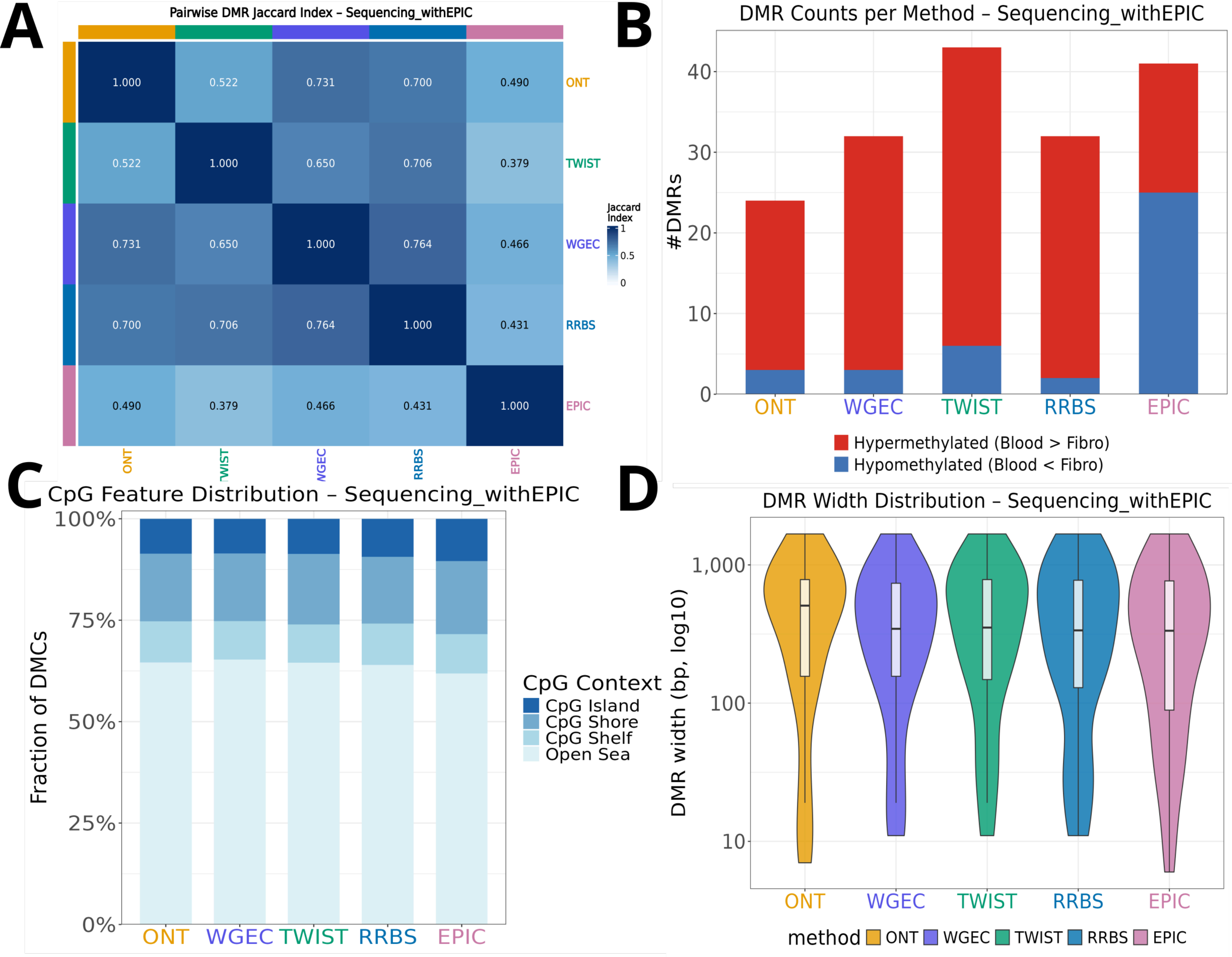

A
Pairwise DMR Jaccard Index – Sequencing_withEPIC
1.000 0.522 0.731 0.700 0.490 ONT
0.522 1.000 0.650 0.706 0.379 TWIST
0.731 0.650 1.000 0.764 0.466 WGEC
0.700 0.706 0.764 1.000 0.431 RRBS
0.490 0.379 0.466 0.431 1.000 EPIC
Jaccard Index
1
0.5
0
ONT TWIST WGEC RRBS EPIC
B
DMR Counts per Method – Sequencing_withEPIC
#DMRs
0 10 20 30 40
ONT WGEC TWIST RRBS EPIC
Hypermethylated (Blood > Fibro)
Hypomethylated (Blood < Fibro)
C
CpG Feature Distribution – Sequencing_withEPIC
Fraction of DMCs
0% 25% 50% 75% 100%
ONT WGEC TWIST RRBS EPIC
CpG Context
CpG Island
CpG Shore
CpG Shelf
Open Sea
D
DMR Width Distribution – Sequencing_withEPIC
DMR width (bp, log10)
10 100 1,000
ONT WGEC TWIST RRBS EPIC
method ONT WGEC TWIST RRBS EPIC

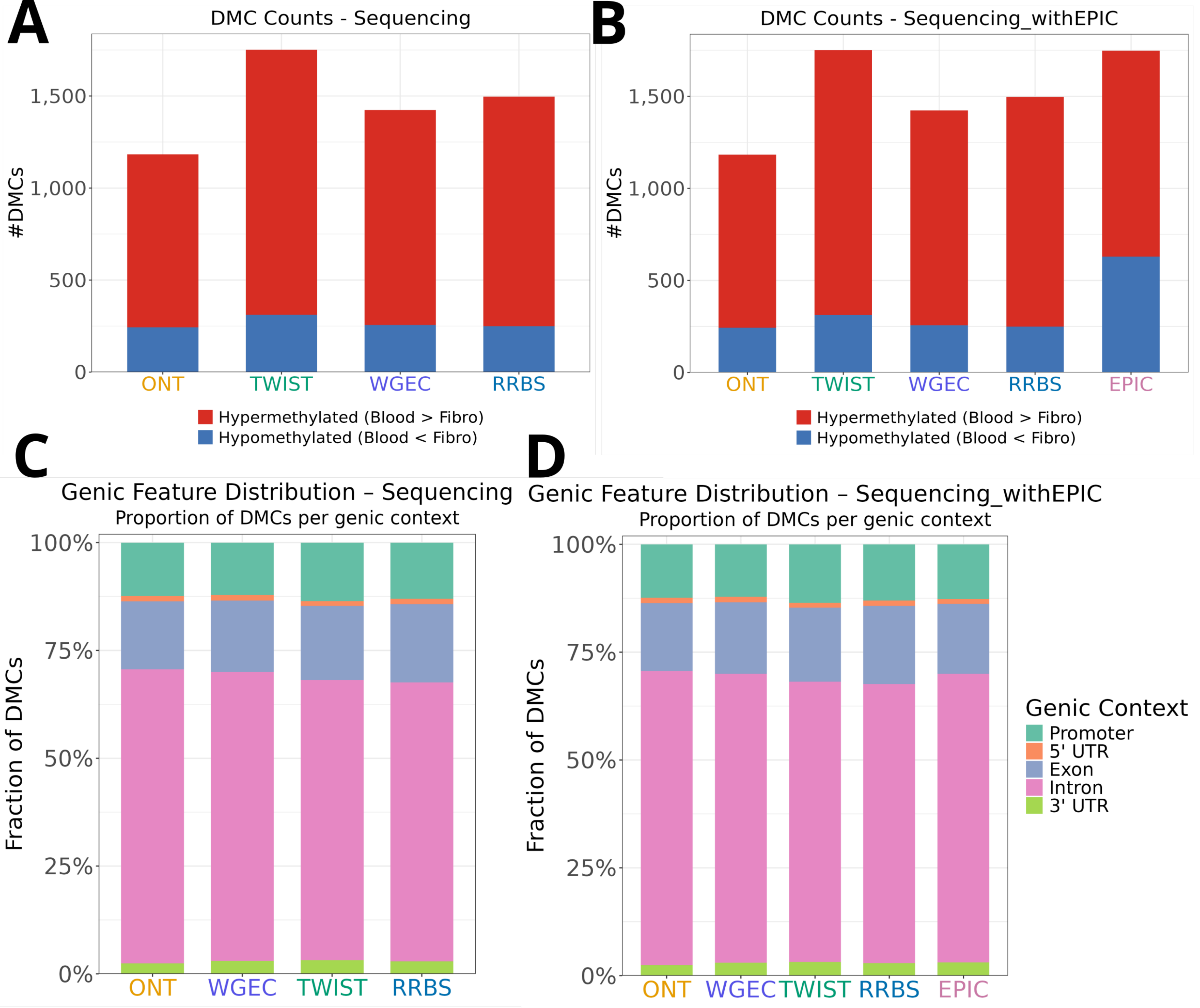

A
DMC Counts - Sequencing
#DMCs
1,500
1,000
500
0
ONT
TWIST
WGEC
RRBS
Hypermethylated (Blood > Fibro)
Hypomethylated (Blood < Fibro)
B
DMC Counts - Sequencing_withEPIC
#DMCs
ONT
TWIST
WGEC
RRBS
EPIC
Hypermethylated (Blood > Fibro)
Hypomethylated (Blood < Fibro)
C
Genic Feature Distribution – Sequencing
Proportion of DMCs per genic context
Fraction of DMCs
100%
75%
50%
25%
0%
ONT
WGEC
TWIST
RRBS
D
Genic Feature Distribution – Sequencing_withEPIC
Proportion of DMCs per genic context
Fraction of DMCs
ONT
WGEC
TWIST
RRBS
EPIC
Genic Context
Promoter
5' UTR
Exon
Intron
3' UTR

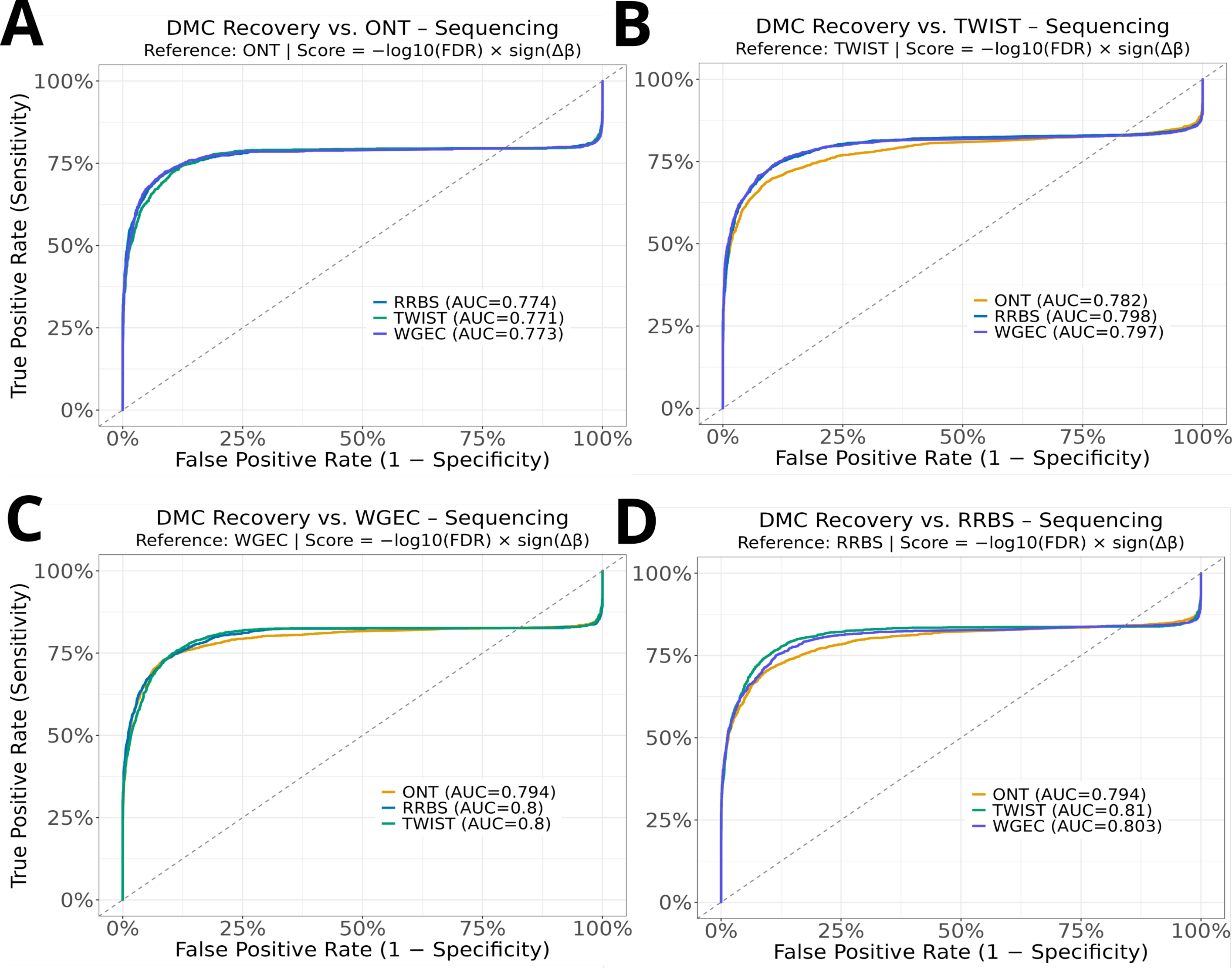
A
DMC Recovery vs. ONT – Sequencing
Reference: ONT | Score = −log10(FDR) × sign(Δβ)
True Positive Rate (Sensitivity)
100%
75%
50%
25%
0%
0%
25%
50%
75%
100%
False Positive Rate (1 − Specificity)
RRBS (AUC=0.774)
TWIST (AUC=0.771)
WGEC (AUC=0.773)
B
DMC Recovery vs. TWIST – Sequencing
Reference: TWIST | Score = −log10(FDR) × sign(Δβ)
ONT (AUC=0.782)
RRBS (AUC=0.798)
WGEC (AUC=0.797)
C
DMC Recovery vs. WGEC – Sequencing
Reference: WGEC | Score = −log10(FDR) × sign(Δβ)
ONT (AUC=0.794)
RRBS (AUC=0.8)
TWIST (AUC=0.8)
D
DMC Recovery vs. RRBS – Sequencing
Reference: RRBS | Score = −log10(FDR) × sign(Δβ)
ONT (AUC=0.794)
TWIST (AUC=0.81)
WGEC (AUC=0.803)

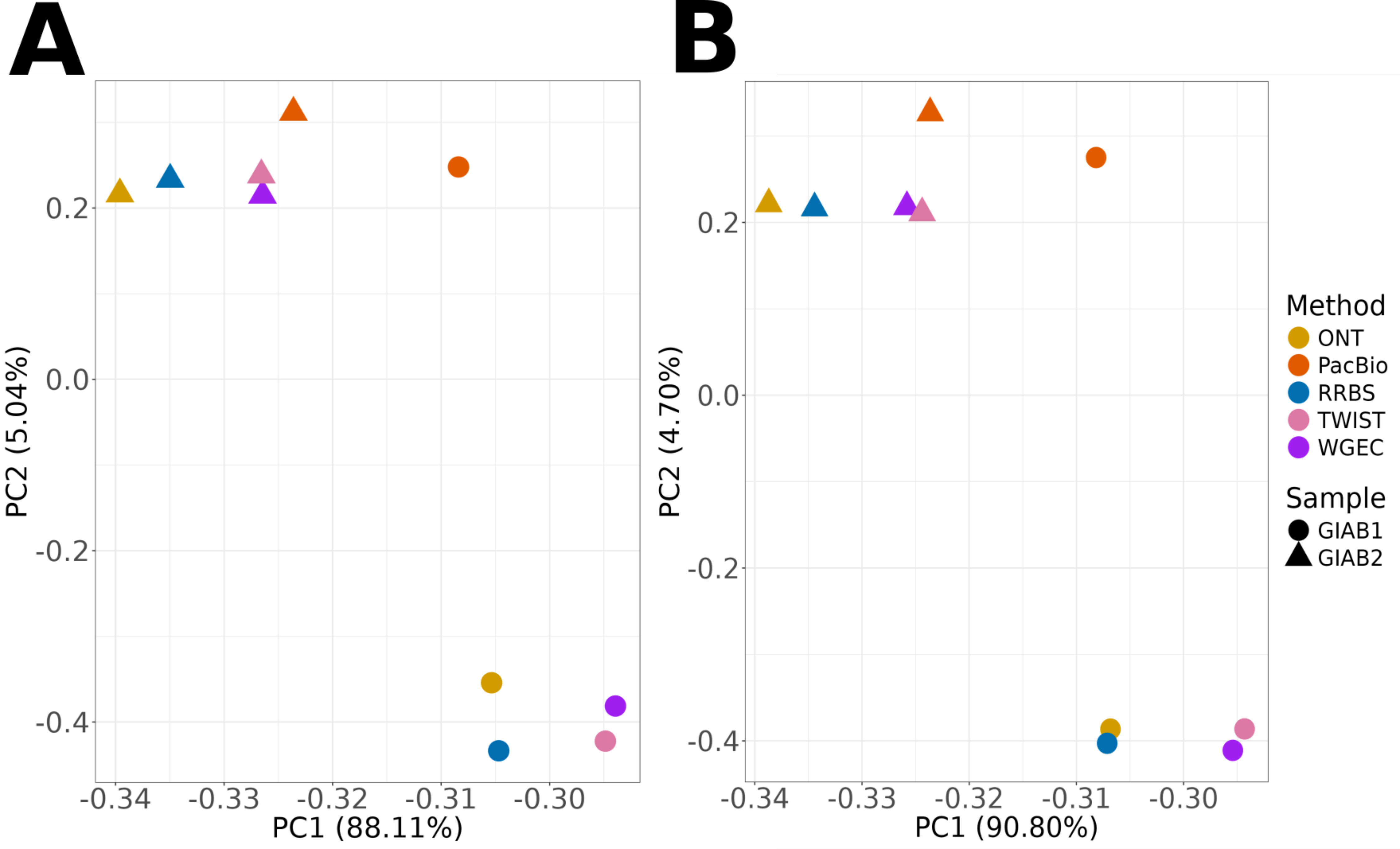
A
B
PC2 (5.04%)
PC1 (88.11%)
PC2 (4.70%)
PC1 (90.80%)
0.2
0.0
-0.2
-0.4
-0.34
-0.33
-0.32
-0.31
-0.30
Method
ONT
PacBio
RRBS
TWIST
WGEC
Sample
GIAB1
GIAB2
